# *Monod*: model-based discovery and integration through fitting stochastic transcriptional dynamics to single-cell sequencing data

**DOI:** 10.1101/2022.06.11.495771

**Authors:** Gennady Gorin, Tara Chari, Maria Carilli, John J. Vastola, Lior Pachter

## Abstract

Single-cell RNA sequencing analysis centers on illuminating cell diversity and understanding the transcriptional mechanisms underlying cellular function. These datasets are large, noisy, and complex. Current analyses prioritize noise removal and dimensionality reduction to tackle these challenges and extract biological insight. We propose an alternative, physical approach to leverage the stochasticity, size, and multimodal nature of these data to explicitly distinguish their biological and technical facets while revealing the underlying regulatory processes. With the Python package *Monod*, we demonstrate how nascent and mature RNA counts, present in most published datasets, can be meaningfully “integrated” under biophysical models of transcription. By utilizing variation in these modalities, we can identify transcriptional modulation not discernible though changes in average gene expression, quantitatively compare mechanistic hypotheses of gene regulation, analyze transcriptional data from different technologies within a common framework, and minimize the use of opaque or distortive normalization and transformation techniques.

## 1. Introduction

Technical advances in single-cell transcriptomics have enabled genome-wide molecular measurements in tens of thousands to millions of cells across diverse biological systems, from tissue samples and embryo models to whole organisms [1–5]. To address challenges related to the processing and interpretation of the associated high-dimensional and noisy datasets, single-cell genomics analyses have settled on an amalgamation of heuristic data normalization, transformation, and dimensionality reduction algorithms [6–8], which on the one hand extend bulk RNA-seq analyses, and on the other adapt large-scale machine learning approaches to remove presumed technical noise and extract biological signal. Although these approaches are frequently employed, they have increasingly recognized caveats related to hyperparameter sensitivity [9–11], distortion or loss of biological signal [12–15], incompatibility of qualitative methods with traditional statistical hypothesis testing approaches [9, 15, 16], and interpretability [16, 17] that may require more attention as the scale and diversity of single-cell datasets grow to include features like chromatin state and protein counts [13, 18].

A promising complementary approach, which respects the omnipresence of noise in the low copy-number regime [19–22], is to fit mechanistic (discrete, stochastic) models to single-cell data. These models are interpretable by construction, can be rigorously formulated in terms of generative models, are straightforwardly compatible with traditional statistical tools, and can reduce the necessity of hyperparameter tuning by making the relationships between biological assumptions and model parameters more explicit. Few large-scale fits of such models to single-cell data have been performed thus far, both due to technical issues only partially addressed by previous work [23–25], and non-technical issues related to descriptive modeling being more prevalent than mechanistic modeling in genomics [26]. Interestingly, this model-based approach is far more typical in the related field of fluorescence transcriptomics [19, 27–30].

Here we present the Python package *Monod*, which supports a model-based approach to single-cell genomics by automating biophysical-model-fitting and certain kinds of statistical testing (Figure 1a). In a typical scenario, the user defines a data likelihood by specifying both assumptions about the underlying biology, like whether transcription is bursty [27], and about the measurement process, like whether transcripts from longer genes tend to be captured more often [24]. *Monod* provides methods to efficiently fit this likelihood to large-scale data, and hence recover biophysically interpretable parameters along with estimates of their uncertainty; it also provides methods to run downstream analysis on these parameters, including comparing different biological hypotheses and testing for significant differences between experimental conditions.

**Figure 1:**
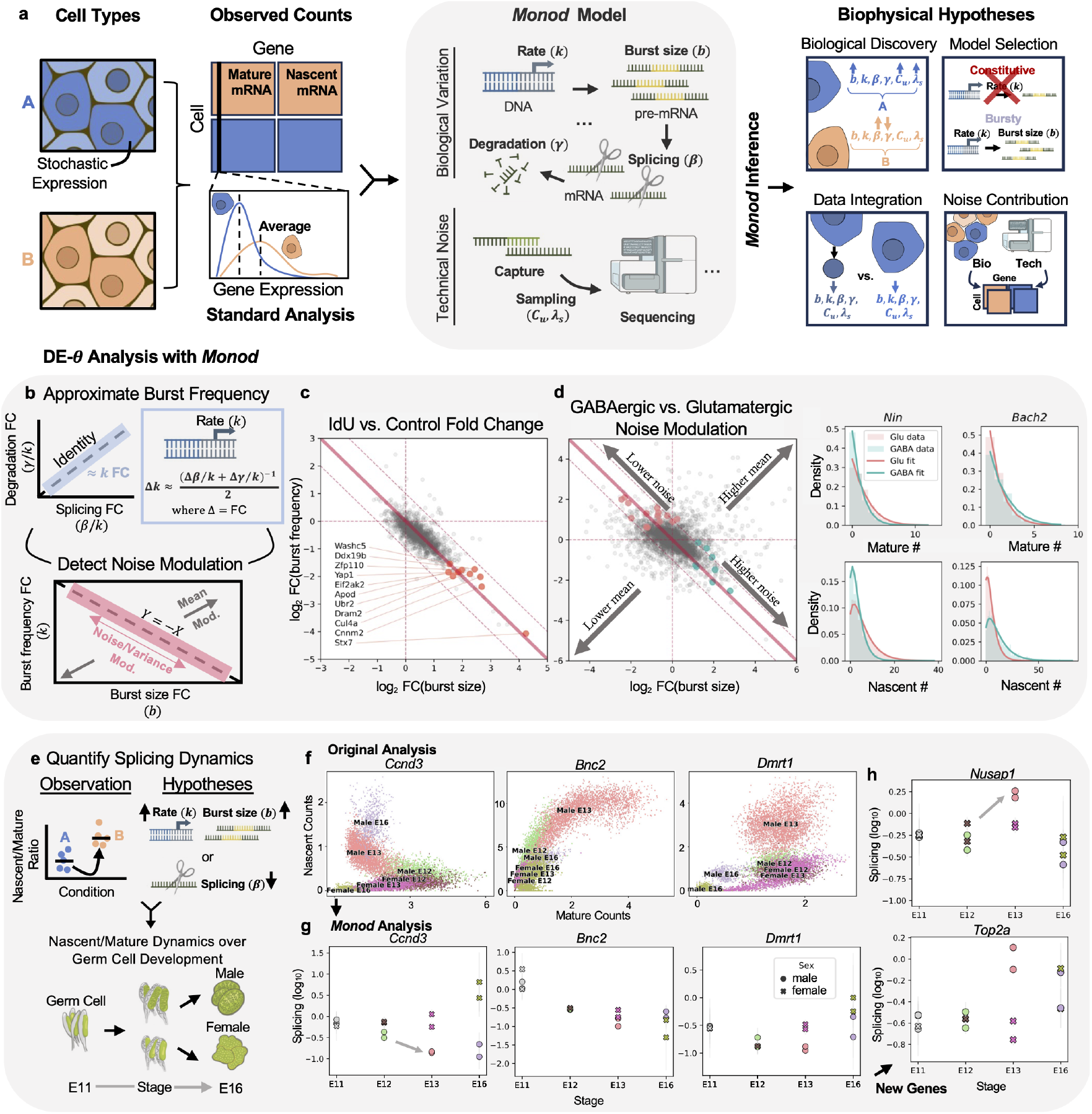
*Monod* inference for biophysical representation. **a**. Overview of the *Monod* framework for modeling multivariate genomics data. *Monod* performs inference of biophysical parameters from joint count distributions to develop mechanistic hypotheses. Example model of bursty transcription shown. Sequencer icon by DBCLS is licensed under CC-BY 4.0. **b**. (Top) How changes in burst frequency (*k*) are calculated, when splicing and degradation rate fold changes (FCs) are near identity. (Bottom) Changes in burst size and frequency denote noise versus mean expression modulation. **c**. Perturbation by IdU, which triggers DNA damage and repair, rarely changes expression levels, but induces genome-wide noise enhancement [54] detectable by *Monod* (lines and gray points: as in **a**; red points and labels: well-fit, moderate-expression genes identified as highly noise-enhanced). **d**. (Left) Differences between mouse glutamatergic and GABAergic cell types, n=4 replicates, include genes with substantial noise but little to no change in average expression (light red points: significantly higher noise in glutamatergic cells; light teal points: significantly higher noise in GABAergic cells; gray points: all other genes; solid diagonal line: parameters where burst size and frequency compensate to maintain a constant average expression; dashed diagonal lines: *±*1 log_2_ expression FC region about the constant-average expression line; vertical and horizontal lines: parameter combinations where burst size and frequency, respectively, do not change). (Right) Cell type-specific distribution shapes (light red: glutamatergic cell type; light teal: GABAergic cell type; histograms: raw counts; lines: *Monod* fits; top row: mature RNA marginal; bottom row: nascent RNA marginal). **e**. How changes in ratios of nascent and mature mRNA counts can derive from different mechanistic hypotheses. Analyses with *Monod* performed on developmental dataset of mouse germ cells (male and female). Figure adapted from [57] (CC BY-NC 4.0). **f**. Nascent and mature counts of key genes in the original study [57]), over the embryo stages and sexes. Figure adapted from [57] (CC BY-NC 4.0). **g**. *Monod* splicing rate fits for these key genes for each of the n=2 biological replicates. Error bars: 99% confidence regions determined by *±*2.576 *σ*. Grey arrows denote examples of changing splicing rates. **h**. Trends for new candidate genes where splicing rate increases over the stages in testes germ cells. Grey arrows denote examples of changing splicing rates.

The model-based approach to single-cell genomics does not replace the standard approach, but rather extends it in some ways and complements it in others. It extends it by improving rigor and suggesting a richer set of biologically interesting hypotheses (i.e., models beyond Poisson and negative binomial) to compare to data, and complements it by supplying natural alternative quantities on which to run statistical comparisons and perform clustering. The purpose of *Monod* is twofold: to leverage prior biophysical knowledge to extract more and different information from the same dataset, and to provide a rigorous computational framework for such analyses, which can be extended far beyond the use cases we consider here (see Chari et al. [31] for one such extension).

In the following, we illustrate the capabilities of *Monod* through a comprehensive description of its modeling principles and a series of vignettes, including analyses of 9 single-cell datasets which span diverse application domains (cell culture, cancer, radiation treatment, development, blood, and brain) and two species (mouse and human). We show *Monod* can utilize nascent and mature mRNA count data to generate quantitative hypotheses about DNA and RNA regulation in heterogeneous cell populations, identify or assess model misspecification, integrate biology across sequencing platforms and techniques, and limit data distortion (Figure 1a). Thus, as we continue to improve the depth and diversity of single-cell, molecular measurements, *Monod* provides a principled statistical foundation to represent relevant biological phenomena and questions about mechanisms of transcriptional variability, and to falsify these models and hypotheses based on the data. Further details about the *Monod* workflow are defined below and in Section 9.1, with a visual representation and example outputs from the workflow shown in Supplementary Figure 1.

## 2. Results

### 2.1 Assumptions and approach of *Monod*

*Monod* takes as input measurements of nascent and mature RNA count data, which can be obtained by aligning single-cell RNA-seq (scRNA-seq) reads to intron-and exon-containing references [32,33], and fits biophysical models of transcription. An advantage of the model-based approach is that it allows one to incorporate prior knowledge into analyses, like the fact that the production of nascent and mature RNA are causally linked. Common bioinformatic analysis pipelines often treat nascent and mature RNA as independent rather than causally linked, conflate them, or ignore nascent mRNA entirely [13, 33].

All *Monod* models have two components: a biological component, which implements prior knowledge about the underlying biophysics, and a technical component, which implements prior knowledge about how the measurement process tends to distort the underlying biology. *Monod* supports a variety of model types, but we will focus on models with the following structure:

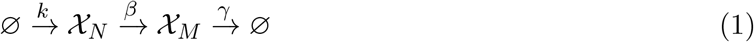

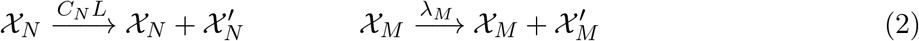

The underlying biological assumptions are that nascent RNA *𝒳*_*N*_ is produced (with *k* the transcription rate), converted into mature RNA *𝒳*_*M*_ (with *β* the splicing rate), and degrades (with *γ* the degradation rate). The underlying technical assumption is that a given mRNA molecule may be captured and reverse transcribed into cDNA multiple times, with the capture rate *C*_*N*_ *L* of nascent RNA proportional to the gene length *L* and the capture rate of mature RNA independent of gene length [24, 25]. *Monod* assumes that input data is approximately at steady state, so it fits all rate parameters in units of *k*, which is equivalent to imposing *k* = 1.

*Monod* supports a variety of biophysically plausible transcription models, each of which make slightly different assumptions about transcription, splicing, and degradation. This flexibility allows users to compare how well different biological hypotheses explain a given dataset; in general, one does not expect any single model to always perform best.

The simplest assumption one can make is that the probability per unit time of each process occurring is constant, or equivalently that the waiting time associated with each process is exponentially distributed. If one assumes this for each process, one obtains the *constitutive model*, which yields Poisson-distributed RNA count distributions [34, 35] that do not exhibit the overdispersion commonly observed in real RNA count datasets [27, 36]. There are two well-known ways to complexify the constitutive model to incorporate overdispersion: the first is to assume that RNA is produced in “bursts”, with the burst size *B* assumed to be geometrically distributed [27, 37–43] (the *bursty model* ); the second is to assume that the transcription rate *k* is gamma-distributed due to cell-cell heterogeneity [23, 44] (the *extrinsic model* ). Each approach only modifies the first arrow in model 1 of the above schematic.

*Monod* currently supports 6 models of transcription, including the constitutive, bursty, and extrinsic models. The quasi-bursty limit of the Cox–Ingersoll–Ross model [23], and models which replace the exponentially-distributed waiting times of either degradation or splicing with deterministic waiting times [45, 46], are also supported. *We recommend the bursty model as the default choice for most analyses*, as it is consistent with known biophysics and empirically fits many single-cell datasets well. See Section 2.4 for more discussion of each model type and how they compare in practice, and see Section S1 for the detailed definitions and properties of different models of biological and technical variability.

The inference process involves estimating two types of model parameters: biological parameters like *β* and *γ*, and the technical parameters *C*_*N*_ and *λ*_*M*_ . Because the technical parameters are only weakly identifiable [24], *Monod* assumes they are shared across genes (but not necessarily across cell types). *Monod* estimates the technical parameters through a grid search, and conditional on a given set of technical parameters, estimates biological parameters via gradient descent on the Kullback-Leibler divergence between the empirical and fitted distributions. Once a set of parameters that minimize the aforementioned divergence are identified, *Monod* quantifies parameter estimate uncertainty by exploiting the Fisher information matrix of the data–model divergence. *Monod* also supports *post hoc* tests of goodness-of-fit, which it accomplishes by a combination of criteria based on chi-squared statistics, Hellinger distances from the theoretical distribution, and adequate distance of parameters from user-imposed bounds (see example rejected genes in Supplementary Figure 3). For more on the inference process, see Section 9.1 and previous work [24].

Given an assumed model, *Monod* ‘s output is a set of model parameter estimates, parameter uncertainties, and goodness-of-fit metrics (see example outputs in Supplementary Figures 1 and 3). In the following vignettes, we discuss some mechanistic questions which can be interrogated using *Monod* fits, essentially by comparing parameter fits between biological samples or models.

### 2.2 *Monod* generalizes differential expression analyses

The goal of single-cell genomics studies is often to perform exploratory analysis of complex biological systems to develop new questions and hypotheses about how different cell states and types are defined and result in heterogeneous functions. Common approaches use distance metrics or averages of gene expression across cells to identify cell clusters and “neighbors,” and compare expression profiles across conditions and continuous processes. Using biophysical inference, we can generalize this assessment of cell type heterogeneity beyond a descriptive, data-scientific comparison of summary statistics and representation of cells as points in average gene expression space. The representation of cells through their biophysical characteristics provides a more general definition of cell “type,” simultaneously summarizing the molecule distributions and more closely encoding the intuition that cells comprise, regulate, and are exposed to stochastically varying environments [47, 48].

In typical transcriptomics workflows, determination of differences between cell types or states relies on identification of differentially expressed (DE) genes [49–51], which exhibit statistically significant differences in their average gene counts. However, these differences can be defined by both changes in mean and variance of gene expression, and regulatory mechanisms such as modulation of transcriptional noise [47,48] can maintain constant mean and would not be identifiable by standard statistical methods. The identification of differential expression is also impacted by treatment of technical covariates [6]. Thus, the mechanistic *Monod* framework generalizes the notion of differential behavior to modulated parameters, which describe not only how the distribution of gene counts change, but also which underlying biological processes effect this change.

Here we demonstrate how differential parameter behavior captures alterations to mean and variance of gene expression counts, as well as quantifies changes in specific biological processes such as pre-mRNA splicing, that would otherwise rely on more qualitative analyses. This extension of DE to biophysical parameters is denoted as “DE-*θ*,”, where *θ* may be a data moment or an inferred biophysical parameter (Figure 1b).

Modulation of transcriptional noise, which may or may not result in changes to average gene expression, has been shown to affect cell fate determination [52] and DNA damage recovery among other processes [47], leading to growing interest in identifying noise regulators in the cell [53]. To demonstrate the ability of *Monod* DE-*θ* to detect modulations without substantial changes in average expression, we first applied *Monod* to data from a recent study [47] which found that the introduction of a modified nucleotide (IdU) to a culture medium enhances transcriptional noise, but keeps average expression constant, hinting at a genome-wide mechanism for compensation [47, 54] (Figure 1b,c). The inferred parameters, for the bursty model with length-biased capture, demonstrated striking noise amplification in IdU-perturbed cells, with limited differences in mean expression (Figure 1c). This includes genes which generally mediate the cellular stress response: *Zfp110, Eif2ak2*, and *Yap1* regulate apoptosis, as well as *Dram2*, involved in the autophagic response to DNA damage repair, *Cul4a*, involved in the turnover of DNA repair proteins, and *Ddx19b*, potentially active in stress granules (Figure 1c). These findings are consistent with the authors’ conclusions and independent reanalysis, which likewise found that burst size increases and burst frequency decreases in the IdU condition [47,54,55]. The ascribed change in burst frequency reflects the change in inferred splicing and degradation rates being near identity, as it is more likely that such coordinated changes in unrelated processes are a result of changes to the denominator (*k*) (see Methods Section 9.2.2).

We also find that noise modulation is present not just in this more artificial perturbation context, but also in heterogeneous cell types, such as between glutamatergic and GABAergic neurons in the mouse brain [56] (Figure 1d). For example, *Nin* and *Bach2*, which are involved in neuronal development, visually exhibit higher noise in the glutamatergic and GABAergic populations, respectively (Figure 1d). The mature count averages are, on the other hand, fairly close (*Nin* Glu: 1.7, GABA: 0.98; *Bach2* Glu: 0.87, GABA: 1.4). Separately from burst modulation, splicing rate dynamics are also of interest particularly from data containing nascent and mature mRNA counts, to, for example, understand the splicing kinetics over germ cell development/maturation [57] (Figure 1e). Previous analysis inferred changes in splicing by observed changes in the nascent/mature mRNA ratio (Figure 1f). However, multiple alterations to transcription as well as splicing mechanisms can result in such ratio differences (Figure 1e). With *Monod*-inferred parameters, we can identify when the splicing rate itself is likely modulated, which we apply to genes in the original study, the spermatogenesis-related *Bnc2* and *Dmrt1*, as well as cell cycle-related *Ccnd3* [57]. We find greater differences in splicing rate between the ovary and testes germ cells in the E13 stages for *Ccnd3* and *Dmrt1* as posited by Mayère et al. [57] (Figure 1g). Given these parameters, we also identified novel gene candidates which displayed enhanced, rather than reduced, splicing kinetics during development including the mitotic-spindle-associated *Nusap1* and topoisomerase-encoding *Top2a* with roles in transcription and DNA replication [58], demonstrating increased splicing rates between the E12 and E13 stages (Figure 1h).

### 2.3 *Monod* identifies strategies of resistance and recovery

The expanded capabilities of biophysical DE analysis also allow us to explore more complex biological settings, where it is less clear which mechanisms are expected to change and which genes provide actionable targets for followup experiments. For example, in disease contexts it is important to understand how aberrant behaviors in tumor or cancer cell populations promote tumorigenesis and drug resistance, to develop targeted therapeutics and interventions [59]. Likewise, understanding cellular strategies of recovery post-treatment aids in the selection of relevant therapies and promotion of desired cell behaviors [60].

To first explore how cells may promote undesirable behaviors such as drug resistance and malignancy, we analyzed patient samples of pancreatic ductal adenocarcinoma (PDAC) tumors. Single-nucleus RNA-seq (snRNA-seq) was performed, to improve retention of malignant cell populations, on patient samples with and without neoadjuvant treatment [61] (Figure 2a). We analyzed treated samples that received “multicycle chemotherapy (FOLFIRINOX) followed by multifraction radiotherapy with concurrent fluorouracil (5-FU)/capecitabine”(CRT) [61], or CRT and losartan (CRTl) treatment (Figure 2a). Given the variability in patient backgrounds, we compared the biophysical parameters inferred for a CRT (and CRTl) patient sample to the parameters of three untreated samples to identify robust changes per treated patient. We compared the malignant cells (given existing annotations [61]) across the tumor samples, and parameters were inferred for the bursty model of transcription with length-biased sampling over a shared set of 3,000 genes, with 1,030 genes retained based on goodness-of-fit metrics across samples.

**Figure 2:**
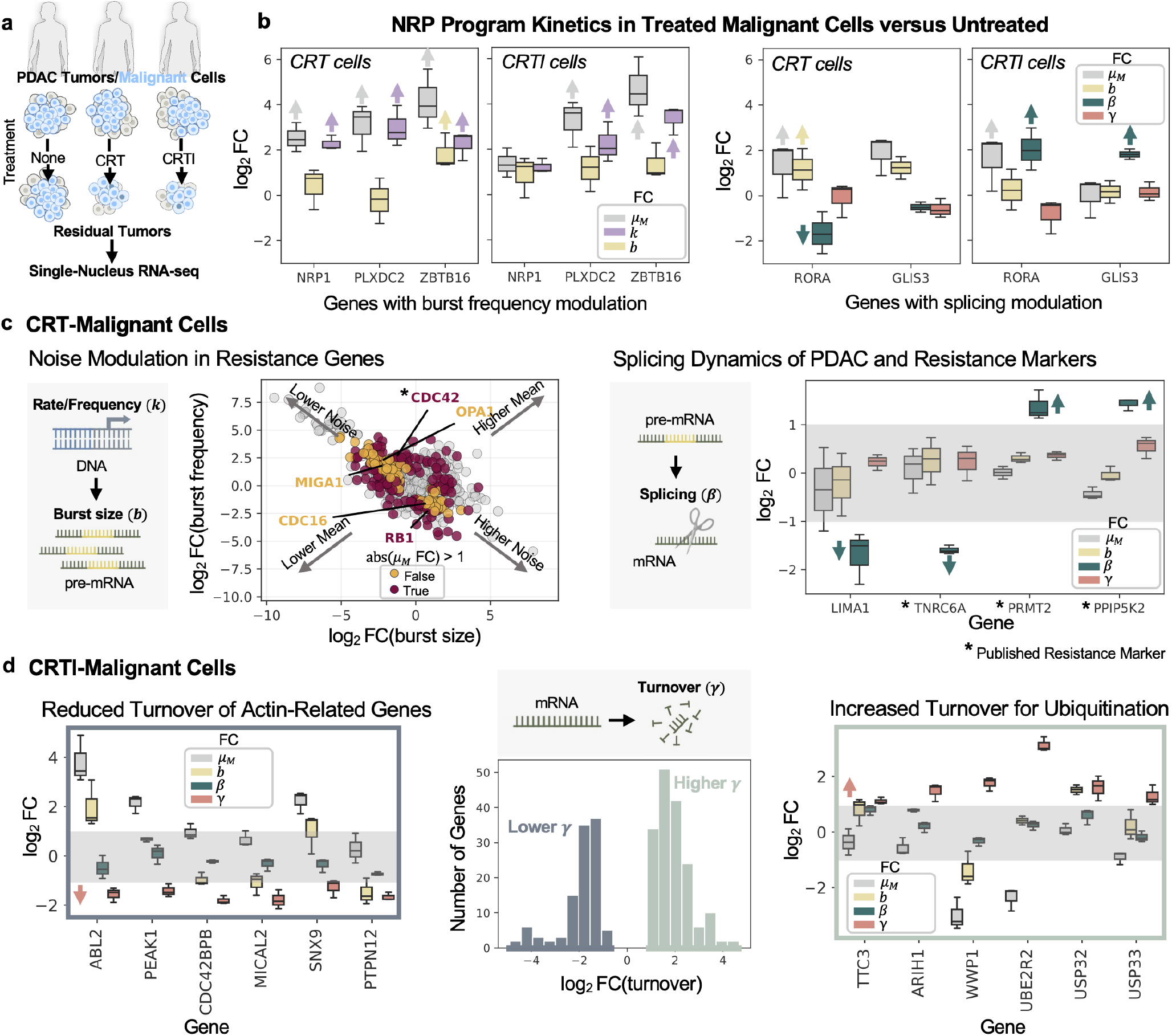
Resistance and proliferation in malignant PDAC cells. **a**. Diagram of snRNA-seq data from residual PDAC tumors [61] post-treatment (or untreated, as in first column) obtained from frozen patient samples. Illustrations adapted from NIAID NIH BIOART Source [129, 130]. **b**. (Left) Boxplots showing log_2_ fold change (FC) of mean (mature) mRNA expression alongside burst size and burst frequency FC, between denoted condition and untreated samples (n=3). Whiskers denote 1.5x the interquartile range where lower boundary is the minimum value and upper boundary is the maximum value, excluding outliers. (Right) Boxplots showing log_2_ FC of mean (mature) mRNA expression alongside burst size, splicing rate, and degradation rate, between denoted condition and untreated samples (n=3). Whiskers denote 1.5x the interquartile range where lower boundary is the minimum value and upper boundary is the maximum value, excluding outliers. **c**. (Left) Plot of log_2_ FC in burst frequency versus burst size for all non-rejected genes in CRT-malignant cells. Genes with log_2_ FC *>* 1 in both/either parameter colored. Red denotes mean expression abs(log_2_ FC) *>* 1 and yellow denotes abs(log_2_ FC) *<* 1 (little/no change). (Right) Boxplots showing log_2_ fold change (FC) of mean (mature) mRNA expression alongside burst size, splicing rate, and degradation rate, between CRT cells and untreated samples (n=3). Whiskers denote 1.5x the interquartile range where lower boundary is the minimum value and upper boundary is the maximum value, excluding outliers. * denotes published gene associated with 5-FU/chemo-resistance. Gray box denotes abs(log_2_ FC) *<* 1. **d**. (Left) Boxplots showing log_2_ fold change (FC) of mean (mature) mRNA expression alongside burst size, splicing rate, and degradation rate, between CRTl cells and untreated samples (n=3). Whiskers denote 1.5x the interquartile range where lower boundary is the minimum value and upper boundary is the maximum value, excluding outliers. Gray box denotes abs(log_2_ FC) *<* 1. (Middle) Histogram of genes with reduced turnover (log_2_ FC *<* −1) or increased turnover (log_2_ FC *>* 1). (Right) Same boxplot as Left, for genes with increased turnover.

As identified in the original publication [61], CRT and CRTl-treated tumors show increased mean expression (where *µ*_*M*_ are scaled pseudobulk mature mRNA counts) of genes belonging to a neural-like progenitor (NRP) malignant program, as compared to the untreated samples (Figure 2b, see Methods Section 9.2.6). These genes relate to neuronal development, migration, and adhesion, which can increase invasiveness and lead to poor prognoses [61]. In addition to mean changes, we also identified which processes were likely modulated, denoted by the arrows in Figure 2b. Burst frequency (or transcription rate *k*) modulation was determined as described above (see Methods Section 9.2.2) (Figure 2b, left), distinct from scenarios where splicing rate specifically had the greatest contribution to the observed fold change in mean expression (Figure 2b, right).

We can additionally identify genes, not covered in previous analyses, which specifically demonstrate transcription-based modulation, as opposed to splicing or degradation-based, potentially through noise regulation. In CRT-treated malignant cells, the tumor suppressor gene *RB1* [62] showed distinct downregulation through burst frequency (transcription rate), while *CDC42*, which regulates cell division and can aid resistance in 5-FU-treated colorectal cancers [63], demonstrated increased burst frequency (Figure 2c, left). A set of genes related to mitochondrial fusion and spindle organization, *MIGA1, OPA1*, and *CDC16* [58], did not have discernible changes in mean, but did demonstrate modulation of burst frequency and/or size suggesting strategies to increase or decrease their transcriptional noise (Figure 2c, left). Their modulation echoes previous literature, which has described the enrichment of genes related to organelle fusion/fission and spindle structure in 5-FU resistant colorectal cancer cells [64]. Beyond transcriptional control, we also find marked splicing rate changes in genes associated with drug resistance or tumorigenesis, where mean expression differences may be low. *TNRC6A* [65], *PPIP5K2* [66] and *PRMT2* (a known PDAC marker) [67] all represent genes associated with 5-FU/chemo-resistance (Figure 2c, right). In particular, increases in *PPIP5K2* and *PRMT2* have been associated with increased drug resistance [66, 67], with both genes also displaying increased splicing rates. Decreased splicing was also observed for *LIMA1*, where lower expression has been associated with increased tumorigenicity [68]. This presents new gene candidates where splicing regulation, in addition to transcriptional machinery, is a potential regulatory step for enacting therapeutic control [59].

In CRTl-treated malignant cells we can similarly identify changes in noise modulation, for example with decreasing noise for *TIMP2*, previously found to be regulated by losartan treatment [69], and increased noise in the proto-oncogene *MET* [70] (Supplementary Figure 4). Notably, we can also extract genes where mRNA turnover is uniquely affected. We refer to the learned degradation rate as general ‘turnover’, as snRNA-seq may better capture nuclear export as opposed to decay/degradation. Within the set of genes with decreased/slower turnover, we found genes related to regulation of actin/cytoskeletal dynamics encoding actin-associated kinases, *ABL2, PEAK1*, and *CDC42BPB*, as well as *MICAL2* and *SNX9* (related to actin depolymerization/polymerization respectively) [58] and *PTPN12* (related to actin reorganization and a potential tumor suppressor [71]) (Figure 2d, left). Such turnover-based strategies have been similarly observed in proliferating cells, where beta-actin mRNA turnover decreased with faster growth rates [72], and in tumor cells where actin-binding proteins can accumulate in the nucleus promoting development of invasive structures [73]. In the opposite direction, we find genes encoding E3 ubiquitin-protein ligases, *TTC3, ARIH1* and *WWP1*, and ubiquitin-conjugating enzymes and deubiquitinating enzymes, *UBE2R2* and *USP33/32* [58], to have increased turnover/export suggesting potential control of protein ubiquitination/turnover by altered mRNA turnover (Figure 2d, right). Together, these results also present gene candidates regulated through their export and stability that can promote resistance and proliferation of these malignant cells.

In addition to understanding how cells develop tumorigenic and resistance properties, exploring their ability to compensate and recover from treatment is also important to enable more effective therapies and treatment plans. For example, the intestine is one of the organs most sensitive to radiation damage: understanding how it heals following radiotherapy is important for promoting healthy recovery and survival of patients treated for abdominal, gastrointestinal, pelvic, and retroperitoneal tumors [60]. Particularly interesting is how immune cells are affected by and respond to radiation treatment. T cells, which are essential actors in the body’s natural response to tumor control [74, 75], are required for successful radiotherapy but are also believed to be damaged during treatment. While studies have investigated the transcriptomic landscapes of T cells following radiotherapy [60, 74], we suggest that a fuller understanding of the regulation of key genes, as revealed by biophysical parameter changes, could inform how therapeutics may be designed and more efficiently administered to enhance recovery from radiotherapy.

We re-analyze data from a study of the transcriptomic response of the intestinal microenvironment to radiation-induced intestinal injury (RIII) in wild-type C57BL/6J male mice [60] (Figure 3a). Mice were treated with 15 Gy of abdominal radiation, and intestinal samples were collected pre-radiation treatment (day 0) and at several time points following treatment (days 1, 3, 7 and 14 post treatment). We selected T cells based on the original study’s annotations, and inferred biophysical parameters for samples (where each sample is a unique mouse) with at least 100 T cells, leaving 11 samples across days 0, 1, 3 and 14, with at least two samples per day. We used the bursty model with Poisson technical sampling noise, and after fitting 3,000 genes shared across samples, subset to the 1,182 that passed goodness-of-fit testing in all samples, further correcting inferred parameters for sample-specific offsets (see Methods Section 9.2.7).

**Figure 3:**
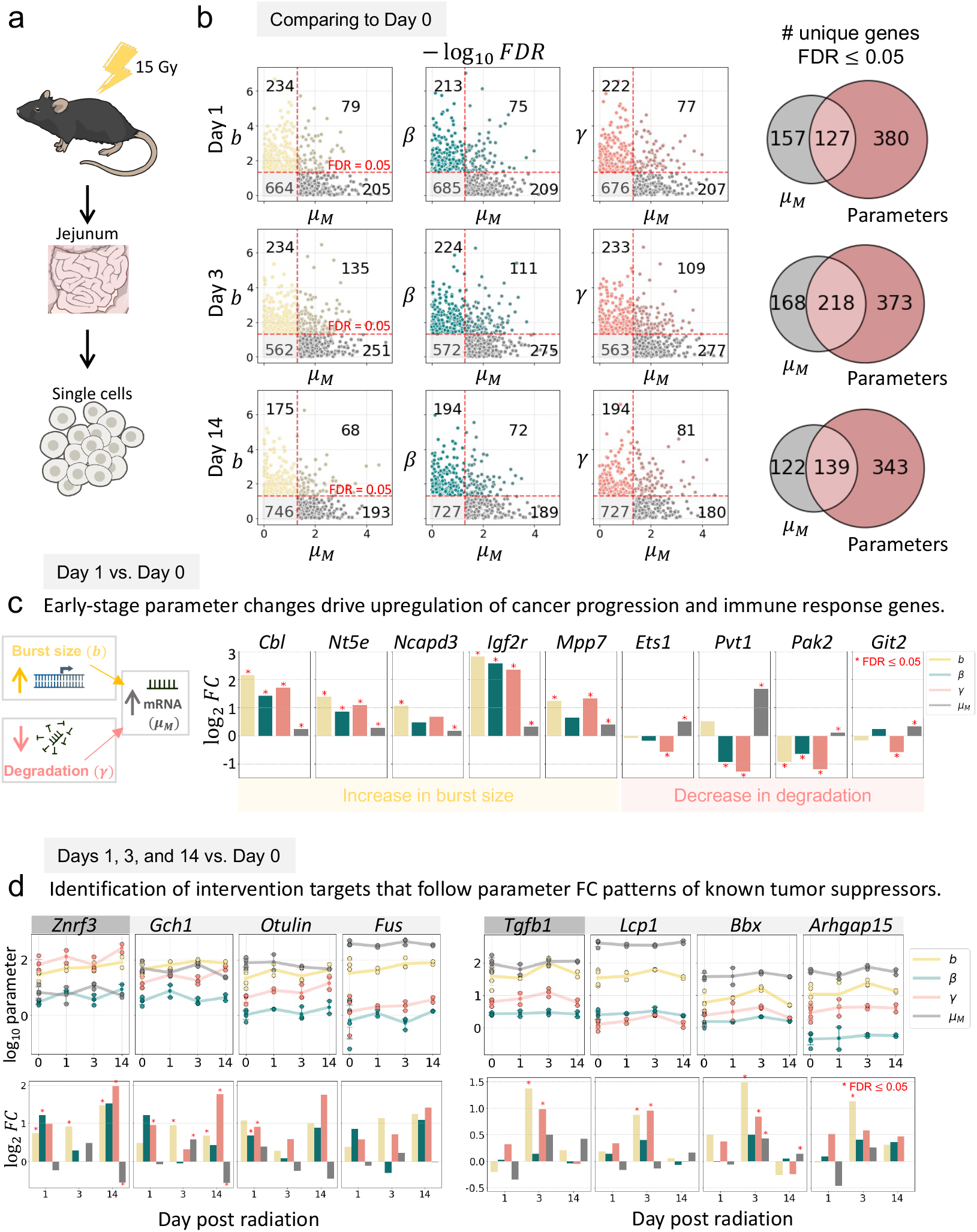
*Monod* analysis of mechanisms of T cell recovery post radiation treatment. **a**. Single cell samples were taken from the intestines of mice exposed to 15 Gy of abdominal radiation before and at several time points after treatment. Illustrations adapted from NIAID NIH BIOART Source [131–133]. **b**. Two-sided *t* -tests comparing T cells at days 1 (n=2), 3 (n=3) and 14 (n=2) post treatment to T cells before treatment (day 0, n=4) flag more genes as significant (false discovery rates, FDR ≤ 0.05) when comparing inferred parameters (*b, β, γ*) than when merely comparing mature RNA counts (*µ*_*M*_ ). In the scatter plots, red dashed lines indicate a significance threshold of an FDR of 0.05. **c**. The log_2_ fold change (FC) of parameters (comparing day 1 to day 0, see Methods Section 9.2.7) show that statistically significant changes in the mature mean expression of genes related to the progression of cancer and immune response can be attributed to statistically significant differences in parameters (i.e., burst size or degradation rate). A red asterisk indicates a FDR ≤ 0.05. **d**. Comparing genes’ parameter changes over the timecourse of recovery can identify similarly regulated genes. The top row shows the log_10_ of parameters and mean mature RNA expression at days 0, 1, 3 and 14. Each o1u6tlined point indicates a separate sample (mouse) with a smaller point at the mean over the log_10_ of samples’ parameters per day. Error bars indicate the interquartile range of the samples’ parameters per day. For certain genes, the log_2_ FCs of each parameter and mean mature expression for days 1, 3 and 14 from day 0 (see Methods Section 9.2.7) exhibit similar patterns as known tumor suppressors (bottom row).

We first show that analysis through the lens of a biophysical model reveals more genes with differences in behavior across the timecourse of recovery. To assess the significance of parameter changes, we performed a *t* -test with unequal variance on the log_10_ of each corrected parameter per gene for day 0 versus all other days. Specifically, we tested for changes in burst size (*b*), relative splicing rate (*β*), relative degradation rate (*γ*) and normalized mature counts (*µ*_*M*_, see Methods Section 9.2.7), correcting *p*-values for the number of tested genes using the Benjamini Hochberg procedure (separately per parameter) to obtain false discovery rates (FDRs). Figure 3b shows that many genes that do not show significant differences in mature counts are detectable via parameter changes: testing the significance of inferred parameters in addition to means allows for the exploration of more genes whose significant difference between days or conditions may lie in regulatory strategy rather than mean expression change. For example, comparing day 1 to day 0, there were 380 genes with significant differences in a parameter but not in mature expression, 127 genes with differences in both, and 157 genes with significant differences in mature expression but not parameters (Figure 3b). We further emphasize that the analysis of parameters does not preclude testing for changes in mean, with our method adding to the list of possible genes for follow-up investigation.

In addition to identifying genes with parameter changes that are not detectable using standard mature expression comparisons, for genes which *do* show significant differences in expression, *Monod* can suggest what cellular regulatory strategy is responsible (e.g., an increase in burst size or a decrease in degradation rate, Figure 3c). For example, we show in Figure 3c (left) five genes which had significant increases between day 1 and day 0 in both burst size and mature RNA counts. As these genes also show increased degradation rate, the observed increase in mature counts is likely due to an increase in burst size (that exceeds the increase in degradation rate). These genes, which are upregulated early in the radiation-response timecourse (day 1 post treatment), have been studied as promising targets in cancer immunotherapy. *Cbl* encodes the E3 ubiquitin ligase Casitas B lymphoma, which regulates immune cells and whose homologue Casitas B lymphoma-b (CBL-b) has been shown to promote immunosuppressive tumor microenvironments (TME) [76]. *Nt5e* encodes 5’-nucleotidase (or Cd73), an immunosuppressive protein whose high expression is also associated with poor outcome of cancer patient’s TMEs [77] and is currently emerging as a novel target for immunotherapy [78]. Other genes whose high expression has been correlated with lower overall survival in cancers, are *Ncapd3*, which is associated with reduced infiltration of immune cells into glioma [79], and the gene that codes for insulin-like growth factor 2 receptor (*Igrf2* ), which was found to be an indicator of poor prognosis in patients with triple-negative breast cancer [80]. On the other hand, the expression of *Mpp7*, which codes for a scaffolding protein, has been shown to have a positive correlation with increased infiltration of T cells in melanomas [81].

Other notable genes show increases in mature RNA expression best explicable by decreases in degradation rate (Figure 3c, right). *Ets1* encodes a transcription factor that is a proto-oncogene that promotes aggressive tumor behavior in a variety of cell types [82]. While not directly associated with cancer progression, the proteins encoded by *Pak2* and *Git2* are known to interact [83], and are crucial for the stability and suppressive function of regulatory T cells [84]. With knowledge about what specific point in the RNA lifecycle (e.g., bursty transcription or degradation) may lead to increased expression of a gene, researchers could design strategies to discourage known tumor promoters and encourage known tumor suppressors for increased efficacy and complementarity of radio- and immunotherapy.

Finally, we can identify genes whose parameters show similar dynamics (fold changes) over the timecourse of recovery from radiation treatment as compared to known tumor suppressing genes (see Methods Section 9.2.7). For example, we found the three genes with fold change patterns most similar to *Znrf3* (Figure 3), whose encoded protein negatively influences cancer progression by inhibiting key proliferative (WNT) signaling pathways [85, 86]; and another three with patterns most similar to *Tgfb1*, which potentially suppress tumorigenesis at early cancer stages but promote tumorogenesis at advanced cancer stages [87]. Of these genes, some have been reported as important in cancer: for example, *Gch1* is a known immunosuppression gene with upregulation previously linked to reduced survival in breast cancer [88]; *Lcp1* encodes a prognostic biomarker in gastric cancer [89]; and *Arhgap15* codes for a protein that is reported to slow colorectal tumor growth [90]. Other identified genes have not been as well-characterized in cancer studies: *Otulin* encodes an immune-response peptidase [91], and *Fus* and *Bbx* are more general DNA damage-response genes [92, 93]. Analysis of biophysical parameters allows one to explore how genes that have been previously implicated in cancer prognoses are experiencing similar regulatory changes in response to radiation treatment, as well as to identify genes that have *not* been previously implicated as targets for follow up therapeutic development.

### 2.4 Model selection and insight about gene regulation strategies

*Monod* currently supports 6 sophisticated yet tractable models of transcription, previously shown to describe and distinguish between features of experimental data [23, 25, 45] for thousands of genes across multiple cell types. We emphasize that *Monod* supports *modular* model comparison, in the sense that one can freely modify assumptions about a specific model component (e.g., whether degradation occurs with a deterministic or stochastic waiting time) to obtain a new model to test against data. We recommend using *Monod* with the bursty model by default, as it captures observed overdispersion [36] and known biophysics [27, 94] while only minimally complexifying the constitutive model. But the bursty model may not *always* fit the data best, and it is worth testing to what extent a less complex, more complex, or simply different model performs better in practice.

In this section, we use *Monod* to analyze mouse brain data with 2,951 genes and six cell types (Allen sample B08 [56]) and show that the bursty model almost always outperforms the simpler constitutive model, and tends to perform fairly well against alternative models (see Figure 4a for schematics of the 5 models we consider in this section). Moreover, we show that it is useful to consider a richer space of transcription models, since specific models may fit specific genes or cell types better or worse, which suggests *Monod*-derived fits yield insight about gene- and cell-type-specific gene regulation strategies. We reject models (for a given cell type and gene) if they fail to pass the goodness-of-fit thresholds described in Section 9.3.

**Figure 4:**
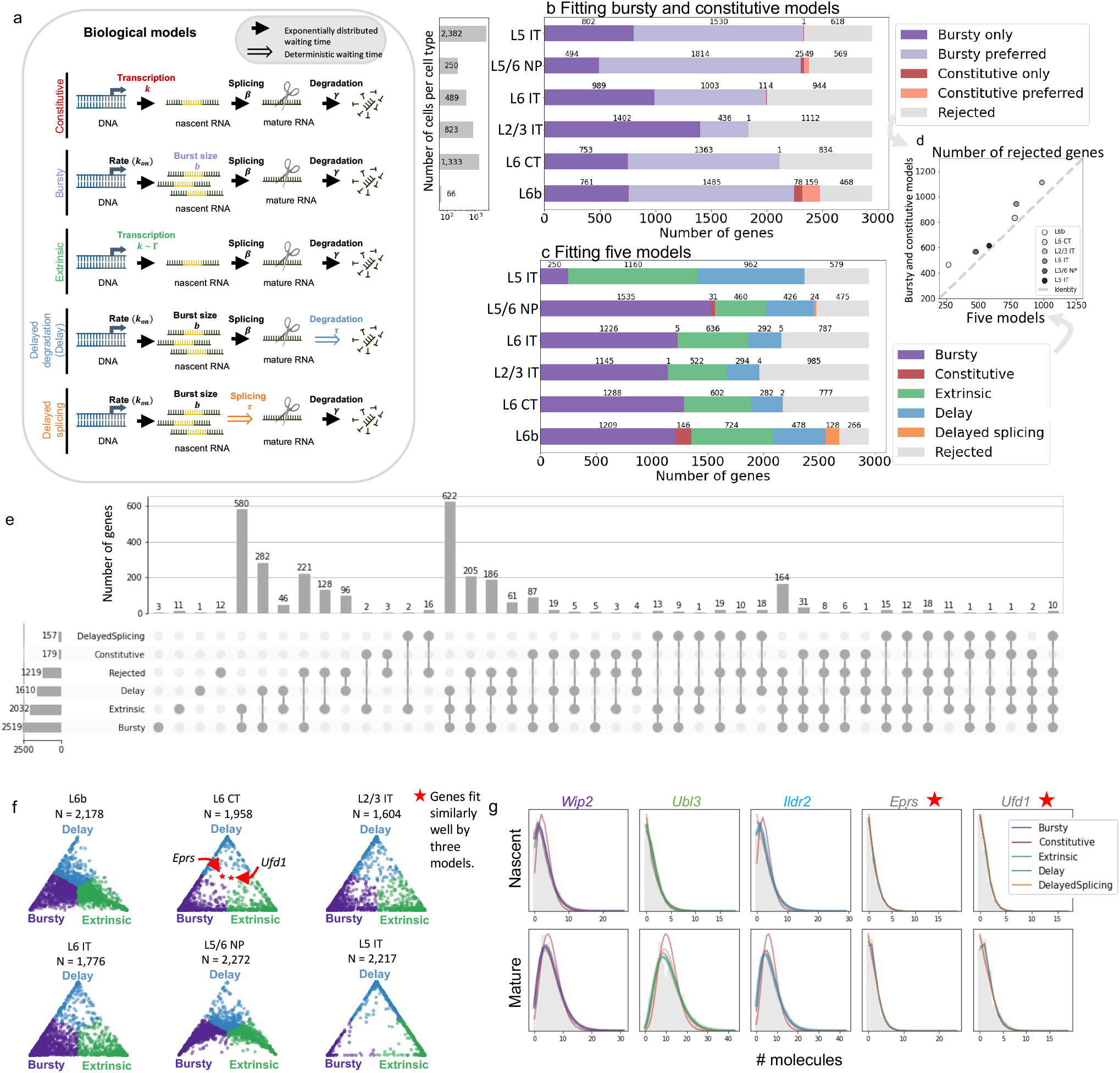
*Monod* facilitates the comparison of different biophysical models on a per gene basis for thousands of genes across cell types. Results are shown for Allen sample B08 [56]. **a**. Diagrams of five of the biological models implemented in *Monod*, varying in transcription and processing kinetics (Section S1.1). **b**. Of the 2,951 fit genes in each of six cell types, most genes display behavior more consistent with the bursty model than with the constitutive model using the Akaike information criterion (AIC) and two goodness-of-fit metrics 9.3. **c**. For the same six cell types and genes, the number of genes that are best fit by each of the five considered biological models, colored by model. Gray indicates genes for which all models were rejected). **d**. Fewer genes are rejected when fitting and comparing five models than fitting and comparing only the bursty and constitutive models. **e**. The same gene can exhibit different behavior in different cell types: the upset plot shows the number of genes that are best fit by a given model or combination of models across the six cell types. **f**. Ternary diagram of genes’ normalized AIC weights for the three most commonly selected models, colored by the model with the largest AIC weight. Only genes for which none of these models were rejected are displayed, with the number indicated. **g**. Predicted marginal nascent and mature distributions for each model plotted over normalized empirical histograms of molecule counts for the indicated genes in layer 6 neurons (L6 CT). *Eprs* and *Ufd1*, which had similar AIC weights for the bursty, extrinsic, and delay models, are noted with a red star in the ternary diagrams.

We first compare the constitutive model to the bursty model. For many genes, only the bursty model passes goodness-of-fit thresholds. When neither the bursty nor the constitutive models are rejected for a gene, the bursty model is usually a better fit according to Akaike information criterion (AIC, see Section 9.3, Figure 4b). It is extremely rare that the constitutive model fits the data better, perhaps unsurprisingly as the constitutive model is a special case of the bursty model and the overdispersion of single-cell count data is well known, but diverging from previous studies that advocate the Poisson model [95].

Although the transcriptional behavior of many genes is consistent with the bursty model, comparing all five of the (comparably complex) proposed models reveals that the extrinsic, delayed degradation, and delayed splicing models are better fits for some genes in different glutamatergic cell types (Figure 4c). This comparison also suggests differences between cell types’ transcriptional regulation strategies, with more genes best fit by the extrinsic model in layer 6 intratelencephalic (L6 IT) neurons than in layer 5/6 near-projecting (L5/6 NP) neurons, for example (Figure 4c). Further, including the additional models increases the number of genes in each cell type for which one or more models pass goodness-of-fit thresholds: fewer genes are rejected by all models when considering more models (Figure 4d). As all models were compared in a standard way using the same data, fitting pipeline, and statistical rejection criteria, the discrepancies between proposed models could be evidence of real biological variety in transcriptional and post-transcriptional regulation mechanisms.

While cell type comparisons show distinctions between the number of genes best fit by each model in a given cell type, *Monod* also provides the tools to investigate how genes exhibit expression patterns consistent with different transcriptional models across cell types (Figure 4e). We found the model with the largest AIC weight for every gene in every cell type, then for each gene collapsed models over cell types to a set of unique models. The upset plot in Figure 4e shows how many genes exhibited each combination of models preferentially across the six cell types. For example, only three genes exhibited *only* bursty behavior preferentially, while 580 genes were best fit by the bursty model in at least one cell type and by the extrinsic model in at least one other cell type. All but 27 genes exhibited expression trends described best by at least two unique models, with the most common set (622 genes) being bursty, extrinsic, and delayed degradation. Interestingly, no genes were best fit by only the delayed splicing or only the constitutive model across the six cell types (Figure 4g).

While these trends may reflect biological differences, they may also highlight differences in the mutual identifiability of the proposed models. For example, the bursty, extrinsic, and delayed degradation model all account for correlation between nascent and mature counts, while in the constitutive and delayed splicing model, nascent and mature counts are uncorrelated. Thus the genes that are best fit by the three models or some subset may be in fact displaying similar behavior across different cell types, with the correlation between their nascent and mature counts, ruling out the constitutive and delayed splicing models, as the salient feature driving the fits (similarly to Gorin et al. [45]).

To explore this question, we can asses the identifiability of models, or the extent to which the “best” fit model is “better” than other models. While we can assign a gene to the model with the largest AIC weight (Section 9.3), there is not always a clear distinction between model fits. To illustrate this, we show a simplex of normalized AIC weights for the bursty, extrinsic, and delayed degradation models, colored by the assigned regulatory model: the weights for the “best” fit models are sometimes arbitrarily close to the other two models’ weights (Figure 4f). In this way, genes with clear preference for a given model can be distinguished from those which are similarly fit by two or more models. These results can also be verified by visualizing model-based probabilities and normalized empirical count histograms. Figure 4g shows genes that were not rejected by only the bursty model (*Wipf2* ), only the extrinsic model (*Ubl3* ), and only the delayed degradation model (*Ildr2* ) in layer 6 neurons (L6 CT). *Eprs* and *Ufd1*, indicated with red stars in Figure 4f, were fit similarly well under the three models. In the histograms for *Wipf2, Ubl3*, and *Ildr2*, we underline the (Poisson) constitutive model’s poor ability to recapitulate the overdispersed data, consistent with Figure 4b-c.

*Monod* makes fitting subtly distinct mechanistic models of transcription for thousands of genes straightforward, thereby leading to increased ease of model comparison and selection. Performing goodness-of-fit and likelihood tests on *Monod* fits provides a route for discerning between biophysical models that may have generated the data. However, we note that definitive model selection is beyond the scope of *Monod* : rather than to come to definite conclusions, *Monod* should be used to explore data and to compare and generate hypotheses that can later be tested experimentally.

### 2.5 Principled integration of multiple modalities

How should one reconcile or combine scRNA-seq data with other types of single-cell data, like single-nucleus RNA sequencing (snRNA-seq) data [96, 97], single-molecule fluorescence data (e.g., obtained via RNA seqFISH+ [98]), and data reflecting information beyond RNA counts, like protein abundances [99,100] and chromatin accessibility [101–103]? Since different modalities may be better at capturing various aspects of the underlying biology—for example, snRNA-seq data is enriched for intronic reads [13], and hence one might expect it to be more informative about nuclear processes— integrating data across modalities allows different advantages to be simultaneously exploited.

Currently popular approaches to single-cell data integration are largely inspired by machine learning, and typically use a shared latent space to make data from different modalities comparable [104–108]. However, these approaches are not without problems; Tyler et al. have argued that they often remove biologically meaningful variation [109], and more generally it is unclear to what extent heuristic approaches to single-cell data analysis are trustworthy [15].

We advocate an alternative approach rooted in the mechanistic worldview. First, one defines a generative model whose biological component is shared, but whose technical/observation component differs across modalities; then, one fits the model to data; finally, one uses Bayes’ rule to combine parameter estimates from different modalities according to their uncertainty (Figure 5a). *Monod* supports this procedure by quantifying the uncertainty of its parameter estimates (Section 9.1.3). In the rest of this section, we walk through an example of this approach to integration.

**Figure 5:**
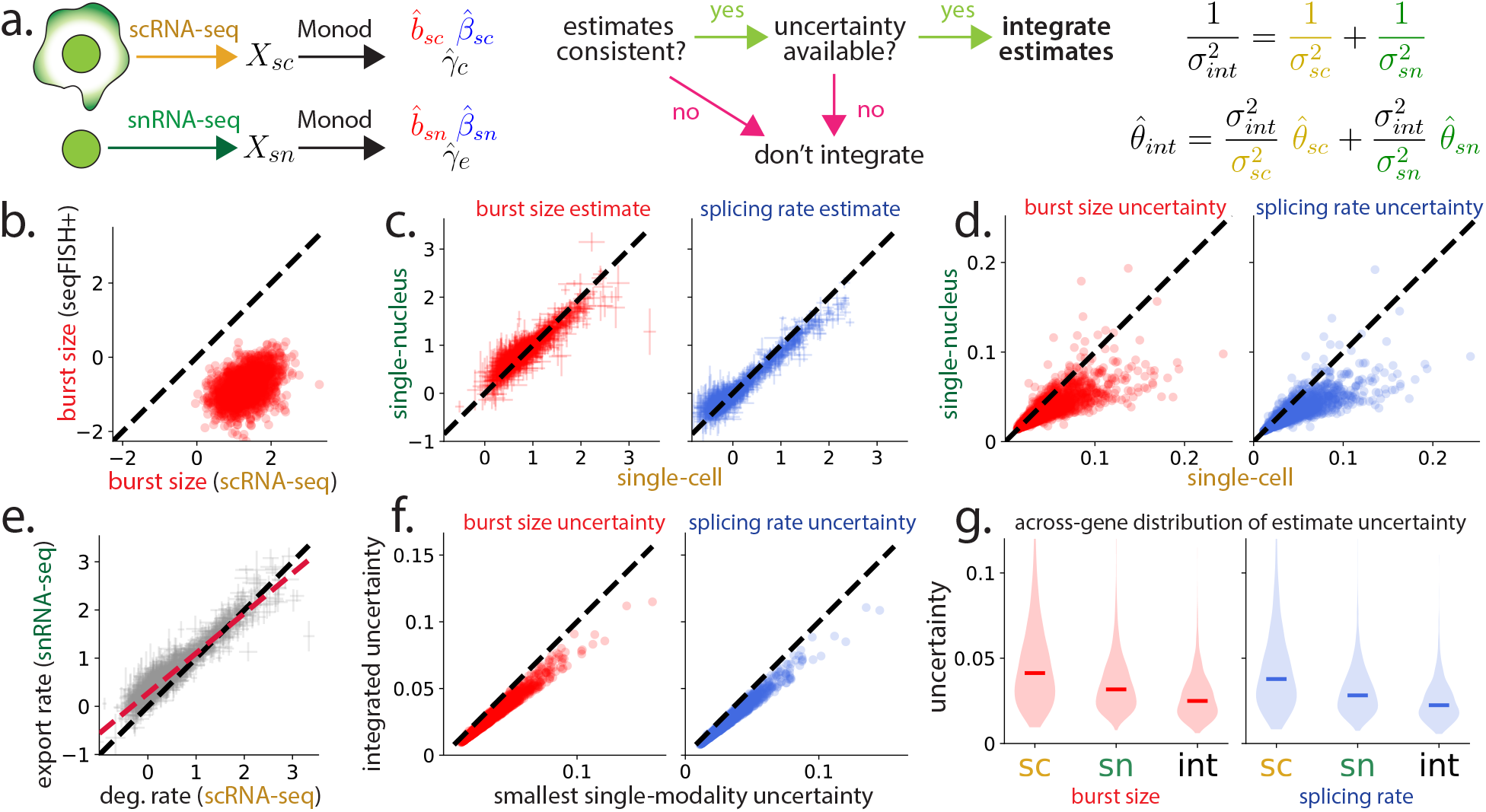
*Monod* ‘s quantification of parameter estimate uncertainty supports a biophysically meaningful approach to integrating data from different modalities. **a**. Schematic depiction of integration, with fitting single-cell and single-nucleus data to the bursty model used as an example. Integrating data from different modalities is useful (when it is possible) because it reduces the uncertainty of parameter estimates. **b**. Consistency of seqFISH+ and scRNA-seq burst size estimates for 1, 964 genes in two unrelated mESC data sets, log_10_ space. Black line: identity. They are not very consistent (Pearson *R* = 0.424). **c**. Consistency of snRNA-seq and scRNA-seq estimates of burst size (Pearson *R* = 0.918) and splicing rate (Pearson *R* = 0.972) for 1896 genes for which the bursty model fit well. Error bars: 99% confidence regions determined by *±*2.576 *σ*. **d**. Uncertainties (i.e., standard deviations of posterior in log_10_ space) of snRNA-seq and scRNA-seq estimates of burst size and splicing rate. **e**. Consistency of scRNA-seq and snRNA-seq inferred ‘degradation’ rates. Crimson line: best fit. Black line: identity. The snRNA-seq data have a positive offset, and mean higher than the scRNA-seq data (two-sided *t*-test; *{t, p}* = {−11, 2.1 *×* 10^−27^}). **f**. Uncertainty after integrating snRNA-seq and scRNA-seq estimates vs. lowest single-modality uncertainty across all genes. Note that all points are below the identity line, which reflects integration reducing uncertainty. **g**. Distribution of estimate uncertainty before and after integration. Horizontal lines: medians.

Before integration, one must check whether estimates from different modalities are consistent, since inconsistencies may indicate model misspecification issues. We consider two example comparisons: between seqFISH+ [110, 111] and 10x v3 scRNA-seq data [112], both collected from mouse embryonic stem cells (mESCs) in unrelated experiments; and between snRNA-seq and scRNA-seq data, both generated by 10x Genomics from the same mouse brain tissue sample [113, 114]. seqFISH-derived burst size estimates obtained via the procedure of Takei et al. [111] only coarsely agreed with *Monod*-derived estimates from analogous scRNA-seq data (Figure 5b), possibly due to some combination of measuring slightly different features of the underlying biology, and not adequately accounting for fluorescence-related technical noise. Curiously, the *averages* obtained from seqFISH and scRNA-seq agreed to an even lesser degree (Supplementary Figure 9), suggesting that the burst sizes are, in a sense, more reliable than the raw expression values, and that appropriately accounting for probe signal in the spirit of [29] may, in the future, help integrate the underlying biophysics. Meanwhile, *Monod*-derived estimates of burst size and splicing rate based on snRNA-seq and scRNA-seq mouse brain data were extremely consistent (Figure 5c), which suggests that integration may be advisable in the latter case.

Interestingly, according to *Monod* ‘s quantification of parameter uncertainty, estimates of burst size and splicing rate from snRNA-seq data are usually more precise than the corresponding scRNA-seq-based estimates (Figure 5d; 84.9% of burst size estimates and 90.7% of splicing rate estimates are more precise), which is consistent with the intuition that snRNA-seq data ought to be more informative about nuclear processes. We obtain this result in spite of the rather smaller size of the single-nucleus dataset (Supplementary Table 8). On the other hand, on biophysical grounds we expect snRNA-seq data not to be informative about RNA degradation, and instead for its mature RNA decay rate fits to measure nuclear export (Section 9.4.2). Consistent with this intuition, we find mature RNA decay rates significantly different for each data type (Figure 5e), with snRNA-seq-derived rates usually being somewhat higher.

Let *µ*_*i*_ denote the average parameter estimate given data from some modality, and *σ*_*i*_ denote the corresponding uncertainty. Assuming a sufficiently large number of cells are available, Bayes’ rule implies (see Section 9.4) that the integrated estimate *µ*_*int*_ and its uncertainty *σ*_*int*_ satisfy

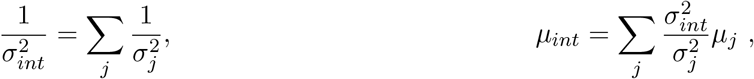

i.e., that the uncertainty of the integrated estimate is smaller than each of the single-modality uncertainties. Applying the above formulas to estimates derived from snRNA-seq and scRNA-seq, we find that uncertainty is reduced, sometimes by up to 30% (Figure 5f, g).

### 2.6 Assessing loss of biological signal after pre-processing

Because *Monod* fits mechanistic models to single-cell data, it enables a precise accounting of variation—in particular, variation due to (i) technical effects, (ii) cell type heterogeneity, and (iii) intracellular noise. One useful way to summarize this accounting is in terms of a noise decomposition in the spirit of previous work [48, 115], which expresses the coefficient of variation (CV) matrix associated with the data in terms of contributions due to each type of variation (see Section 9.5). For each gene and for three types of overall variation one can track (nascent variance, mature variance, and nascent-mature correlations), this decomposition allows one to generate a set of three weights that determine the “fraction” of overall variation due to each source (Figure 6a, upper).

**Figure 6:**
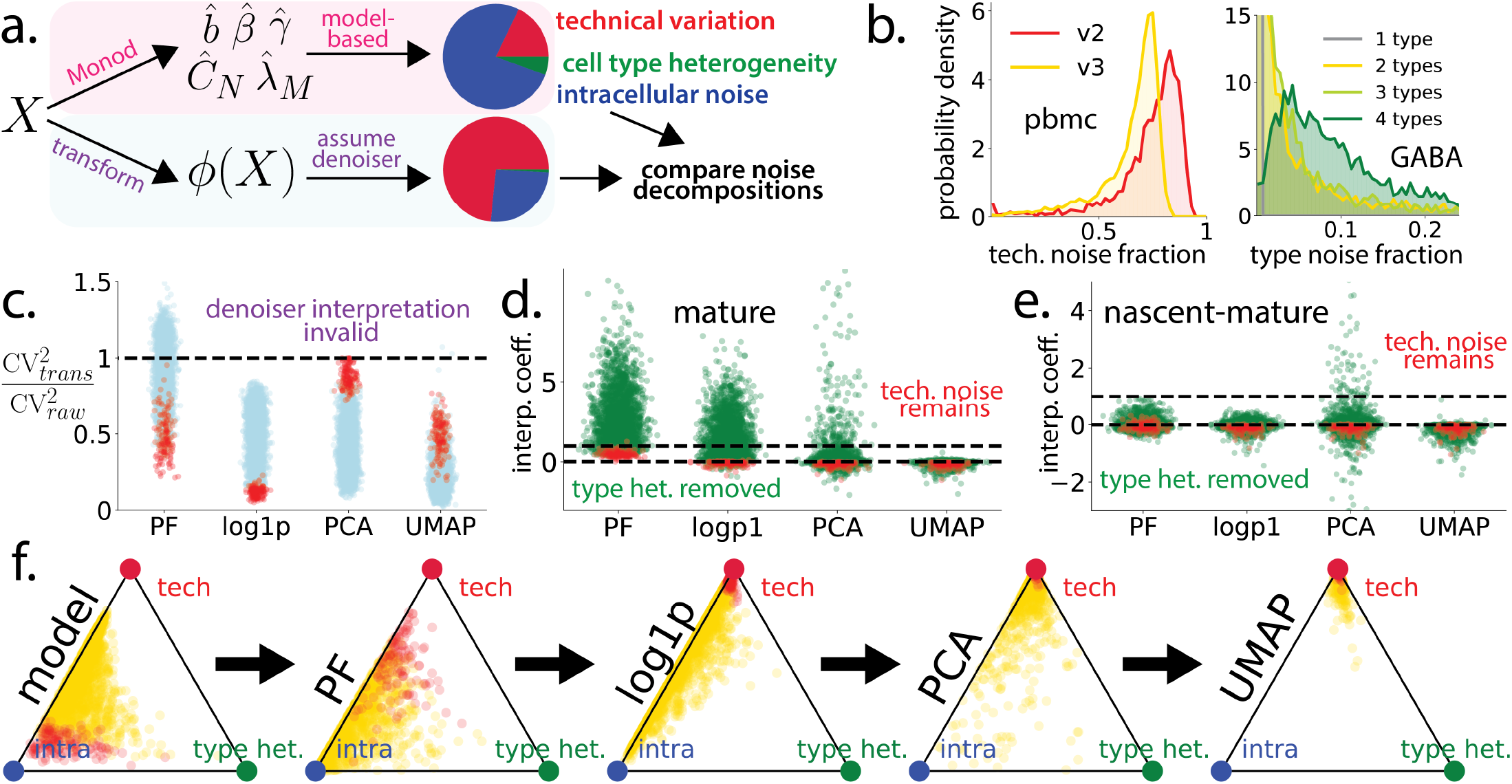
*Monod* enables quantitative assessment of common data transformations. *Monod* can estimate how much observed variation is due to technical effects, cell type heterogeneity, and intracellular noise; this noise decomposition can be used to assess whether common data transformations effectively erase biological signal. **a**. Schematic of the idea: a noise decomposition based on *Monod*-derived fits (in this figure, to the bursty model) can be compared to one that assumes a data transformation *ϕ* (e.g., PF, log1p, PCA, UMAP) effectively removes technical noise. **b**. The *Monod*-derived noise decomposition behaves as expected. Technical noise fractions derived from v2 PBMC data (red; 2500 genes, 3 cell types) tend to be higher than those derived from matched v3 data (gold). Cell type heterogeneity fractions tend to be higher when more cell types are included, GABAergic neuron data (2951 genes, 4 cell types). **c**. Applying many common transformations to scRNA-seq data tends to reduce the mature RNA squared coefficient of variation (CV^2^; 64.1% for PF, 100% for log1p, 100% for PCA, 99.9% for UMAP), making a “denoiser” interpretation reasonable. Red dots: highly expressed (mature mean in 95th percentile) genes. This and the following panels depict glutamatergic neuron data (2951 genes, 6 cell types). **d**. Post-transformation mature CV^2^ tend to violate a model-agnostic upper bound *b*_*upper*_ or lower bound *b*_*lower*_ in an example pre-processing workflow that sequentially applies four data transformations (PF→log1p→PCA→UMAP). Mature CV^2^ were written in the form (1 − *c*) *b*_*lower*_ + *c b*_*upper*_, and values of *c* are plotted. *c >* 1 means not all technical noise was removed, while *c <* 0 means at least some cell type heterogeneity was removed. **e**. Analogous plot for a measure of nascent-mature variation (see Section 9.5). **f**. Noise decomposition weights (mature RNA) throughout example workflow visualized as triangle plots. Applying more “denoising” transformations misattributes more observed variation to technical effects (points concentrate in top corner).

These noise fractions behave as one might intuitively expect. For example, 10x v2 human blood cell (PBMC) data is generally noisier than matched 10x v3 data, and accordingly more of the overall variation is attributed to technical noise (Figure 6b, left). When a GABAergic neuron data set is analyzed with more cell types included (i.e., with data from at least one of the 4 cell types left out), more variation is attributed to cell type heterogeneity (Figure 6b, right).

Even if one is not especially interested in fitting mechanistic models to transcriptomic data, *Monod* ‘s accounting of variation allows it to play an interesting role: it can diagnose the extent to which potentially biologically meaningful variation is lost or distorted after data is pre-processed. This function of *Monod* is important, since typical transcriptomic workflows involve a large number of heuristic correction steps [6], including “cell size” normalization, proportional fitting (PF), log-transformation (log1p), and dimensionality reduction techniques like principal component analysis (PCA) and UMAP [116]. At least part of the motivation for performing these pre-processing steps is to “denoise” the data—i.e., remove or reduce variation due to technical effects—so that the input to downstream analyses is more biologically meaningful. Here, we use *Monod* and glutamatergic neuron data (2951 genes, 6 cell types) to interrogate this motivation.

If we take the “denoising” interpretation of pre-processing steps at face value, the CV matrix associated with the transformed data should reflect cell type to cell type and intracellular variation, but not technical noise (see Section 9.5), which yields an alternative noise decomposition (Figure 6a, lower). Interpreting transformations as denoisers appears to be largely reasonable, since for most genes standard transformations, applied *individually* rather than sequentially, reduce the CV (Figure 6c).

On the other hand, CVs of transformations applied sequentially often appear to indicate a loss of biologically meaningful variation. Assuming transformations remove some combination of technical and intracellular noise but not cell type heterogeneity, which is usually important for downstream analyses [6], one can derive fairly model-free upper and lower bounds for the CV. Upper bound violations indicate that not all technical noise was removed, while lower bound violations indicate at least some variation due to cell type heterogeneity was removed. In an example workflow where four transformations (PF, log1p, PCA, and UMAP) were sequentially applied to the glutamatergic neuron data, lower bound violations are extremely common after the last two transformations (Figure 6d). Similar statements can be made about correlations between nascent and mature RNA (Figure 6e), and nascent CVs (not shown). Visualizing the transformation-associated noise decompositions suggests that, after more pre-processing of the raw data, more observed variation is misattributed to technical effects (Figure 6f). These transformations are heuristic, and the “correct” way to analyze their implicit assumptions is far from clear. However, we obtain similarly severe violations of model-free bounds using a simpler, single-species noise baseline (Section 9.5.5), suggesting that such common practices as, e.g., utilizing normalized data in lieu of raw data [109, 117], benchmarking methods based on the PCA space [14], or directly analyzing the UMAP embedding [15, 16] may all distort the signal of interest in the underlying data to an unacceptable degree.

Although this analysis focuses on the distributions of individual genes, it yields additional troubling implications for the analysis of gene–gene relationships. The discovery of gene regulatory networks (GRNs) and gene modules often relies on leveraging gene–gene correlation matrices (Section 9.5.6). We often do not know about the activity and identifiability of these relationships *a priori* : even if regulatory motifs are present in the underlying biology, they are typically mediated by unobserved proteins. Interestingly, the collection of multimodal data provides a natural positive control: nascent and mature RNA *are* causally related; their correlation is higher than typical gene–gene correlations (Supplementary Figure 15). However, the iterative application of transformations substantially distorts them. Counterintuitively, the supposed removal of technical noise often decreases or changes the sign of nascent/mature correlations (Supplementary Figure 16). As a consequence, their strong signal is “diluted”: for instance, of the genes with high nascent/mature correlations (in top quartile of the mature/mature correlation distribution), the values stay in the top quartile only 60–80% of the time a transformation is applied (Supplementary Figure 17 and 18). For the lower-magnitude nascent/mature correlations, the preservation appears to be considerably worse, and the transformations do not recover the biological signal.

The troubling conclusion is that these known, simple, and biologically meaningful molecule– molecule relationships cease to be distinctly identifiable as the *consequence* of a standard preprocessing step for analysis. In contrast, the “underlying” biological correlations implied by successful *Monod* fits are strictly higher than those observed in the raw data, matching intuition (Supplementary Figure 19). *Monod* cannot yet model gene–gene relationships. However, there is existing work on biophysical GRN inference [118], and we anticipate that these problems are amenable to a mechanistically satisfactory analysis by treating coupling through co-bursting modules [119] or underlying chromatin states [120].

## 3 Discussion

The *Monod* method we have proposed facilitates principled, mechanistic inference and treatment of multimodal scRNA-seq data, that redefines common single-cell analysis tasks through biophysical data representation. Our work also demonstrates mechanistic inference at scale from relatively standard sequencing data types, that would otherwise entail more laborious and error-prone protocols to measure the relevant kinetics [121, 122]. While *Monod* is currently restricted to only a handful of transcriptional models that are tractable by quadrature, technical noise models comprising catalysis or dropout, the two modalities of nascent and mature RNA, correlation structure that does not include inter-gene relationships, and heterogeneity structures comprising a single homogeneous cell type at a time, many of these assumptions can be relaxed and extended. This is in contrast to current approaches to single-cell RNA sequencing analysis that correspond to much more simplistic, unrealistic, and opaque models of biology that make contradictory assumptions and violate fundamental constraints. Advancements in combining machine learning techniques with mechanistic inference [55, 123–125] demonstrate how the *Monod* framework could be extended in the future to more complex biophysical settings and models without analytical solutions. Likewise, the biophysical models in *Monod* provide a basis for integrating other single-cell measurements, such as chromatin information [120] and protein counts [37, 124].

We also note that all examples here have demonstrated inference across predefined, homogeneous populations of statistically equivalent cells; the design of *Monod* presupposes it is applied downstream of cell type identification. However, there are potential advantages to applying this framework upstream: the standard procedure for clustering or cell-type determination is based on mature mRNA counts only, and without a model of the data in mind, it is not necessarily clear how to take advantage of nascent mRNA data [31]. This results in discarding a large fraction of reads containing valuable information, and raises unsolved problems regarding the “correct” way to, e.g., co-cluster single-nucleus and single-cell RNA sequencing data, which have substantially different RNA distributions.

As an alternative, we can use the biophysical basis of *Monod* to define the clusters of interest, i.e., learn “clusters” or “cell types” of cells which display similar dynamics for the cellular processes of interest [31]. This amounts to fitting a mixture model representation of the model class we consider, to learn what populations of cells emerge with shared bursting, splicing, and degradation kinetics [31]. The same mechanistic foundation can be generalized and coupled to a continuous mixture representation to model and infer continuous cell states [126] and trajectories [127].

Overall, even with simple systems such as the bursty transcription model for integrating nascent and mature RNA counts, we have demonstrated the potential for interesting discovery and generalization. Subtle differences and novel behaviors between cells in various states of development, treatment, and disease can be identified and ascribed to biophysical phenomena; biophysical hypotheses can be compared in their ability, or lack thereof, to capture important dataset features; data from multiple sequencing and imaging approaches can be described in a common framework; the balance of technical noise and biological variation can be described and investigated in a self-consistent fashion. In light of the ever-growing complexity and resolution of high-throughput genomics data, *Monod* demonstrates a framework poised to take advantage of other multimodal data types, and to parametrize and generate more complex mechanistic hypotheses.

## 4 Acknowledgments

We thank Sina Booeshaghi and Meichen Fang for useful discussions in the course of developing *Monod*. G.G. and L.P. were partially funded by NIH U19MH114830 and NIH 5UM1HG012077-02. M.C. is supported by the National Science Foundation Graduate Research Fellowship Program under grant no. 2139433. T.C. was supported by the Eric and Wendy Schmidt Center at the Broad Institute of MIT and Harvard. Part of this work was performed during G.G.’s Data Sciences Co-op with Celsius Therapeutics, Inc.

## 5 Author Contributions

All authors contributed extensively to the work presented in this paper. The first draft of the paper was conceptualized and written by G.G. and L.P. Section 2.1 was predominantly conceptualized by G.G., J.J.V., and L.P. Section 2.2 was predominantly conceptualized and executed by G.G., T.C., and L.P. Sections 2.3 and 2.4 were predominantly conceptualized and executed by M.C. and L.P. Section 2.5 was predominantly conceptualized and executed by G.G., J.J.V., M.C., and L.P. Section 2.6 was predominantly conceptualized and executed by G.G., J.J.V, and L.P.

## 7 Tables

N/A

## 9 Methods

### 9.1 Overview of *Monod* inference

Here we briefly describe how models are defined and inference performed in the *Monod* framework. Further details are available in the package API at https://monod-examples.readthedocs.io. The *Monod* package uses algorithms implemented in the *NumPy* [134], *SciPy* [135], and *numdifftools* [136] Python packages. Currently, all inference models require nascent and mature mRNA information, which can easily be obtained from realigning scRNA-seq datasets [32, 33]. For the datasets used in this study, unspliced and spliced (or nascent and mature mRNA) counts were obtained through *kallisto* | *bustools* 0.26.0 [32, 137]. We use the `kb ref` function with the `--lamanno` option to create exon- and intron-containing reference, and the `kb count` function with the `--lamanno` option which takes in the FASTQ inputs to produce counts from the reads pseudo-aligned to the reference. For the *kallisto* | *bustools* version 0.29.0 the `--workflow nac` option is used instead, and the mature and ambiguous counts summed, which produces the spliced matrices of the `--lamanno` option. The details of processing for each specific dataset are reported below. See Supplementary Figure 1 for a visual overview of the *Monod* inference procedure.

#### 9.1.1 Biophysical model definition

The biological models in *Monod*, i.e., the models describing the biophysical processes within the cell, all have the following generic structure:

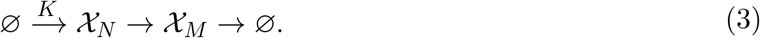

For simplicity, *𝒳*_*N*_ denotes nascent or unspliced mRNA species, whereas *𝒳*_*M*_ denotes mature or spliced mRNA species. These concepts are not strictly synonymous, for example, “nascent” species may refer to mRNA in the process of transcription. We therefore adopt this convention to emphasize that the models are generic, as long as a two-step process accurately describes the underlying physics (Section 2.6 of Carilli et al. [126]). *K* is stochastic process governing mRNA arrivals, the transition from *𝒳*_*N*_ to *𝒳*_*M*_ is isomerization or splicing, and the removal of *𝒳*_*M*_ is efflux or degradation. In this generic formulation, we do not specify the precise meaning of “→”, which may represent Markovian or deterministic transitions. To compute probabilities, we need to first evaluate the probability generating function (PGF):

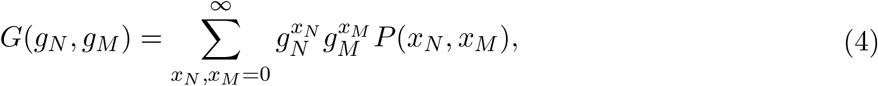

where *P* (*x*_*N*_, *x*_*M*_ ) is the joint probability mass function of the nascent and mature counts. We typically consider the log-PGF in terms of shifted complex variables *u*_*z*_ := *g*_*z*_ −1, where we introduce the generic transcript subscript *z* ∈ *{N, M}*.

To account for the imperfect sequencing process, we then define a set of technical noise models. All of the models assume that the reverse transcription, amplification, and identification of the *in vitro* mRNA pool is fundamentally an independent and identically distributed process across all cells and molecules of a single gene, although the identically distributed assumption is relaxed across genes. For inference, we define the PGF *H*, which is the function composition of *G* with the PGFs *G*_*z,t*_ of the sequencing processes:

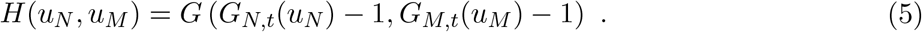

Here *t* denotes the PGF for “technical” sequencing processes. The governing rates of the biological and technical models constitute the inferred parameters *θ*. The full forms and derivations for each of the models implemented in *Monod* can be found in Section S1.

#### 9.1.2 Parameter inference procedure

To perform inference, *Monod* takes in the nascent and mature RNA molecule count matrices and a user-selected biological and technical noise model. By default, *Monod* filters out genes with very low observed average and maximum expression 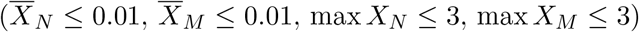, and those with excessively high observed maximum expression, which are too computationally intensive to fit (max *X*_*N*_ ≥ 400, max *X*_*M*_ ≥ 400). Genes are selected from the remaining genes for inference. The *Monod* inference procedure is then as follows:

1. Initialize a grid over possible values of technical sampling parameters (for the nascent and mature counts respectively), which spans several orders of magnitude to capture physically-realistic sampling rates of the transcriptome [138].

For each grid point, in parallel:

2. Initialize the biological parameters using the analytical moments of the desired model. These moments are derived and reported in Section S1.4.
3. Following the procedure outlined in [28, 37], evaluate the generating function, potentially using numerical quadrature, then invert it using a multivariate inverse fast Fourier transform to obtain the steady-state probability distribution. This evaluation may be performed in parallel for each gene.
4. Compute the Kullback-Leibler Divergence (KLD) between the empirical histogram and the analytically-derived likelihood, and add it across genes, to obtain the objective function.
5. Repeat steps 3 and 4, iteratively updating the biological parameters to minimize the total KLD. We use the *SciPy* `optimize.optimize` function [135] for optimization.

Once the optimization has been completed for each grid point,

6. Determine the optimal set of technical parameters as the grid point with the lowest total KLD summed over all genes. This produces a final, optimal tuple of technical sampling parameters and the associated, inferred biophysical parameters.

Runtime across models per technical sampling parameter gridpoint ranged from tens of seconds to a little over a minute for almost 3,000 genes (see Supplementary Table 9). Memory usage for the inference procedure was similar across datasets of varying numbers of cells, with a slight increase for an increased number of genes (see Supplementary Table 10). For example, memory usage for T cell sample from mouse intestines [60] for sample S0, which includes 536 cells, with 10 genes was 303.3 MiB and for 1000 genes was 343.7 MiB. Similarly, for sample S26 with 798 cells, the maximum memory usage was 311.8 MiB for 10 genes and 337.1 MiB for 1,000 genes. The inference procedure itself took little memory, with the main bottleneck being loading and storing input and inference result objects.

#### 9.1.3 Rejection statistics and parameter uncertainty

Once we have obtained parameter estimates for each gene, there are several metrics in *Monod* which are used to reject genes that demonstrate poor fits and/or model misspecification. Genes are rejected if (1) the Hellinger distances between the empirical histograms and the model predicted probabilities were above 0.05, and (2) the model was rejected by a chi-squared test with *p* = 0.05. We adjust the chi-squared threshold with a Bonferroni correction for the number of tested genes. The chi-squared test was implemented using the function `scipy.stats.mstats.chisquare` [135]. In addition, we exclude genes that are near the bounds of the search grid (within 1% of bounds, relative to the size of the search domain).

To compare biophysical hypotheses and fits for genes under different model specifications, the Akaike information criterion (AIC) is calculated per model (*i*) as follows:

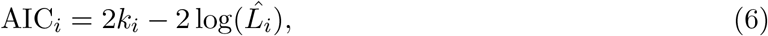

where *k*_*i*_ is the number of parameters in model *i* and 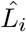 is the likelihood of the observed counts under model *i* at the optimal parameters.

Uncertainty estimates for the inferred parameters, or standard error (*σ*) values, are calculated from the square root of the diagonals of the inverse Fisher information matrix (FIM), and used for error bar calculation. The FIM is calculated as the Hessian matrix of the KLDs between the empirical histograms and the distributions induced by the final inferred parameters. Plots of the gene fits, KLDs, and gene rejection statistics are automatically generated for the user as shown in Supplementary Figures 1, 2 and 3.

### 9.2 Mechanistic differential expression

#### 9.2.1 Statistical testing

At the conclusion of the procedure in Section 9.1, we obtain a set of fits, parameter uncertainties, and goodness-of-fit (GoF) criteria under one or more models. We seek to move beyond averages and explain the differences between single-cell samples and cell types in terms of biophysical distribution parameters, in the spirit of Munsky et al. [139]. In this section, we propose and apply an approach for the identification of cell type differences which would be poorly detectable using standard average-based procedures, and demonstrate its performance using an experiment studying transcriptional noise amplification.

Given a set of *Monod* fits, we can identify differentially expressed (DE) genes at the level of kinetic parameters (Section S4.6 of Gorin and Pachter [24]). Briefly, we define DE genes as those genes displaying high-magnitude variation between conditions or cell populations, as contrasted with genes whose variation is consistent with Gaussian random noise [24].

In the *n* = 1 scenario, typical statistical tests cannot be used. To identify DE genes, we perform orthogonal distance regression (ODR) between the parameter values of the two conditions being compared (for each parameter *θ*). We then fit a normal distribution to the model residuals across genes. This distribution has a mean (putatively, a low-magnitude offset due to uncertainty in fitting the technical noise parameters) and a variance (putatively, the variance of residuals under the null hypothesis). Next, we identify genes with outlier residuals (high-magnitude variation), defined to have a *Z*-statistic that yields a *p <* 0.01. Outlier genes are removed, and the regression and residual fitting procedures repeated, with iterative removal of outlier genes, incrementally restricting the genes to those that appear to follow the null hypothesis. The final set of outlier genes constitutes the DE genes. This procedure is implemented in *Monod*, and may be used for coarse pairwise comparisons, but we do not use it in this study.

In the *n >* 1 scenario, we can use typical statistical tests to compare the parameters. We apply the *t*-test without the assumption of equal variances, and restrict the analysis to genes that were successfully fit (decreasing *n* for some genes). To avoid overrejecting based on the *t*-test, we first debias the parameter estimates by running the *n* = 1 ODR procedure on pairs of samples, then subtracting the mean of the inferred null normal distribution of residuals to heuristically account for uncertainty in the technical noise parameters, and encode the intuition that the residuals should be zero-centered.

In this study, we use the notation DE-*θ* to indicate that the parameter *θ* exhibited a Bonferroni-corrected *p*-value lower than 0.1 and mean log_2_ fold change higher than 1 (between the two cell types or conditions). We also compute the mean log_2_ fold difference in mature RNA averages between the cell types/conditions, to identify DE-*θ* genes with a magnitude lower than 1. In other words, we can extract genes that have detectably large differences in biophysical parameters, but do not, on average, exhibit large differences in mature RNA averages *µ*_*M*_ .

#### 9.2.2 Signatures of frequency modulation

In the bursty model of transcription (Section S1.1.1), we fit the rate parameters log_10_ *β* and log_10_ *γ*, setting the burst frequency *k* to unity. This is formally equivalent to fitting 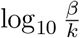 and 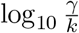 although the model’s transient dynamics depend on three independent parameters, the steady state behavior of the system is completely characterized by two parameter combinations. Hence, one cannot uniquely identify all three parameters given a single steady state dataset.

The models we present are not natively adapted to detect changes in *k*: to unambiguously distinguish between modulation of upstream and downstream processes, time-resolved data are mandatory. However, if the magnitudes of changes in 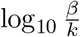 and 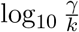 are approximately equal, frequency modulation is the more plausible scenario (see, e.g., Section S7.10.3 of Gorin and Pachter [24]).

We propose that the modulation of *k* can be motivated by biological argument. *β* and *γ*, the rates of splicing and degradation, use a one-step, first-order, memoryless reaction as a highly simplified representation of a series of chemical transformations effected in tandem with a spliceosome or a ribonuclease (RNase) complex respectively. However, spliceosomes and RNases are promiscuous, whereas transcription is highly regulated. Therefore, we hypothesize that targeted modulation of the burst frequency upstream at the gene locus is more mechanistically plausible than the synchronized and targeted modulation of the downstream processes.

If we *assume β* and *γ* are constant between conditions or cell types, we can compute an estimate of *k* modulation between population 1 and population 2:

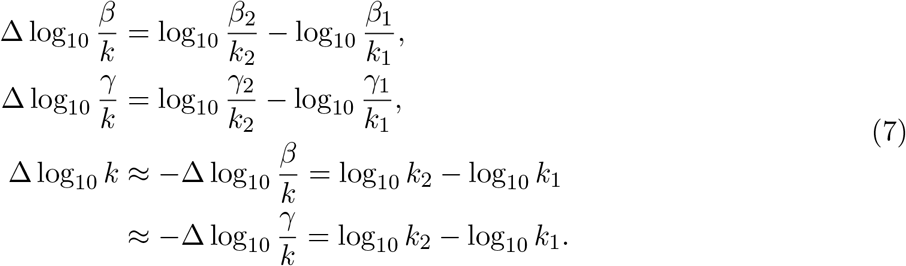

Therefore, if the approximate equality 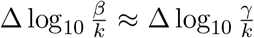 holds, we can propose that Δ log_10_ *k* has a similar magnitude, but the opposite sign. We average the two to estimate the burst frequency modulation:

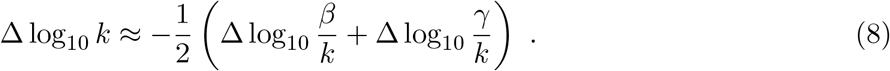

#### 9.2.3 Data processing: mouse neurons

To illustrate the approach, we compared the parameters for glutamatergic and GABAergic cell types from four mouse datasets (B08, C01, F08, and H12) generated by the Allen Institute for Brain Science [56, 140]. We previously performed the fits and identified genes that suggested substantial parameter modulation [24]. Here, we revisit the fits and summarize the key findings. We report the *t*-test results from these analyses in Supplementary Data Table 1.

We identified DE-*θ* genes as described above, for the parameter values between the GABAergic and glutamatergic cell types, where the mean log_2_ fold difference in mature RNA averages between the cell types also had magnitude lower than 1. We also omitted data points that were discarded by goodness-of-fit testing, as described in Section 9.1.3.

Next, we averaged the mean log_2_ fold changes in *β* and *γ* to obtain an estimate of the log_2_ fold change in the burst frequency, as in Equation 8. We plotted the resulting aggregated fold changes in burst size and burst frequency against each other, highlighting genes that were DE-*θ* for some biological *θ*, but not DE-*µ*_*M*_ .

We colored these genes by their effect on noise. It is elementary to show that, if the mean remains constant, a decrease in *b* compensated by an increase in *k* or equivalently a decrease in *β/k* and *γ/k*, leads the joint distribution of nascent and mature RNA to become bivariate Poisson. For example, if a gene was found to exhibit significantly higher *b, β*, or *γ* in GABAergic cells, we assigned it to the GABAergic set, as it suggests relative noise amplification in this cell type. This analysis is shown in Figure 1d.

The structure of this plot bears further discussion, as it provides a convenient summary of useful statistical properties. The solid diagonal line denotes the set of *b* and *k* combinations that yield a constant mean (all other parameters held equal). The dashed diagonal lines are offset by unity, and show the range of parameters that give averages with a lower than twofold change in the mean. The dashed vertical and horizontal lines correspond to no change in *b* and *k*, respectively. Qualitatively, moving toward the top right corresponds to increasing the mean; moving toward the bottom left corresponds to decreasing the mean; moving toward the top left corresponds to decreasing the noise to the Poisson limit; moving to the bottom right corresponds to increasing the noise.

To demonstrate the qualitative impact of noise modulation, we visualized the distributions, as well as the fits, in both cell types, based on data from the B08 dataset. We selected the genes *Nin* and *Bach2*, which are associated with neuronal development, as discussed in Gorin and Pachter [24].

In addition, we computed these genes’ mature count averages in each cell type. This demonstration is given in Figure 1d, right panels.

#### 9.2.4 Data processing: mouse embryonic stem cells

To demonstrate the potential of this approach for detecting broad trends in transcriptional modulation without replicates, we considered the transcriptomes of mouse embryonic stem cells with and without 5’-iodo-2’-deoxyuridine (IdU) perturbations. This dataset was generated by Desai et al. [47, 54] to investigate the effect of IdU incorporation on transcriptional bursting properties; the authors found that the perturbation appeared to increase the noise genome-wide, but did not affect averages.

##### Preprocessing

We used *kallisto* | *bustools* 0.26.0 to pre-process data (Section 9.1), using a pre-built *M. musculus* genome from https://support.10xgenomics.com/single-cell-gene-expression/software/downloads/latest (mm10, 2020-A version) to construct the reference. We obtained the raw FASTQ files for the DMSO (control) and IdU datasets, both of which were generated using the 10x v2 chemistry. From `kb count` (as described in Section 9.1), we obtained unspliced (intron-containing) and spliced (non-intron-containing) RNA count matrices [32, 141].

##### Filtering

We filtered the dataset to remove “low-quality” cells or empty droplets. First, we removed all barcodes that did not pass the default *kallisto* | *bustools* filter. Next, we removed all cell barcodes that were associated with fewer than 4 *×* 10^3^ molecular barcodes, computed over all genes, corresponding to standard knee plot filtering procedures.

Next, we used *Monod* 0.2.6.0 to extract genes with moderate to high expression. We removed genes based on the *Monod* filter described in Section 9.1.2. This procedure produced a set of 4,373 genes that met the thresholds in both cell populations. We randomly selected 2,000 genes for further analysis, ensuring that the genes analyzed in the original report (*Nanog, Sox2, Pou5f1, Klf4, Wdr83, Stx7, Hif1an, Mtpap, Farsa, Wipi2*, and *Snd1* ) were included.

##### Inference and analysis of biophysical parameters

To fit a mechanistic model, we used *Monod* 0.2.6.0. We set up a 20 *×* 21 grid over the {log_10_ *C*_*N*_, log_10_ *λ*_*M*_ } domain listed in Supplementary Table 7.

At each grid point, we iterated over the 2,000 genes, using gradient descent to identify the conditional maximum likelihood estimate of {log_10_ *b*, log_10_ *β*, log_10_ *γ}*, where the rates *β* and *γ* are defined in units of burst frequency *k*. We used the conditional method of moments estimate as the starting point and performed 15 steps of gradient descent. The procedure was parallelized over up to eighty processors (Intel Xeon Gold 6152, 2.10GHz). Runtimes varied between sixteen and seventeen minutes.

To identify the optimal sampling parameters, we identified the grid point with the lowest total KLD, computed over all genes. To ensure we obtained the true optima under the bursty model, we performed four rounds of fixed-point iteration. First, we rejected a subset of genes if they were detected by the chi-squared test with *p* = 0.01 with a Bonferroni correction, and their Hellinger distance from the data distribution exceeded 0.05. Next, we recalculated the optimum based on the remaining data, and repeated the procedure. This procedure did not change the optimum for any of the datasets. Further, we investigated the stability of the optima under gene subsampling, and found them to be stable and consistent.

The discovered optima were not consistent between datasets (orange points, Supplementary Figure 6), and the likelihood landscapes were rugged and inconclusive (hatched region, Supplementary Figure 6). This observation accords with our previous analyses of 10x v2 datasets (e.g., panels a. of figures in Section S7.6 of [24]): the older v2 technology does not appear to provide enough information to identify the technical noise parameters. Therefore, we somewhat arbitrarily used the grid point closest to log_10_ *C*_*N*_ = −6.5, log_10_ *λ*_*M*_ = −1.2, near the optimum discovered for a mouse neuron dataset in Figure 3e of [24]. We analyzed the datasets under that set of parameters, recomputing the goodness-of-fit statistics accordingly.

We computed the mean log_2_ fold change in burst size and burst frequency (Equation 8), and plotted them against each other, using the conventions in Figure 1d. We omitted data points that were discarded by goodness-of-fit testing. Finally, we identified all genes with log_2_ fold change higher than 1.5 in *b* as well as *k*, which demonstrated significant noise amplification. To focus on genes with biologically interesting effects, we selected only those which had a mature RNA mean greater than unity in at least one of the conditions, and reported them.

#### 9.2.5 Data processing: mouse germ cells

To assess the use of biophysical inference of the development timeline, we utilized the germ cells scRNA-seq collected in Mayère et al. [57] over several stages of mouse embryo development. In this study, germ cells were collected from male and female mouse embryos from the E11 stage to the E16 stage, with biological replicates per stage and sex.

##### Preprocessing

We utilized the unspliced and spliced count matrices provided in the original study [57].

##### Filtering

As described above, we filtered out “low-quality” cells or empty droplets, removing cell barcodes with fewer than 10^4^ molecular barcodes, computed over all genes, corresponding to standard knee plot filtering procedures. We then used *Monod* 0.2.6.0 to extract 3,000 genes with moderate to high expression (Section 9.1.2), and included genes discussed in the study [57] for analysis. Each embryonic stage and biological replicate was treated separately, as samples were not necessarily sequenced together. We used samples/stages with at least 50 cells.

##### Inference and analysis of biophysical parameters

To fit a mechanistic model, we used *Monod* 0.2.6.0. We set up a 10 *×* 11 grid over the {log_10_ *C*_*N*_, log_10_ *λ*_*M*_ } domain listed in Supplementary Table 7. Optimal technical parameters, as described above, were fit for *each* germ cell sample.

We then rejected a subset of genes if they were detected by the chi-squared test with *p* = 0.01 with a Bonferroni correction, and their Hellinger distance from the data distribution exceeded 0.05 (i.e., genes with poor fits to the bursty transcription and sequencing model). Next, we recalculated the optimum based on the remaining data, and repeated the procedure.

To make all inferred parameters comparable, we also perform “debiasing,” subtracting the constant term from the orthogonal distance regression of the parameters of each stage versus the earliest (E11) stage for comparison, as discussed in Section 9.2.1. We obtain parameter uncertainties by calculating approximate 99% confidence intervals (C.I.s), i.e., 2.756 *σ*, using the *σ* estimates from the Fisher information matrix (Section 9.1.3), and display them as error bars in the figures.

Inferred parameters and trends were compared to previous analyses in the original study, which utilized read-depth normalized and log-transformed expression to analyze *Tbrg4* trends and unspliced/spliced ratios [57].

#### 9.2.6 Data processing: PDAC tumor samples

Biophysical inference was performed on malignant cells from pancreatic ductal adenocarcinoma (PDAC) residual tumors in a patient with FOLFIRINOX and fluorouracil (5-FU)/capecitabine (CRT) treatment, a patient with CRT and losartan (CRTl) treatment, and three untreated patients. We only used 10x v3 snRNA-seq samples from the original study.

##### Preprocessing

Nascent and mature (unspliced and spliced) count matrices were generated using The *kallisto* | *bustools* 0.29.0 with the `--workflow nac` option. The mature and ambiguous counts were summed so as to be equivalent to the spliced matrices from the earlier `--lamanno` workflow.

##### Filtering

Cells were filtered for cell barcodes in the original study that denoted malignant cells [61]. We then used *Monod* 0.2.6.0 to extract 3,000 genes with moderate to high expression (Section 9.1.2), and included genes discussed in the study [61] for analysis. Each patient sample was treated separately, as samples were not necessarily sequenced together. We used samples that were run with the 10x v3 technology, with at least 50 annotated malignant cells.

##### Inference and analysis of biophysical parameters

Using *Monod* 0.2.6.0, we set up a 5 *×* 6 grid over the {log_10_ *C*_*N*_, log_10_ *λ*_*M*_ } domain listed in Supplementary Table 7. Optimal technical parameters were determined for *each* patient sample of malignant cells.

Genes were then rejected if they were detected by the chi-squared test with *p* = 0.01 with a Bonferroni correction, and their Hellinger distance from the data distribution exceeded 0.05 (i.e., genes with poor fits to the bursty transcription and sequencing model). We then recalculated the optimum based on the remaining data, and repeated the rejection procedure. In total, 1,030 genes were not rejected across all samples and used for all downstream analysis.

As with the germ cell data above, we also performed “debiasing,” subtracting the constant term from the orthogonal distance regression of the parameters of each stage versus an untreated patient sample for comparison (see Section 9.2.1). Fold changes (FCs) between samples were then calculated for the corrected parameters of the CRT patient, and the CRTl patient, versus all three untreated samples (thus n=3 FC ‘replicates’).

To identify genes where transcription (burst size *b* or burst frequency *k*) was altered (abs(log_2_ FC) *>* 1) consistently across all three comparisons, we denoted genes where *β/k* (splicing) and *γ/k* (degradation) FCs were of the same sign and similar magnitude (with a ratio of 1.5 or less) as likely denoting burst frequency changes. Otherwise, for genes with consistent changes (across all three replicates) in splicing or degradation that did not show near identical FC behavior, the splicing and degradation rate FCs were treated independently. To compare fold changes in mature RNA counts between samples, we summed counts across cells to get pseudobulk expression values per sample, then normalized these values as in DESeq2 [51]. Specifically, this is done by creating a reference sample by taking the geometric mean of mature gene counts per gene across all samples, dividing the value per gene per sample by the corresponding value per gene in the reference sample, then finding the median of these ratios as the size-factor per sample. Samples are finally normalized by dividing by the size-factor.

#### 9.2.7 Data processing: Irradiated T cells

We applied Monod to T cells from the jejunum of mouse intestines following radiation-induced intestinal injury (RIII) [60] to infer biophysical parameters across the timecourse of recovery. The original study aimed to characterize the transcriptomic response of the intestinal microenvironment to RIII by treating WT C57BL/6 J male mice with 15 Gy of abdominal radiation. Organs from mice pre-radiation treatment (day 0) and several days following treatment (days 1, 3, 7, and 14) were collected and single cells were extracted and sequenced (10x Genomics v2 chemistry).

##### Preprocessing

The *kallisto* | *bustools* version 0.29.0 the `--workflow nac` option was used to generate the nascent and mature counts matrices, summing the ambiguous and mature counts matrices (which is equivalent to the ‘spliced’ count matrix of the earlier `--lamanno` workflow).

##### Filtering and inference

Cells were first subset to those given annotations in the original dataset [60]. Choosing to focus on the immune response to radiation treatment, we selected T cells as the target cell type, then subset to samples with more than 100 T cells/sample. This left 11 samples across four time points (samples S0, S22, S23, and S26 from day 0; S2 and S32 from day 1; S10, S8, and S9 from day 3; and S38 and S40 from day 14).

For these 11 samples, we fit the bursty model with Poisson sampling noise using *Monod* 0.2.6.0 as described in Section 9.1.2 over a 10 by 11 grid of {log_10_ *C*_*N*_, log_10_ *λ*_*M*_ } within the ranges listed in Supplementary Table 7. We fit the same 3,000 genes across all samples, then performed goodness of fit testing at a manual set technical sampling parameter grid point (-6.78, -1.25). We rejected genes using a chi-squared cutoff *p* = 0.01 with a Bonferroni correction per sample and if their Hellinger distance from the empirical count distribution was greater than 0.05 in any sample, leaving 1,182 final genes for analysis.

##### Analysis

Parameters were “debiased” to remove sample specific effects by regressing all genes that passed QC per sample against S38 (as it had the most genes that were not rejected following GoF testing) and subtracting the mean of the orthogonal distance regression, as described in Section 9.2.1.

To find statistically significant differences in parameters, we performed for each day (days 1, 3, and 14 post radiation) against day 0 (control day) a *t* -test to the “debiased” log_10_ of inferred burst sizes, relative splicing rates, and relative degradation rates. We used `scipy.stats.ttest_ind` with unequal variances [135]. To comparably find statistically significant differences in mature mean expression values, we first added counts per cell to obtain “pseudo-bulked” expression values per sample, then normalized these values as in DESeq2 [51]. Specifically, this is done by creating a reference sample by taking the geometric mean of mature gene counts per gene across all samples, dividing the value per gene per sample by the corresponding value per gene in the reference sample, then finding the median of these ratios as the size-factor per sample. Samples are finally normalized by dividing by the size-factor. We performed a *t* -test with unequal variance per day against day 0 on the log_10_ of these normalized mature counts. We corrected the raw p-values per parameter using a Bonferroni correction for the number of tested genes (1,182) to find false discovery rates (FDR). We flagged as significant all parameters with a false discovery rate of less than or equal to 0.05.

We further calculated the average fold change between days 1, 3, and 14 and day 0 (separately per day) by averaging the raw values per day and taking the ratio against the average of the raw values on day 0.

To identify genes with similar behavior across time points, we formed vectors per gene with parameter *and* mature expression FCs against day 0 for all days and took the Euclidean distance against the FC vector for a given target gene. This enables discovery of genes with similar parameter changes across all days post radiation that further result in similar expression changes.

### 9.3 Biophysical model selection and rejection

To compare biophysical hypotheses across genes and cell types, we fit five available transcriptional models with *Monod* (models defined in Section S1.1), using scRNA-seq data for the mouse brain sample B08, originally generated by the Allen Institute for Brain Science [56].

#### Preprocessing

We used *kallisto* | *bustools* 0.26.0 to pre-process data as described in Section 9.1, using a pre-built *M. musculus* genome from: https://support.10xgenomics.com/single-cell-gene-expression/software/downloads/latest (mm10, 2020-A version). We obtained the raw FASTQ files for the dataset B08 [56, 140], which was generated using the 10x v3 single-cell chemistry. Next, we used the `kb count` function as described in Section 9.1, to produce unspliced (intron-containing) and spliced (non-intron-containing) RNA count matrices [32, 141].

#### Filtering

We filtered the dataset to remove “low-quality” cells or empty droplets. First, we removed all barcodes that did not pass the default *kallisto* | *bustools* filter. Next, we removed all cell barcodes that were associated with fewer than 10^4^ molecular barcodes, computed over all genes, corresponding to standard knee plot filtering procedures.

Next, we split the dataset by cell type and subtype annotations [56]. We extracted seven classes: six glutamatergic subtypes (L2/3 IT, L5 IT, L6 IT, L5/6 NP, L6 CT, and L6b) and their union (“glutamatergic”). We omitted low-abundance cell types (L6 IT Car3 and L5 ET, with fewer than ten barcodes) from analysis and inclusion in the “glutamatergic” category. We then used *Monod* 0.2.6.0 to extract genes with moderate to high expression given the default filter (Section 9.1.2).

#### Inference of biophysical parameters for five models

For each of the 2,951 genes that passed previously described *Monod* 0.2.6.0 expression thresholds (Section 9.1.2) and each of six glutamatergic cell subtypes (L2/3 IT, L5 IT, L6 IT, L5/6 NP, L6 CT, and L6b), we fit five biological models using *Monod* 0.2.6.0. In particular, we used model arguments to fit biological models of bursty transcription (“Bursty”), constitutive transcription (“Constitutive”), constitutive transcription with extrinsic noise (“Extrinsic”), bursty transcription with delayed degradation (“Delay”), and bursty transcription with delayed splicing (“DelayedSplicing”). Models are described in Section S1. For each model, we also included Poisson technical noise with length bias for nascent/unspliced molecules and Poisson noise without length bias for spliced molecules, setting up a 10 *×* 11 grid over the {log_10_ *C*_*N*_, log_10_ *λ*_*M*_ } domain listed in Supplementary Table 7.

Optimal technical parameters per dataset and cell type were discovered by iterating over all genes at each grid point, identifying the conditional maximum likelihood estimates of the biological model’s parameters (using gradient descent with a maximum of 15 steps), then identifying the grid point with the lowest KLD computed over all genes. Biological parameters were initialized at the conditional method of moments estimate. Two rounds of this fixed-point iteration were performed to increase confidence in the achieved optima.

#### Rejection criteria and AIC weights

As described in Section 9.1.3, gene fits under each model are rejected based on Hellinger distance and the chi-squared test *p*-value, between the empirical histograms and fit distributions. In addition, we rejected genes that were too near the parameter domain bounds. The AIC metric, described in Section 9.1.3, was also used to compare multiple model fits for a given gene. In particular we use AIC (or Akaike) weights to make model comparisons, where:

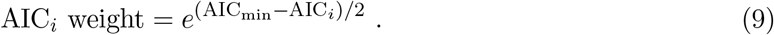

AIC weights were normalized to sum to one, corresponding to the interpretation of these values as model posteriors [142]. Genes were assigned to the model with the largest normalized AIC weight.

### 9.4 Principled integration of single-cell technologies

In this section, we describe a fairly general theoretical view of integrating data from different modalities, as well as the specific approaches we use to treat single-nucleus and seqFISH+ data.

#### 9.4.1 Bayes’ rule and integration

##### Warm-up: single-modality case

To begin, consider the single-modality case: let ***x***_*i*_ denote the vector of observed counts after sequencing cell *i* (out of *n*_*c*_ total cells), and *p*(***x***_*i*_|***θ***) denote the corresponding parameterized likelihood. By Bayes’ rule, we have

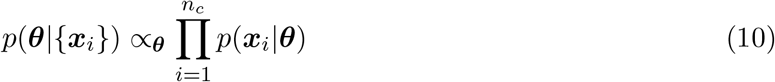

assuming each cell is independent, and that we have a uniform prior on ***θ***. Especially when *n*_*c*_ is large, the posterior is generally tight around the maximum likelihood estimate (MLE) ***θ***^∗^; in particular, we can expand the likelihood as

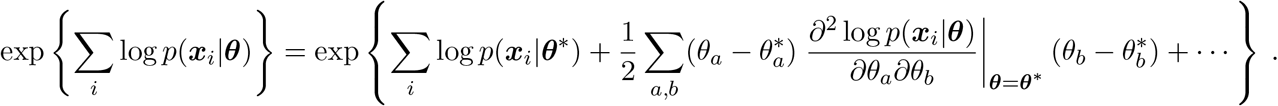

Assuming *n*_*c*_ is sufficiently large, by the central limit theorem, we have [143]

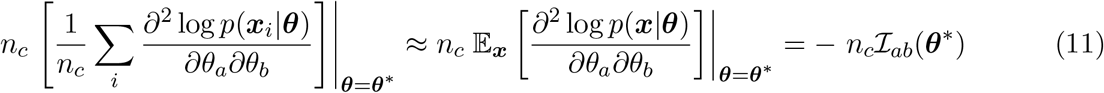

where we used the definition of the Fisher information matrix (FIM) in the last step. We now have

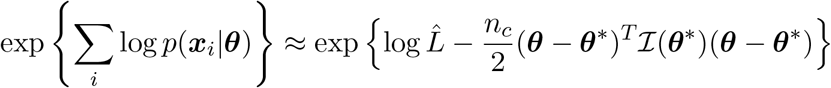

where 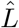 denotes the likelihood at ***θ*** = ***θ***^∗^. Hence, when *n*_*c*_ is sufficiently large,

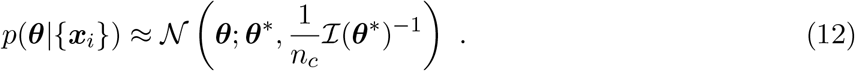

##### Multi-modality case

Assume instead that we have access to data from multiple modalities. Since the math looks essentially the same regardless of the number of modalities we use, we will assume two here for simplicity. Let 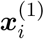 denote the vector of observed counts after sequencing cell *I* (out of 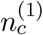 total cells) via modality 1, and 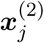 the analogous vector after sequencing cell *j* (out of 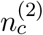 total cells) via modality 2. We assume here that data from different modalities is independent, since this is true in practice. By Bayes’ rule, if we once again assume a uniform prior we have

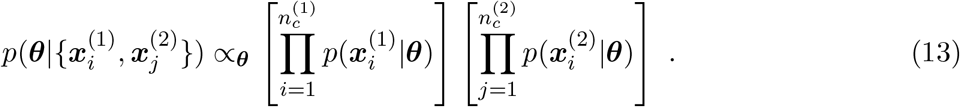

Assuming both 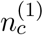 and 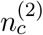 are sufficiently large, by the previous argument we approximately have

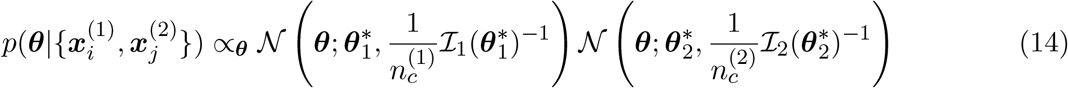

where 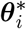 is the MLE obtained by using only data from modality *i*, and 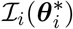 is the corresponding FIM. The posterior associated with a product of normally-distributed likelihoods is well-studied in the context of Bayesian cue combination; the result is 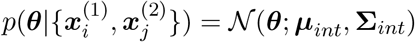, where

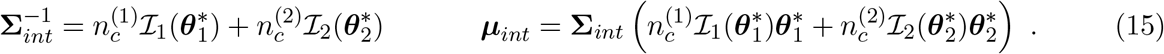

If we additionally assume that the off-diagonal terms of the FIM (which control correlations between estimates of different parameters) are negligible, the above expressions simplify somewhat. For any given parameter *θ*_*k*_, we have

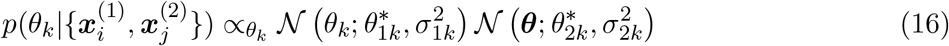

where 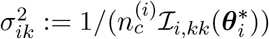. This implies 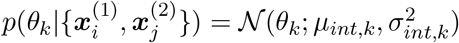, where

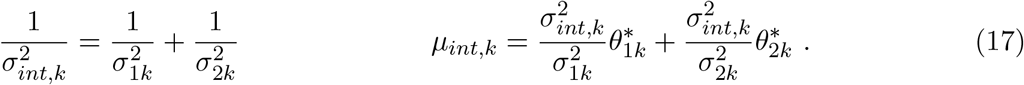

We use the above equations to determine integrated estimates and the associated uncertainties in Section 2.5 and Figure 5.

##### Caveat about uncertainty in technical noise parameters

In practice, technical noise parameters appear to be much harder to infer than biology-related parameters (e.g., burst size). Partly due to this issue, *Monod* treats these two types of parameters differently in its optimization process, and only quantifies uncertainty about biology-related parameters. Technical noise parameters are optimized via grid search (*C*_*N*_ by *λ*_*M*_ ), and for each grid point the other parameters are optimized via gradient descent. At the highest-likelihood technical noise grid point and MLE, uncertainty in biology-related parameters is quantified by numerically computing the Hessian.

In the context of the preceding derivation, we essentially assume that uncertainty about technical noise parameters is not the dominant contributor to uncertainty in other parameters, and so can be neglected. One justification for this assumption is that the technical noise parameters *C*_*N*_ and *λ*_*M*_ are assumed to be the same for each gene, and hence much more data is used to infer them than the biology-related parameters (which only use data from individual genes). From another perspective, this approach is consistent with estimating, e.g., “size factors” for cells and samples [51, 144] from raw data, then fitting models conditional on these estimates, and eliding any associated uncertainty.

#### 9.4.2 Nuclear data integration

We would like to coherently integrate single-cell and single-nucleus RNA sequencing data. To do so, we need to specify the relationship between the two modalities. We can establish such a relationship from first principles by making assumptions regarding the underlying biophysical processes. For example, by proposing that nascent RNA are restricted to the nucleus, we can reasonably assume that the nascent RNA dynamics should be identical between the two modalities. On the other hand, the mature RNA distributions and dynamics may have substantial differences, as nuclei are depleted in this species relative to the entire cell. In this section, we propose a possible foundation for the integration of these modalities.

To compare the distributions of single-cell and single-nucleus datasets and explain them using a mechanistic argument, we used mouse neuron datasets generated by 10x Genomics.

##### Mechanistic model definition

To describe the stochastic dynamics and sampling in a single-cell dataset, we use the formulation given in Section 9.1.1 and outlined in more detail in Gorin and Pachter [24]. To connect this model to nuclear data, we note that formally, it can arise from the following model:

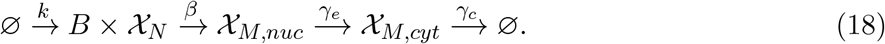

where *𝒳*_*M,nuc*_ and *𝒳*_*M,cyt*_ are nuclear and cytoplasmic mature RNA species, respectively. The rate *γ*_*e*_ describes the efflux of nuclear RNA, whereas the rate *γ*_*c*_ describes the degradation of cytoplasmic RNA.

In the limit *γ*_*c*_ ≪ *γ*_*e*_ or *γ*_*c*_ ≫ *γ*_*e*_, the model in Section 9.1.1 approximately holds for cytoplasmic data: if one of these stages is considerably longer-lived, the two-stage processing of mature RNA can be effectively described by a one-stage model. In this case, *γ* can be interpreted as the lower rate. We typically assume that the first limit is most relevant, although orthogonal data suggest that the details are highly gene- and tissue-dependent [145]. We note that it is, in principle, straightforward [119] to implement a model that explicitly incorporates both parameters; however, for computational facility, we use the simpler reduced model and discard genes that fail to fit it. On the other hand, for nuclear data, the model holds for *γ* = *γ*_*e*_.

##### Preprocessing

We used *kallisto* | *bustools* 0.26.0 to pre-process data. We downloaded a pre-built *M. musculus* genome from https://support.10xgenomics.com/single-cell-gene-expression/software/downloads/latest (mm10, 2020-A version). To build intronic and exonic references, we used the `kb ref` function with the `--lamanno` option. We obtained the raw FASTQ files for the “Brain 4” and “Brain Nuclei 4” datasets from two multiplexing experiments, both generated using the 10x v3 chemistry. We selected these datasets because they had the highest average molecule counts per cell in both technologies. Next, we used the `kb count` function with the `--lamanno` option, as well as the `-x 10xv3` whitelist option to quantify the datasets, producing unspliced (intron-containing) and spliced (non-intron-containing) RNA count matrices [32, 141].

##### Filtering

We filtered the dataset to remove “low-quality” cells or empty droplets. First, we removed all barcodes that did not pass the default *kallisto* | *bustools* filter. Next, we removed all cell barcodes that were associated with fewer than *T* molecular barcodes (*T* = 3 *×* 10^3^ for sc and 6 *×* 10^3^ for sn), computed over all genes, corresponding to standard knee plot filtering procedures. In addition, we removed cells with more than 10^5^ barcodes, as they may reflect obscure technical noise sources unique to single-nucleus data.

Next, we used *Monod* 0.2.6.0 to extract genes with moderate to high expression (Section 9.1.2). This procedure produced a set of 5,690 genes that met the thresholds in all of the cell populations. We randomly selected 2,000 genes for further analysis.

##### Inference and analysis of biophysical parameters

To fit a mechanistic model, we used *Monod* 0.2.6.0. We set up a 20 *×* 21 grid over the {log_10_ *C*_*N*_, log_10_ *λ*_*M*_ } domain listed in Supplementary Table 7.

At each grid point, we iterated over the 2,000 genes, using gradient descent to identify the conditional maximum likelihood estimate of {log_10_ *b*, log_10_ *β*, log_10_ *γ}*, where the rates *β* and *γ* are defined in units of burst frequency *k*. We used the conditional method of moments estimate as the starting point and performed 15 steps of gradient descent. The procedure was parallelized over up to fifteen processors (Intel Xeon Gold 6152, 2.10GHz). Runtimes varied between 2.2 hours for the whole-cell dataset and 3.8 hours for the nuclear dataset.

To identify the optimal sampling parameters, we identified the grid point with the lowest total Kullback-Leibler divergence, computed over all genes. To ensure we obtained the true optima under the bursty model, we performed four rounds of fixed-point iteration. First, we rejected a subset of genes if they were detected by the chi-squared test with *p* = 0.01 with a Bonferroni correction, and their Hellinger distance from the data distribution exceeded 0.05. Next, we recalculated the optimum based on the remaining data, and repeated the procedure. This procedure did not change the optimum for any of the datasets. Further, we investigated the stability of the optima under gene subsampling, and found them to be stable and consistent.

The optimum discovered for the single-nucleus dataset demonstrated noticeably higher molecule observation probabilities (orange points, Supplementary Figure 6). This observation was supported by basic observations of the dataset statistics: despite the depletion of cytoplasmic RNA, the single-nucleus dataset had as much mature RNA as the single-cell dataset, and approximately half an order of magnitude more nascent RNA (Supplementary Figure 8, left). To illustrate these trends, we computed the offset from the ratio of the dataset-wide means. In addition, the single-nucleus dataset appeared to exhibit lower noise levels (Supplementary Figure 8, right).

From physical considerations, the two independent experiments, performed using different technologies, may not necessarily have the same sampling parameters. However, as the samples were taken from the same tissue, they should have the same physics of transcription and splicing. Therefore, we somewhat arbitrarily assumed that the single-cell optimum was sufficiently accurate, and chose a set of single-nucleus sampling parameters that provided the lowest squared errors for the log_10_ *b* and log_10_ *β* parameters. As shown by the blue points in Supplementary Figure 6, the optimum so discovered lay approximately half an order of magnitude above the optimum for the single-cell data, and within the top 5th percentile for the sampling parameter likelihood landscape (hatched region). We analyzed the single-nucleus data under that set of parameters, recomputing the goodness-of-fit statistics accordingly.

To illustrate the differences between the datasets, we plotted the inferred parameters and the identity line. To quantify uncertainty in the parameters, we exploited the Fisher information matrix as described in Section S4.3.4 of Gorin and Pachter [24]; we visualized the error bars, which represent the 99% confidence intervals for the biological parameters, conditional on the sampling parameter values. Finally, we applied a *t*-test, implemented through `scipy.stats.ttest_ind` [135], to the pairs of single-cell and single-nucleus parameter estimates. We omitted genes rejected by goodness-of-fit procedures from these computations and visualizations.

#### 9.4.3 Fluorescence data integration

Recent studies have quantified the number of intron-containing RNA molecules and mRNA molecules for thousands of genes in single cells in a spatially resolved manner using a series of fluorescent probes designed that hybridize to either intron-containing RNA transcripts or spliced mRNA transcripts [110, 111]. These studies report burst sizes for genes by averaging over intron-containing fluorescent spot intensities. However, comparing single-cell fluorescent reported burst sizes to scRNA-seq count data is not immediately possible: fitting scRNA-seq data for burst sizes using *Monod* allows principled comparison of these two single cell technologies.

We fit the bursty model of transcription to 10x v3 scRNA-seq data from mouse embryonic stem cells (mESCs) [112] using *Monod* to find burst sizes per gene, then compared the inferred burst sizes to those obtained using seqFISH+ on mESCs.

##### Preprocessing: seqFISH+ in mESCs

We downloaded the reported fluorescence intensity values associated with intron-containing and mRNA molecule spots in mESCs from Takei et al. [111] for two biological replicates. For the two replicates, we found burst sizes per gene as the authors reported by averaging the intron-containing spot intensities over all cells for a given gene.

##### Preprocessing: scRNA-seq in mESCs

We obtained raw 10x v3 FASTQ files from Khateb et al. [112] using the Sequence Read Archive run accession number SRR18364193. We then quantified the dataset using *kallisto* | *bustools* 0.26.0 with a pre-built *M. musculus* genome from https://support.10xgenomics.com/single-cell-gene-expression/software/downloads/latest (mm10, 2020-A version). Specifically, we called `kb count` with the `--lamanno` workflow option and `-x 10xv3` whitelist option to produce unspliced (intron-containing) and spliced (non-intron-containing) RNA count matrices [32, 141].

##### Filtering

To remove low-quality cells, we filtered the data by first discarding all barcodes that did not pass the default *kallisto* | *bustools* filter. We then also removed cells with fewer than 700 total unique molecule identifiers (UMIs), leaving 12,167 high-quality cells.

We then subset to genes with moderate to high expression using *Monod* 0.2.6.0 (Section 9.1.2). From genes that passed these thresholds, we randomly selected 2,000 genes to fit.

##### Inference and analysis of biophysical parameters

We used *Monod* 0.2.6.0 to fit the bursty model of transcription (Section S1.1.1) with Poisson technical noise (with the default length bias capture for nascent molecules). We set up a 20 *×* 21 grid over the {log_10_ *C*_*N*_, log_10_ *λ*_*M*_ } domain listed in Supplementary Table 7.

At each grid point, we iterated over the 2,000 genes, using gradient descent to identify the conditional maximum likelihood estimate of {log_10_ *b*, log_10_ *β*, log_10_ *γ}*, where the rates *β* and *γ* are defined in units of burst frequency *k*. We used the conditional method of moments estimate as the starting point and a maximum of 20 steps of gradient descent.

We identified optimal sampling parameters by finding the grid point with the lowest total Kullback-Leibler divergence between predicted fits and empirical distributions, computed over all genes. Finally, applied model selection criteria to reject poor fits by enforcing they pass a chi-squared test with *p* = 0.01 (corrected for 2,000 tested hypotheses with Bonferroni correction), and that their Hellinger distance from the data distribution exceeded 0.05.

### 9.5 Noise decomposition for assessing loss of signal after pre-processing

In order to characterize the performance of typical data processing and dimensionality reduction techniques we need a meaningful baseline for “good” performance. In this section, we derive a non-trivial decomposition of the coefficient of variation (CV) matrix, whose diagonal entries correspond to the squared coefficients of variation of nascent and mature RNA. Under the assumption that standard transformations act as “denoisers,” i.e., that they remove technical noise but not biologically meaningful variation (e.g., due to differences between cell types), we show that transformations imply a noise decomposition generally different from one obtained by fitting a mechanistic model; the extent to which the two differ can be used to assess a loss of biological signal. We also derive more generic bounds on the post-transformation CV matrix. For simplicity, we assume throughout that a pre-annotated dataset from a single tissue is available, and view the results of our noise decomposition analysis as conditional on the assumption that the cell type annotations are sufficiently accurate.

#### 9.5.1 Generative model and general noise decomposition derivation

Let *C* denote cell type, ***z*** denote the vector of true molecule counts, and ***x*** denote the vector of observed molecule counts. Assume the joint probability mass function has the form

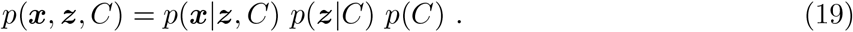

In words: assume the distribution of intracellular molecule counts is generally cell-type-dependent, and that technical noise can depend on both the true number of molecules and cell type.

The noise decomposition we exploit in the main text is a straightforward consequence of the law of total covariance. In particular, two separate applications of it produce the equations

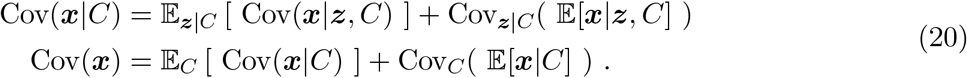

Combining these results, we obtain an interesting decomposition of the observed covariance matrix:

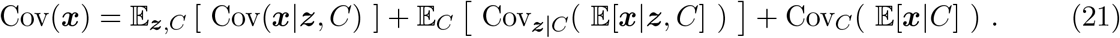

Overall observed variation is due to some combination of (i) technical noise, (ii) intracellular noise, and (iii) cell type heterogeneity; the above decomposition makes this statement quantitatively precise, since the first term can be identified with the contribution of technical noise, the second with the contribution of intracellular noise, and the third with the contribution of cell type heterogeneity. If there is no variation of some kind, the relevant term in the above decomposition is zero.

This decomposition can be converted into a decomposition of the coefficient of variation (CV) matrix by pre- and post-multiplying by the inverse of a diagonal matrix 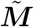 whose entries are the components of the observed mean 𝔼[***x***]. The observed CV matrix 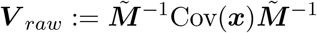 equals

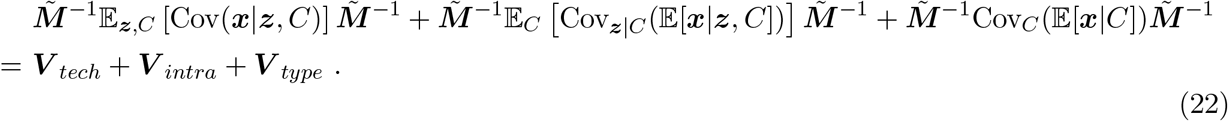

By definition, each term in the above decomposition is a positive semidefinite matrix, which allows us to interpret each term as capturing a “fraction” between zero and one of the overall variation.

In particular, if we assume ***x*** = (*x*_*N*_, *x*_*M*_ )^*T*^ is a two-dimensional vector that measures nascent and mature RNA counts, variation in nascent and mature counts can be decomposed as

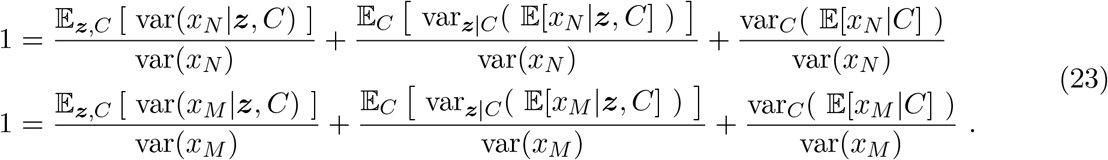

The above can be obtained by pre- and post-multiplying the (now assumed 2 *×* 2) matrix Cov(***x***) by the unit vectors (1, 0)^*T*^ and (0, 1)^*T*^, respectively. An analogous decomposition of variation in nascent-mature correlations can be obtained by pre- and post-multiplying by 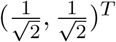 :

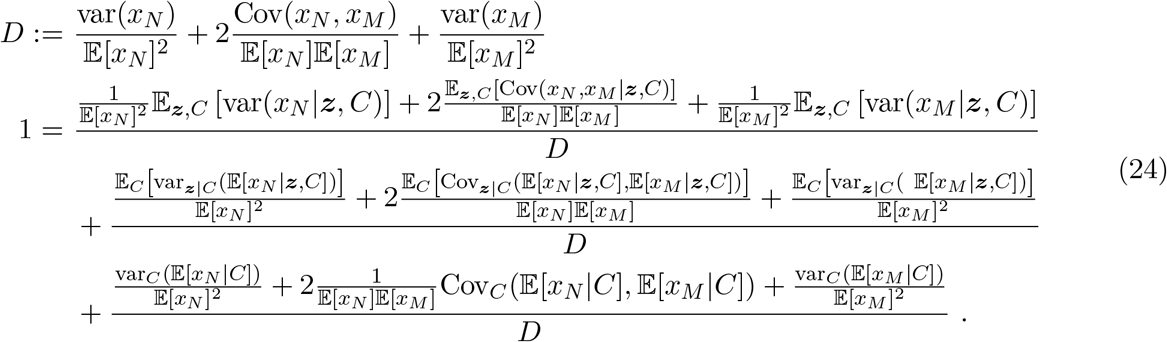

#### 9.5.2 Specializing the noise decomposition

The noise decomposition we just derived is fairly general, and we would like to incorporate some of the assumptions used by currently-supported *Monod* models. These assumptions are as follows:

- We assume each molecular species is observable.
- We assume each molecule is sequenced separately (in probability theory language: that the distribution *p*(***x***|***z***, *C*) is *infinitely divisible*).
- We assume the average effect of technical noise on the true molecule count ***z*** does not depend on cell type, i.e., that 𝔼 [***x***|***z***, *C*] = 𝔼 [***x***|***z***].

The first assumption implies ***x*** and ***z*** are vectors of the same length. The second implies

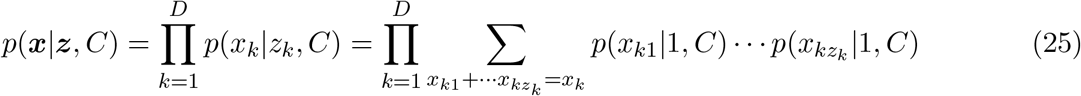

where *p*(*x*_*ki*_|1, *C*) is the same for all *i* = 1, …, *z*_*k*_. Our first two assumptions together imply

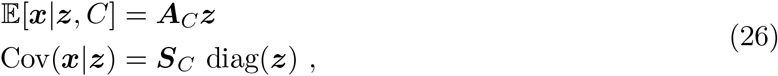

where ***A***_*C*_ and ***S***_*C*_ are (generally cell-type-specific) diagonal matrices whose diagonal entries are positive, and diag(***z***) is the diagonal matrix formed from the components of ***z***. These moments imply that our covariance decomposition becomes

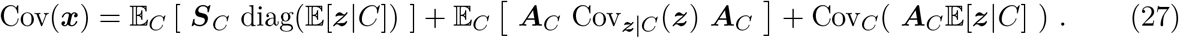

Our last assumption implies that ***A***_*C*_ = ***A***, i.e., that it does not depend on cell type. We now have

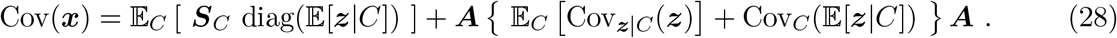

The equivalent CV matrix statement is that 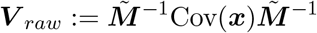 equals

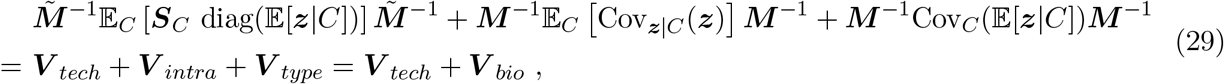

where ***M*** denotes the diagonal matrix whose entries are the components of 𝔼 [***z***]. A notable feature of this decomposition is that the sum of the latter two terms is precisely the CV matrix associated with ***z*** rather than ***x***, i.e., the analogous system without technical noise. Hence, there is a clean separation between “technical” (the first term) and “biological” variation (the last two).

If one assumes the catalysis technical noise model, as we do in the main text, ***S***_*C*_ = ***A***, and the expression slightly simplifies to

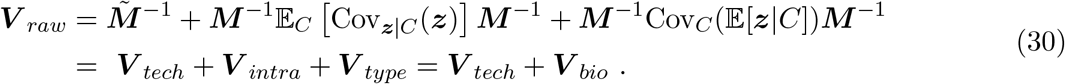

Equation 30 is the decomposition we exploit in the main text.

#### 9.5.3 Measuring noise fractions

Here, we describe two approaches to estimating noise fractions given single-cell data. The first uses fits to a mechanistic model (e.g., produced by *Monod* ), while the second assumes that the role of pre-processing steps (e.g., log1p) is to remove technical noise.

##### Model-based approach

Given fits to a (tractable) mechanistic model, one can use analytic expressions for the relevant means and covariances to determine each noise fraction. In Section 2.6, we fit a model that assumes bursty transcription and catalysis technical noise, so we will illustrate the approach in terms of that model’s moments.

Assuming bursty transcription, the relevant low-order moments of *p*(***z***|*C*) are

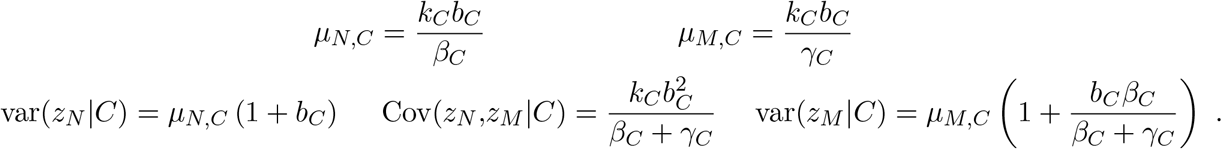

Since not all rates are identifiable from steady state data, *Monod* model fits assume *k*_*C*_ = 1, and we do the same in what follows. The relevant low-order moments of *p*(***x***|***z***, *C*) are

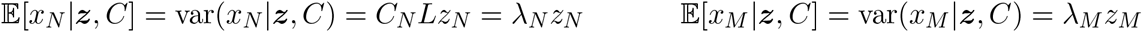

where *L* denotes gene length, and where we define *λ*_*N*_ := *C*_*N*_ *L* to ease notation. We can now write the noise decomposition in terms of model parameters, since we have expressions for all of the relevant moments. For example, the nascent squared CV and noise fractions are

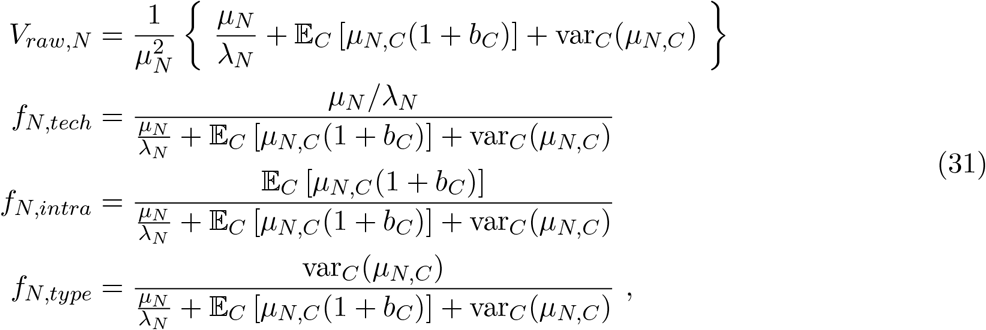

where *µ*_*N*_ denotes 𝔼 [*z*_*N*_ ].

In order to compute the expectations over cell types, we use the observed cell type ratios, and assume they are sufficiently close to the “true” ratios. From a “forward” model construction perspective, we suppose that the sequencing process uniformly samples cells from the underlying population. Naturally, this may not be the case, and physiological considerations may dictate otherwise: for instance, some cells may not be amenable to sequencing due to morphology incompatible with dissociation and capture [146]. In other words, the true cell population distribution may well be unknowable, barring, e.g., carefully calibrated orthogonal bulk RNA-seq experiments [147]. From the considerably more fruitful “reverse” data-analytic perspective, this condition is guaranteed by construction: we *define* the entries of the categorical distribution based on abundances observed in the data, then, still considering the same cells and no others, analyze the variability in the underlying biology.

##### Denoiser-based approach

A conceptually separate approach to estimating noise fractions does not involve model fits, but raw data and pre-processed/transformed (e.g., by applying a log1p transformation) data. This approach takes seriously the “denoiser” interpretation of transformations, which is both part of the motivation for applying these transformations in the first place, and appears empirically reasonable (see Figure 6c).

Let *ϕ* denote a transformation applied to observed RNA counts ***x*** := (*x*_*N*_, *x*_*M*_ )^*T*^ . If we assume that the *only* effect of transformations is to remove technical noise, and not any biologically meaningful variation, we can identify *ϕ*(***x***) with the latent count vector ***z*** := (*z*_*N*_, *z*_*M*_ )^*T*^ . In the previous subsection, we found that (under certain assumptions) the observed CV matrix is equal to the CV matrix of the analogous system without technical noise, plus a contribution due to technical noise. Under the denoiser assumption, we then have

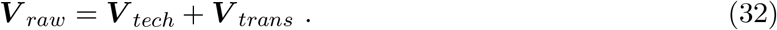

Since both ***V*** _*raw*_ and ***V*** _*trans*_ can be measured, ***V*** _*tech*_ can be inferred from their difference. We can disentangle intracellular and type-to-type variation by applying the transformation separately to data from each cell type. By the law of total covariance and the denoiser assumption (see Equation 30),

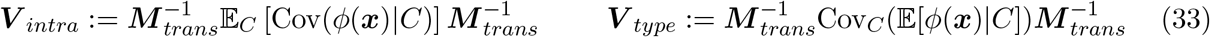

where ***M*** _*trans*_ denotes the diagonal matrix formed from the transformed mean 𝔼 [*ϕ*(***x***)].

##### Transformations used

We considered four commonly-used transformations: proportional fitting normalization (PF; *ϕ*_1_), log-transformation (log1p; *ϕ*_2_), principal component analysis (PCA; *ϕ*_3_), and Uniform Manifold Approximation and Projection (UMAP; *ϕ*_4_) [116]. Let *c* denote the number of cells and *g* the number of genes. Let *𝒳*_*N*_ ∈ N^*c×g*^ and *𝒳*_*M*_ ∈ N^*c×g*^ denote the raw nascent and mature count matrices, respectively. Let *𝒴*_*N*_ and *𝒴*_*M*_ denote possibly transformed matrices of the same dimensions.

PF involves computing a correction factor

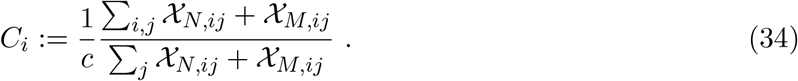

This factor involves averages over both cells and genes. The corresponding transformation is

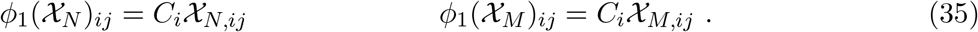

Log-transformation is simply defined via

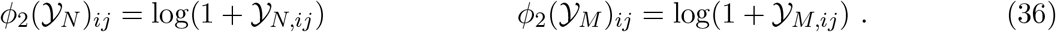

The reconstructed PCA projection *ϕ*_3_ is given by 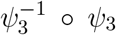, where *ψ*_3_ is implemented through the *scikit-learn* 1.0.1 function `transform` and 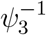 is implemented through the function `inverse_transform`, both associated with a `sklearn.decomposition.PCA` object [148]. The inverse transform is applied to ensure the count matrix has the same dimensions as the original, and hence to make the comparison more fair. We used 50 components to compute the PCA projection.

The reconstructed UMAP projection *ϕ*_4_ is given by 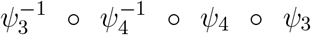, where *ψ*_4_ is implemented through the *umap-learn* 0.5.1 function `transform` and 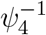 is implemented through the function `inverse_transform`, both associated with a `umap.UMAP` object [116]. We used the 50-dimensional PCA projection and default parameters to compute the UMAP projection.

In Figure 6c, PCA is applied to nascent and mature count matrices (without any additional transformations) separately. UMAP uses the same PCA decomposition in this panel. In the rest of Figure 6, PCA directions are determined by finding the principal vectors associated with the sum of the nascent and mature count matrices.

In Figure 6c, all transformations are applied individually. In the remaining panels, transformations are applied sequentially (PF→log1p→PCA→UMAP; *ϕ*_4_ *°ϕ*_3_ *°ϕ*_2_ *°ϕ*_1_), as they would be in a real pre-processing workflow; for example, in Figure 6d the label “PCA” depicts results for data transformed first via PF, then via log1p, and finally via PCA.

#### 9.5.4 Model-free bounds

The assumption we use to associate transformations with noise decompositions is that transformations *only* remove technical noise; this assumption might be overly strict, since for the purpose of many downstream analyses (e.g., identifying cell types or subtypes via clustering) preserving intracellular variation may not be viewed as crucial. Here, we relax that assumption and assume instead that transformations remove *all* technical noise (since the purpose of applying transformations is generally to produce “cleaner” data), and possibly *some* intracellular variation. This relaxed assumption implies (relatively) model-free upper and lower bounds for the CV matrix.

The upper bound corresponds to assuming *only* technical noise is removed, as we did previously. Assuming the catalysis technical noise model (c.f. Equation 30), define

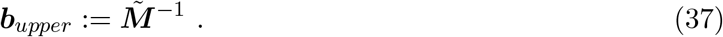

Then ***b***_*upper*_ − ***V*** _*trans*_ is positive semidefinite. The lower bound corresponds to assuming *both* technical and intracellular variation are completely removed, leaving only type-to-type variation. Motivated by Equation 22, define

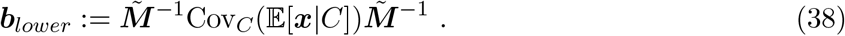

Then ***V*** _*trans*_ − ***b***_*lower*_ is positive semidefinite. Together, these results imply

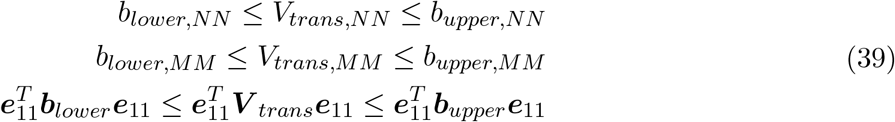

where we define 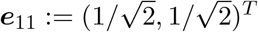. These are the “model-free” bounds used in Section 2.6. We use the second inequality (which bounds post-transformation mature RNA variation) in Figure 6d, and the last (which bounds post-transformation nascent-mature correlations) in Figure 6e.

#### 9.5.5 Single-species model-free bounds

Interestingly, the derivation in Section 9.5.4 can be made more generic yet. First, we relax the assumption of the technical noise model (Equation 37), essentially choosing to make no statements regarding potential retention of technical variability. In this scenario, we obtain a single lower bound per gene, the scalar version of Equation 38:

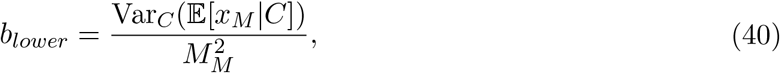

i.e., precisely the empirical, rather than theoretical, version of *f*_*M,type*_ implied by Equation 31, and conceptually a consequence of the law of total variance.

Next, we redefine the transformations in Section 9.5.4 to only involve *X*_*M*_, more closely matching the transformations actually used in practice, but choosing to make no statements regarding the retention of covariance between nascent and mature species. This amounts to defining an analog to Equation 34 as

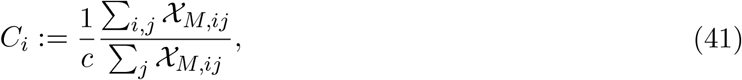

and otherwise applying the standard transformations sequentially. In this scenario, we obtain a theoretical baseline for mature RNA (Supplementary Figure 10a), analogous to the one derived in Section 9.5.4. Given this baseline, we can inspect the effect of transformations and dimensionality reductions upon the CV (Supplementary Figure 10c), particularly upon the high-mean, high-variance genes (Supplementary Figure 10b). As with the more generic case in Equation 32, the denoiser interpretation induces “attributions” of noise, with higher-expression genes losing a higher fraction of their CV (Supplementary Figure 10d-g). With these results and the baseline, we can readily plot the fractions of remaining noise, as in Figure 6d, but on a common scale rather than relative to the window between lower and upper bounds (Supplementary Figure 10h-k). In addition, we computed the fraction of genes violating the bound at each step: 0%, 7.5%, 43.6%, and 7.6% after proportional fitting, log-transformation, PCA, and UMAP respectively.

As the transformations are applied cumulatively, the distribution at step *i* may fall within the admissible region (above the theoretical baseline), but still be quantitatively degraded because it fell outside it at some step *i*^*′*^ *< i*. To characterize the loss of quantitative information about cell type relationships, we plotted the same data points as above, and colored them according to the history of the analysis. If a gene has ever violated the lower bound, we plotted it in a violet color. This analysis is shown in Supplementary Figure 12. In addition, we computed the fraction of genes identified after each step. We found that 0%, 7.5%, 43.6%, and 44.4% of the genes had at some point violated the bound after proportional fitting, log-transformation, PCA, and UMAP respectively.

Figure 6f suggests that the *Monod* model and the normalizations effectively produce opposite attributions: where the model suggests variability in high-expression genes (in red) is largely due to biological effects, the normalizations disproportionately remove variance from those very genes, implicitly encoding that the variability in high-expression genes is largely due to dispensable technical effects. We can use the model-based approach in Section 9.5.3 to estimate the biological variability attributions and agreement with the theoretical baseline (Supplementary Figure 11), with attributions that fundamentally do not cohere with the normalizations’ effects (Supplementary Figure 13). Interestingly, this attribution, which nominally adds the *f*_*M,intra*_ and *f*_*M,type*_ attributions from the mature RNA analog of Equation 31, mostly represents the former term: conflating all of the cell subtypes together and running *Monod* once on the entire glutamatergic dataset produces practically identical results (Supplementary Figure 14).

#### 9.5.6 Nascent/mature correlations and their connections to GRN inference

The procedure in Section 9.5.4 is separately applied to nascent and mature count matrices, whereas the one in Section 9.5.5 is applied to the mature count matrix only, to emphasize the consequences for the exonic (protein-coding) transcriptome the field is traditionally interested in. The same sequence of transformations may, in principle, be applied to the matrix *Ƶ* := *Ƶ*^*N*^ ⊕ *Ƶ*^*M*^, the concatenation of the nascent and mature counts, which yields a *c ×* 2*g* matrix [31]. Although the joint distribution of nascent and mature RNA can provide somewhat different biological [25,31,149] and technical [150, 151] insights than either of the individual modalities or their sum, this approach is not typical. However, it is instructive in the unexpected context of gene regulatory networks (GRNs).

GRN inference encompasses the identification of regulatory interactions between genes. The robust inference of such interactions is a key challenge in transcriptomics [6, 152–154], with a broad variety of strategies and prospects [152, 153, 155]. Although this subfield is far too broad to summarize in detail here, one key barrier is, conceptually, establishing causation from correlation: the strength of association between two genes may represent real regulatory motifs, or merely reflect confounding. A related, and more fundamental, challenge is the biochemistry of regulation: sequencing reads out noise-corrupted proxies for RNA content, whereas the regulation proceeds through protein and DNA interactions. In other words, even ostensibly known gene–gene interactions, *if* they are active in a particular system, may not be identifiable from RNA data alone.

The relationship between nascent and mature RNA provides a counterpoint. Mature RNA are produced from nascent RNA; causality is guaranteed. Although the process may have unobservable intermediate species, e.g., transient RNA molecules with no poly(A) content, the proximity of the source and product is far closer than in protein-mediated processes. Finally, this process is guaranteed to occur, whereas some GRN links may be inhibited in a particular biological system.

As expected, the nascent–mature correlation distribution skews higher than the overall distribution of mature–mature correlations (Supplementary Figure 15a). For a sizeable fraction of genes, the correlation between nascent and mature molecules for gene *j* is higher than inter-gene correlations between gene *j* and other genes *j*^*′*^≠ *j*. We quantify this by calculating the quantile of the nascent–mature correlation with respect to the empirical distribution of mature–mature correlations for each gene (Supplementary Figure 15b).

GRN inference typically uses normalized data as an input [152, 154]. We may naturally hope that this pre-processing treats two goals consistent with the impression of data transformation as “denoising”: (1) preserving the highest-magnitude, most evident relationships between nascent and mature RNA, and (2) enhancing the less prominent relationships. To that end, we quantify the transformations’ effect on absolute and relative correlations between nascent and mature RNA. Although this moment-based measure is relatively simplistic, it is in line with the broad range of popular methods that directly manipulate correlation matrices to identify gene interactions and modules [152, 153, 156].

We find that the nascent/mature correlations (Supplementary Figure 16) are substantially distorted by transformations, with the sign of the correlation being frequently inverted by size normalization, PCA, and UMAP. In addition, their magnitude relative to the mature/mature correlation distribution is generally not preserved, whether considering (Supplementary Figure 17) or omitting (Supplementary Figure 18) the signs.

Additionally, given *Monod* fits, we can directly compute biological correlations within a particular cell population. These correlations immediately follow from results in Section S1.4. We observe that the correlations follow the expected trend: the “underlying” correlations, prior to corruption by technical noise, are higher than observed correlations (Supplementary Figure 19).

#### 9.5.7 Dataset processing

##### Preprocessing: 10x PBMCs and Allen GABAergic Neurons

We obtained raw 10x v2 and v3 FASTQ files from the 10x Genomics website for the 1k PBMC samples [157, 158]. For the GABAergic cells we use the cell types (*Vip, Sst, Lamp5, Pvalb* neurons) from the B08 sample generated by the Allen Institute for Brain Science [56, 140], for which we previously performed the fits [24]. We quantified the datasets using *kallisto* | *bustools* 0.26.0 with a pre-built GRCh38 or mm10 genome from https://support.10xgenomics.com/single-cell-gene-expression/software/downloads/latest (2020-A version), as described in Section 9.1.

##### Filtering

For the PBMC datasets, we used the precomputed 10x Genomics *K*-means clustering results with *K* = 4, assigning cluster 1 to T cells, cluster 2 to monocytes, cluster 3 to B cells, and cluster 4 to erythrocytes and low-quality cells, determining these assignments by manual curation. We also selected cells with greater than 1,000 or 10,000 UMIs total (in the v2 and v3 data respectively). We then selected for the same genes across both 10x v2 and v3 PBMC datasets with moderate to high expression using *Monod*, as described in Section 9.1.2. From genes that passed these thresholds, we randomly selected 3,500 genes to fit.

##### Inference and analysis of biophysical parameters

For the PBMC datasets, based on previous technical sampling parameters fits in Gorin and Pachter [24], we set {log_10_ *C*_*N*_ = −6.105263157894737, log_10_ *λ*_*M*_ = −1.125} for inference with *Monod*. We fit the bursty model of transcription with (Poisson) length-biased sampling, as was done with the Allen GABAergic data, to each cell type. We then applied chi-squared testing and Hellinger distance criteria as in Section 9.1.3 to reject genes with poor fits.

## 10 Data availability

The datasets released by the Allen Institute for Brain Science were downloaded from http://data.nemoarchive.org/biccn/grant/u19_zeng/zeng/transcriptome/scell/10x_v3/mouse/raw/MOp/, and filtered according to the metadata annotations at http://data.nemoarchive.org/biccn/grant/u19_zeng/zeng/transcriptome/scell/10x_v3/mouse/processed/analysis/10X_cells_v3_AIBS/ [56, 140, 159]. The paired single-cell and single-nucleus mouse brain datasets, as well as the human PBMC datasets, were obtained from the 10x Genomics website. The control and IdU-perturbed mouse embryonic stem cell (mESC) data from the study by [47] was obtained from GSE176044 at the Gene Expression Omnibus (GEO). The mouse germ cell dataset from [57] was obtained from GEO repository GSE136220, the patient-derived PDAC tumor samples [61] from GSE202051, and the intestinal radation therapy data [60] from GSE165318. The single-nucleus mESC data [112] was obtained using the Sequence Read Archive run accession number SRR18364193. Fluorescent intensity values for seqFISH+ experiments on mESCs [111] were obtained from Zenodo package 7693825. The pre-built GRCh38 or mm10 genome from https://support.10xgenomics.com/single-cell-gene-expression/software/downloads/latest (2020-A version) was used for quantification of datasets.

All *loom* files with nascent and mature count matrices for the datasets used in this study and all *Monod* fits have been deposited as the Zenodo package 15051840 [160].

## 11 Code availability

Notebooks that reproduce all of the filtering, fitting, and analysis procedures have been deposited also in the Zenodo package 15051840 [160] and are available at https://github.com/pachterlab/monod_examples/tree/main/manuscript_computation. This GitHub repository also contains a Google Colaboratory notebook that illustrates the *Monod* workflow on a small dataset.

The *Monod* software used for all analysis is available as a pip installable package with an API available at https://monod-examples.readthedocs.io.

## S1 Supported models

Here we outline the models implemented in *Monod*, including the biological and technical noise models which encapsulate stochastic mRNA production and molecule capture by the sequencing process. We additionally provide the moment derivations for each for these models.

### S1.1 Biological noise models

In this section, we define the biological models of interest through their generating function definitions and solutions, which follow the general form outlined in Section 9.1.1.

#### S1.1.1 Bursty

The three-parameter bursty model describes the following biology:

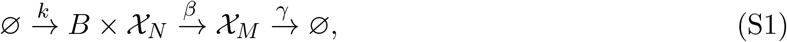

where *B* is a geometrically-distributed random variable on ℕ_0_ with mean *b*, and *k* is the burst frequency, set to unity with no loss of generality at steady state. This system has the following log-PGF [37]:

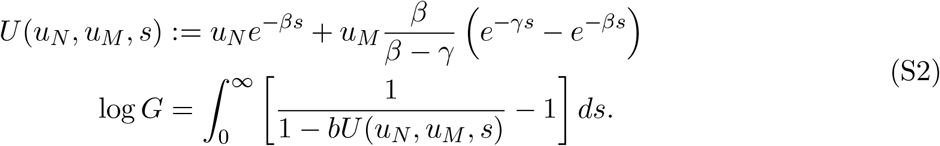

This model encodes the bursty production of mRNA, and is usually derived as the limit of a transcriptional process driven by a randomly switching promoter [27]. Specifically, if the promoter is described by a telegraph process with on rate *k*, off rate *k*_*off*_, and transcription rate in the on state *k*_*init*_, then in the physiological limit of rare, high-amplitude bursts, the Chemical Master Equation (CME) reduces to the memoryless process in Equation S1 with *b* := *k*_*init*_*/k*_*off*_ (Section S1.3 of [119]). In addition, bursty transcription can be obtained from an Ornstein-Uhlenbeck stochastic differential equation with exponentially-distributed jumps [23].

#### S1.1.2 Constitutive

The two-parameter constitutive model describes the following biology:

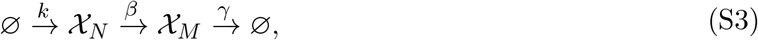

where *k* is set to unity with no loss of generality at steady state. This system has the following PGF [34]:

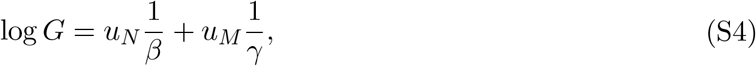

i.e., the joint distribution is the product of two independent Poisson distributions. This model encodes the unregulated production of mRNA.

#### S1.1.3 Extrinsic

The three-parameter extrinsic noise model describes the following biology:

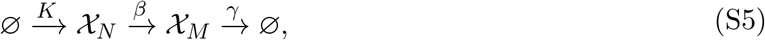

where *K* is a gamma distribution with shape *ν* and scale 1. This system has the following log-PGF:

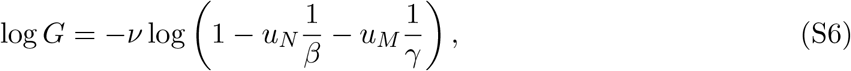

which follows immediately from the properties of generating functions [119]. This is the log-PGF of the bivariate negative binomial distribution, with negative binomial marginals [161]. This model encodes driving by a slow, Gamma-distributed transcriptional process [23, 162–164].

#### S1.1.4 CIR-like

The three-parameter CIR-like noise model describes the following biology:

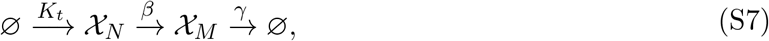

where *K*_*t*_ is, informally, the time derivative of a subordinating inverse Gaussian process [23]. This system emerges in the fast limit of transcription driven by a Cox–Ingersoll–Ross process, and has the following log-PGF:

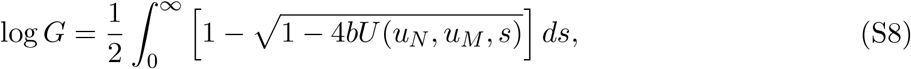

where *U* is identical to the expression in Equation S2, whereas *b* is a burst size-like “gain” that appears in the definition of *K*_*t*_. This model encodes driving by a fast chemical Langevin equation, which in turn represents a high-concentration limit of the constitutive birth-death process [23].

#### S1.1.5 Bursty with delayed splicing

The three-parameter bursty model with delayed splicing describes the following biology:

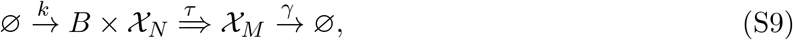

where *τ* is the deterministic retention time for *X*_*N*_ and *k* is set to unity with no loss of generality at steady state. This model has the following log-PGF:

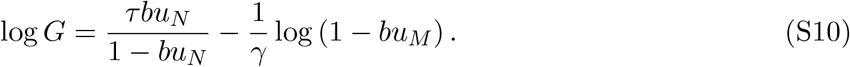

Since this equation is separable in *u*_*N*_ and *u*_*M*_, the PGF is the product of two independent marginal PGFs; the nascent marginal is geometric-Poisson [45, 165] whereas the mature marginal is negative binomial.

#### S1.1.6 Bursty with delayed degradation

The three-parameter bursty model with delayed degradation describes the following biology:

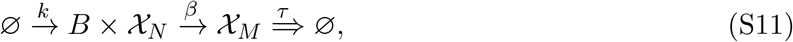

where *τ* is the deterministic retention time for *X*_*M*_ and *k* is set to unity with no loss of generality at steady state. This model has the following log-PGF [45]:

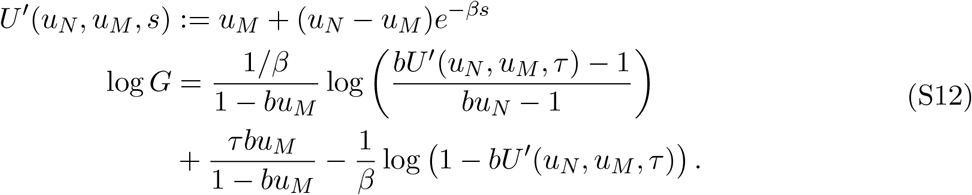

### S1.2 Technical noise models

In this section, we define the technical noise models of interest through their generating function definitions and solutions, which follow the general form outlined in Section 9.1.1. These models account for the artifacts and biases introduced during the sequencing process.

#### S1.2.1 Null process

In the simplest case, the null process, the sequencing is perfect and every mRNA is captured and reported as a unique molecule:

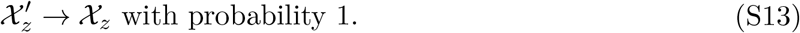

The PGF of this process is *G*_*z,t*_(*g*_*z*_) = *g*_*z*_ = *u*_*z*_ + 1, yielding *G*_*z,t*_(*u*_*z*_) − 1 = (*u*_*z*_ + 1) − 1 = *u*_*N*_, i.e., *G* = *H*.

#### S1.2.2 Bernoulli process

In the Bernoulli process, each molecule has probability *p*_*z*_ of being captured and reported; multiple priming is impossible:

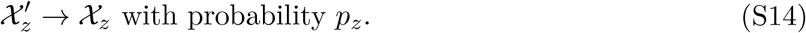

The PGF of this process is *G*_*z,t*_(*g*_*z*_) = *p*_*z*_*g*_*z*_ + (1 − *p*_*z*_) = *p*_*z*_*u*_*z*_ + 1, yielding *G*_*z,t*_(*u*_*z*_) − 1 = *p*_*z*_*u*_*z*_.

#### S1.2.3 Poisson process

In the Poisson process, each molecule has a rate *λ*_*z*_ of being captured and reported; multiple priming is possible, but rare if *λ*_*z*_ is low:

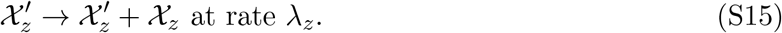

This PGF of this process over unity time is 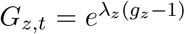, i.e., 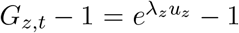, which cannot be simplified further.

### S1.3 Moment computations

It is numerically convenient to initialize optimization at the method of moments estimate [24]. We can avoid manual computations (Section S1.2 of [24]) by symbolically manipulating the generating functions with the MATLAB R2022a Symbolic Math Toolbox [166, 167].

#### S1.3.1 First moments

First, we recall that the probability-generating function can be differentiated once to obtain the marginal averages. For example,

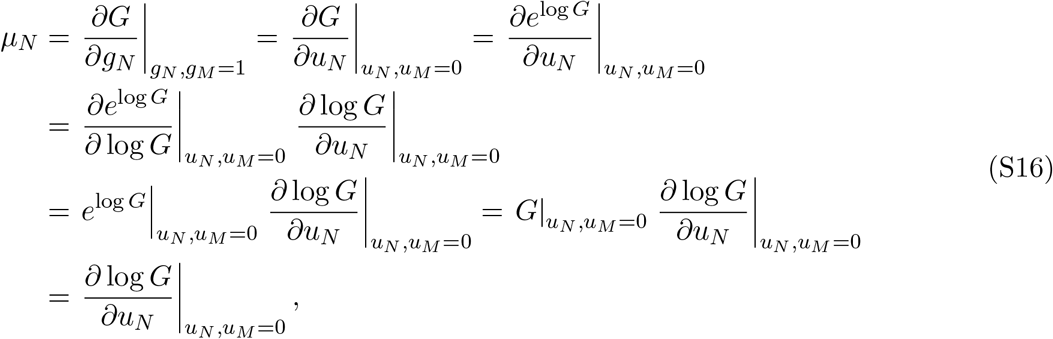

where the third line follows from applying the definition of the exponential function, and the fourth uses the normalization criterion, whereby *G* = 1 at *u*_*N*_ = *u*_*M*_ = 0. Analogously,

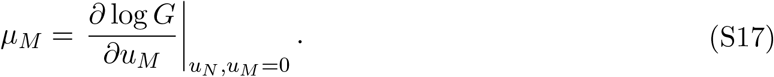

For the closed-form log *G* expressions, we can evaluate this quantity directly. For the expressions that involve an integral, we need to interchange operators:

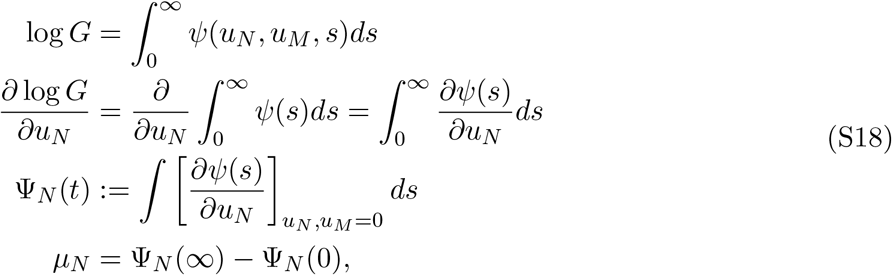

and analogously,

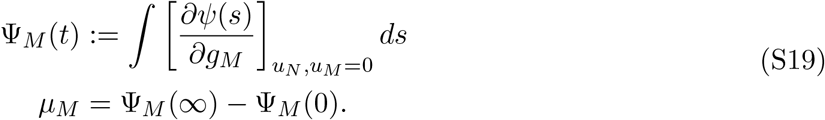

#### S1.3.2 Second moments

The second moments can be obtained similarly:

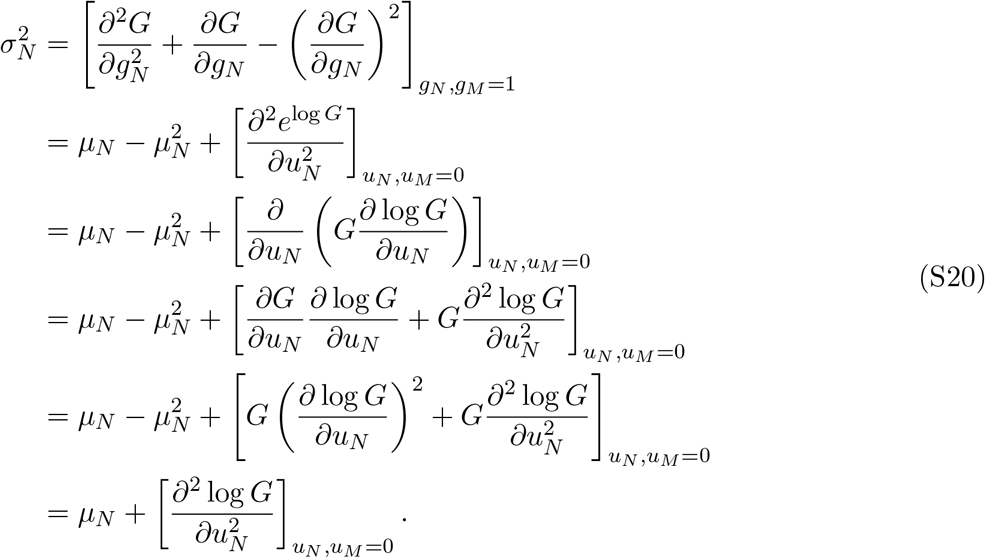

For the closed-form log *G* expressions, we can evaluate this quantity directly. Otherwise, we need to use symbolic integration:

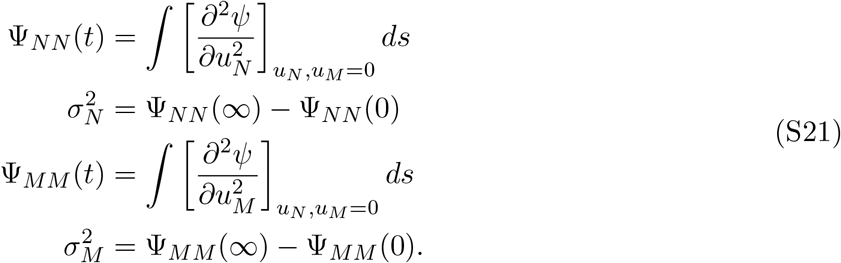

Finally, to obtain the covariance Cov_*NM*_, we exploit the following identity:

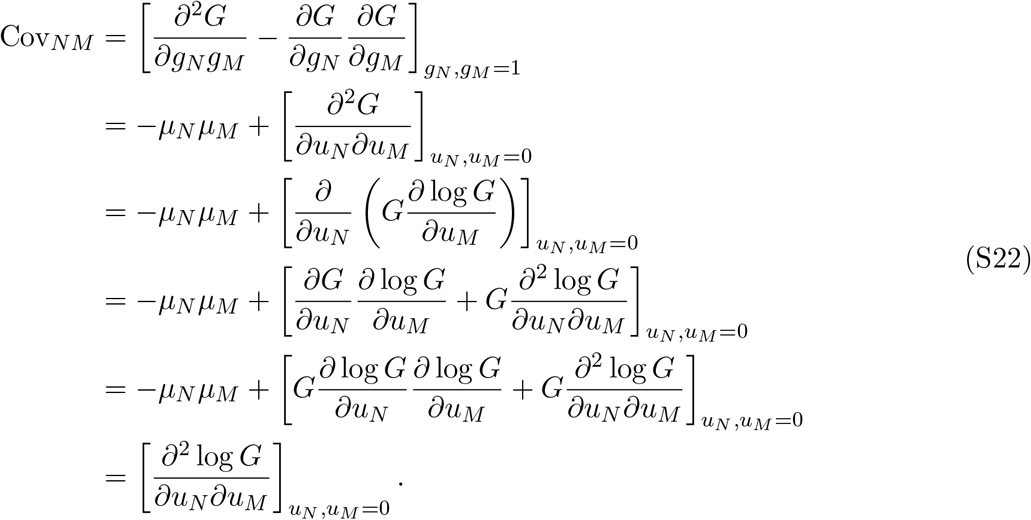

Again, if log *G* is analytically tractable, we can evaluate this quantity directly. If it is not, we can integrate:

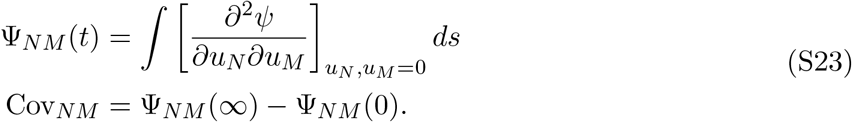

### S1.4 Moment results

#### S1.4.1 Null process

We report the results for the noise-free model in Supplementary Table 1. For compactness, we report the shifted Fano factor, defined as 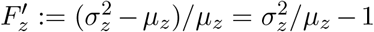, instead of variances. We use 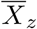 to indicate the sample average for species 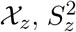 to indicate the sample variance, and 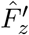 to indicate the sample shifted Fano factor 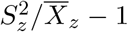. We report the method of moments estimates in Supplementary Table 2. These estimates are not unique; we default to using the nascent RNA data for simplicity.

#### S1.4.2 Bernoulli process

We report the results for the Bernoulli dropout model, conditional on a particular set of *p*_*N*_ and *p*_*M*_, in Supplementary Table 3. To compute these moments, we used the procedure in Section S1.3, defining log *G* or *ψ* in terms of auxiliary variables 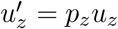, then computing the derivatives with respect to *u*_*z*_ as usual. We report the method of moment estimates in Supplementary Table 4. Evidently, bespoke routines are unnecessary: in practice, we rescale the results from Supplementary Table 2.

#### S1.4.3 Poisson process

We report the results for the Poisson sequencing model, conditional on a particular set of *λ*_*N*_ and *λ*_*M*_, in Supplementary Table 5. To compute these moments, we used the procedure in Section S1.3, defining log *G* or *ψ* in terms of auxiliary variables 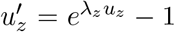, then computing the derivatives with respect to *u*_*z*_ as usual. We report the method of moment estimates in Supplementary Table 6. Here, we can likewise reuse the null process estimates.

### S1.5 Noise decomposition

The models afford analytical noise decompositions. For example, we can define the mature RNA squared coefficient of variation 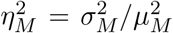 under the bursty model. If we assume there is no technical noise, we find:

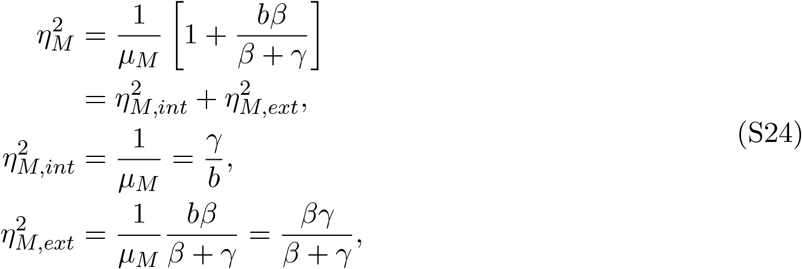

where 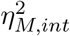 is the noise contribution from the discrete nature of the process and 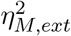 is the noise contribution due to the bursty transcriptional dynamics. We denote this noise component as *extrinsic*, as it emerges from variation in the transcription rate [23]. Interestingly, *b* does not explicitly occur in 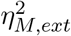 : the value of the burst size serves as a scaling factor and cancels out in the definition of CV^2^.

Under the Bernoulli technical noise model, we find that the following behavior holds:

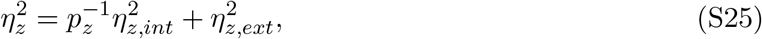

i.e., the variation induced by imperfect sampling is multiplicative with respect to the intrinsic noise term. We can find the fraction of variation attributable to each noise component:

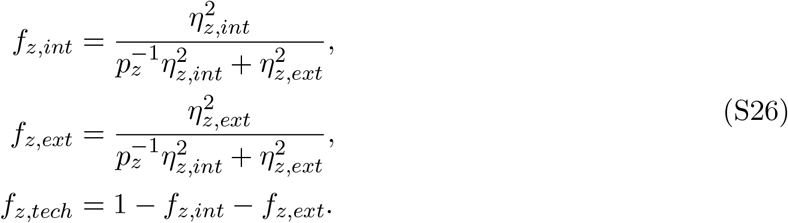

Under the Poisson technical noise model, we find that the following behavior holds:

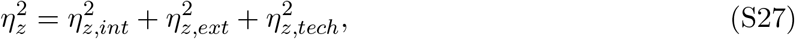

i.e., the variation induced by Poisson-like sampling is additive. This additive term is 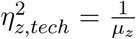 which depends on *λ*_*z*_. We can find the fraction of noise attributable to each noise component:

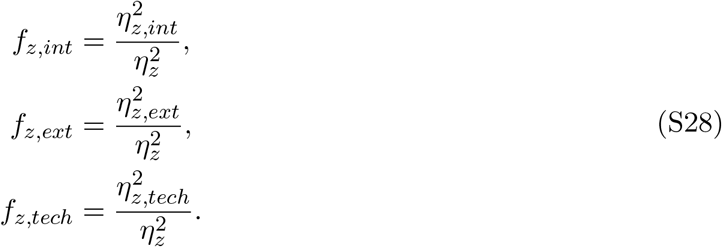

## S2 Supplementary tables

### S2.1 Theoretical results

#### S2.1.1 Null process

**Supplementary Table 1:** Lower moments of the null process.

| Model | $\mu_N$ | $\mu_M$ | $F'_N$ | $F'_M$ | $\text{Cov}_{NM}$ |
| --- | --- | --- | --- | --- | --- |
| Bursty | $\frac{b}{\beta}$ | $\frac{b}{\gamma}$ | $b$ | $\frac{b\beta}{\beta+\gamma}$ | $\frac{b^2}{\beta+\gamma}$ |
| Constitutive | $\frac{1}{\beta}$ | $\frac{1}{\gamma}$ | 0 | 0 | 0 |
| Extrinsic | $\frac{\nu}{\beta}$ | $\frac{\nu}{\gamma}$ | $\frac{1}{\beta}$ | $\frac{1}{\gamma}$ | $\frac{\nu}{\beta\gamma}$ |
| CIR-like | $\frac{b}{\beta}$ | $\frac{b}{\gamma}$ | $b$ | $\frac{b\beta}{\beta+\gamma}$ | $\frac{b^2}{\beta+\gamma}$ |
| Delayed splicing | $b\tau$ | $\frac{b}{\gamma}$ | $2b$ | $b$ | 0 |
| Delayed degradation | $\frac{b}{\beta}$ | $b\tau$ | $b$ | $\frac{2b}{\beta\tau}(\beta\tau + e^{-\beta\tau} - 1)$ | $\frac{b^2}{\beta}(1 - e^{-\beta\tau})$ |

**Supplementary Table 2:** Method of moments biological parameter estimates for the null process.

| Model | $\hat{b}$ | $\hat{\beta}$ | $\hat{\gamma}$ | $\hat{\nu}$ | $\hat{\tau}^{-1}$ |
| --- | --- | --- | --- | --- | --- |
| Bursty | $\hat{F}'_N$ | $\hat{b}/\bar{X}_N$ | $\hat{b}/\bar{X}_M$ | | |
| Constitutive | | $1/\bar{X}_N$ | $1/\bar{X}_M$ | | |
| Extrinsic | | $\hat{\nu}/\bar{X}_N$ | $\hat{\nu}/\bar{X}_M$ | $\frac{\bar{X}_N}{\hat{F}'_N}$ | |
| CIR-like | $\hat{F}'_N$ | $\hat{b}/\bar{X}_N$ | $\hat{b}/\bar{X}_M$ | | |
| Delayed splicing | $\hat{F}'_N/2$ | | $\hat{b}/\bar{X}_M$ | | $\hat{b}/\bar{X}_N$ |
| Delayed degradation | $\hat{F}'_N$ | $\hat{b}/\bar{X}_N$ | | | $\hat{b}/\bar{X}_M$ |

#### S2.1.2 Bernoulli process

**Supplementary Table 3:** Lower moments of the Bernoulli process.

| Model | $\mu_N$ | $\mu_M$ | $F'_N$ | $F'_M$ | $\text{Cov}_{NM}$ |
| --- | --- | --- | --- | --- | --- |
| Bursty | $\frac{bp_N}{\beta}$ | $\frac{bp_M}{\gamma}$ | $bp_N$ | $\frac{bp_M\beta}{\beta+\gamma}$ | $\frac{b^2p_Np_M}{\beta+\gamma}$ |
| Constitutive | $\frac{p_N}{\beta}$ | $\frac{p_M}{\gamma}$ | 0 | 0 | 0 |
| Extrinsic | $\frac{\nu p_N}{\beta}$ | $\frac{\nu p_M}{\gamma}$ | $\frac{p_N}{\beta}$ | $\frac{p_M}{\gamma}$ | $\frac{\nu p_N p_M}{\beta\gamma}$ |
| CIR-like | $\frac{bp_N}{\beta}$ | $\frac{bp_M}{\gamma}$ | $bp_N$ | $\frac{bp_M\beta}{\beta+\gamma}$ | $\frac{b^2p_Np_M}{\beta+\gamma}$ |
| Delayed splicing | $b\tau p_N$ | $\frac{bp_M}{\gamma}$ | $2bp_N$ | $bp_M$ | 0 |
| Delayed degradation | $\frac{bp_N}{\beta}$ | $b\tau p_M$ | $bp_N$ | $\frac{2bp_M}{\beta\tau}(\beta\tau + e^{-\beta\tau} - 1)$ | $\frac{b^2p_Np_M}{\beta}(1 - e^{-\beta\tau})$ |

**Supplementary Table 4:** Method of moments biological parameter estimates for the Bernoulli process.

| Model | $\hat{b}$ | $\hat{\beta}$ | $\hat{\gamma}$ | $\hat{\nu}$ | $\hat{\tau}^{-1}$ |
| --- | --- | --- | --- | --- | --- |
| Bursty | $\hat{F}'_N/p_N$ | $\hat{b}p_N/\bar{X}_N$ | $\hat{b}p_M/\bar{X}_M$ | | |
| Constitutive | | $p_N/\bar{X}_N$ | $p_M/\bar{X}_M$ | | |
| Extrinsic | | $\hat{\nu}p_N/\bar{X}_N$ | $\hat{\nu}p_M/\bar{X}_M$ | $\frac{\bar{X}_N}{\hat{F}'_N}$ | |
| CIR-like | $\hat{F}'_N/p_N$ | $\hat{b}p_N/\bar{X}_N$ | $\hat{b}p_M/\bar{X}_M$ | | |
| Delayed splicing | $\hat{F}'_N/(2p_N)$ | | $\hat{b}p_M/\bar{X}_M$ | | $\hat{b}p_N/\bar{X}_N$ |
| Delayed degradation | $\hat{F}'_N/p_N$ | $\hat{b}p_N/\bar{X}_N$ | | | $\hat{b}p_M/\bar{X}_M$ |

#### S2.1.3 Poisson process

**Supplementary Table 5:**
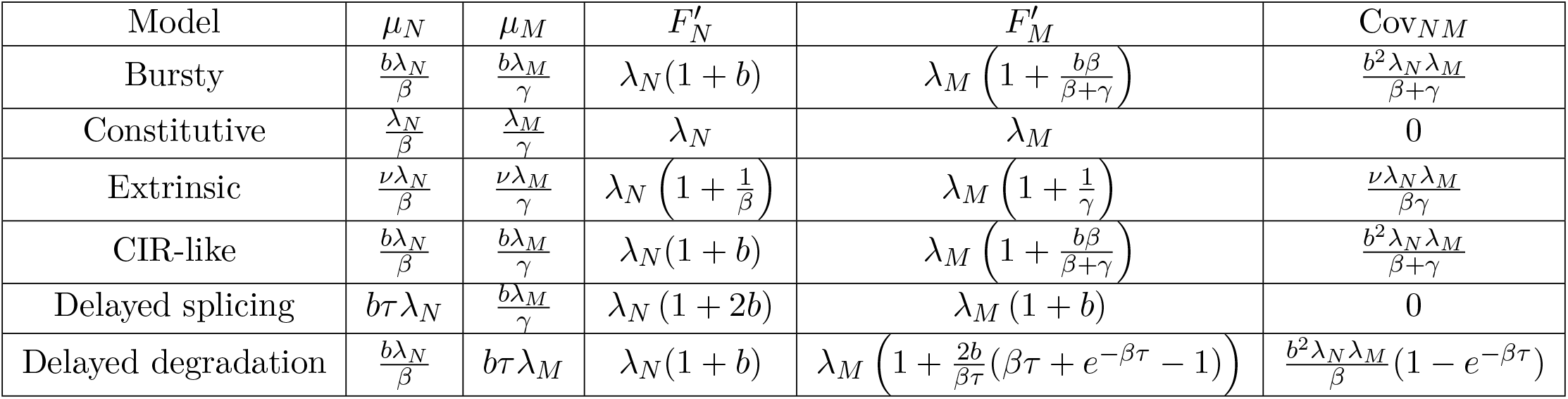
Lower moments of the Poisson process.

| Model | $\mu_N$ | $\mu_M$ | $F'_N$ | $F'_M$ | $\text{Cov}_{NM}$ |
| --- | --- | --- | --- | --- | --- |
| Bursty | $\frac{b\lambda_N}{\beta}$ | $\frac{b\lambda_M}{\gamma}$ | $\lambda_N(1+b)$ | $\lambda_M \left(1 + \frac{b\beta}{\beta+\gamma}\right)$ | $\frac{b^2\lambda_N\lambda_M}{\beta+\gamma}$ |
| Constitutive | $\frac{\lambda_N}{\beta}$ | $\frac{\lambda_M}{\gamma}$ | $\lambda_N$ | $\lambda_M$ | 0 |
| Extrinsic | $\frac{\nu\lambda_N}{\beta}$ | $\frac{\nu\lambda_M}{\gamma}$ | $\lambda_N \left(1 + \frac{1}{\beta}\right)$ | $\lambda_M \left(1 + \frac{1}{\gamma}\right)$ | $\frac{\nu\lambda_N\lambda_M}{\beta\gamma}$ |
| CIR-like | $\frac{b\lambda_N}{\beta}$ | $\frac{b\lambda_M}{\gamma}$ | $\lambda_N(1+b)$ | $\lambda_M \left(1 + \frac{b\beta}{\beta+\gamma}\right)$ | $\frac{b^2\lambda_N\lambda_M}{\beta+\gamma}$ |
| Delayed splicing | $b\tau\lambda_N$ | $\frac{b\lambda_M}{\gamma}$ | $\lambda_N(1+2b)$ | $\lambda_M(1+b)$ | 0 |
| Delayed degradation | $\frac{b\lambda_N}{\beta}$ | $b\tau\lambda_M$ | $\lambda_N(1+b)$ | $\lambda_M \left(1 + \frac{2b}{\beta\tau}(\beta\tau + e^{-\beta\tau} - 1)\right)$ | $\frac{b^2\lambda_N\lambda_M}{\beta}(1 - e^{-\beta\tau})$ |

**Supplementary Table 6:** Method of moments biological parameter estimates for the Poisson process.

| Model | $\hat{b}$ | $\hat{\beta}$ | $\hat{\gamma}$ | $\hat{\nu}$ | $\hat{\tau}^{-1}$ |
| --- | --- | --- | --- | --- | --- |
| Bursty | $\hat{F}'_N/\lambda_N - 1$ | $\hat{b}\lambda_N/\bar{X}_N$ | $\hat{b}\lambda_M/\bar{X}_M$ | | |
| Constitutive | | $\lambda_N/\bar{X}_N$ | $\lambda_M/\bar{X}_M$ | | |
| Extrinsic | | $\hat{\nu}\lambda_N/\bar{X}_N$ | $\hat{\nu}\lambda_M/\bar{X}_M$ | $\frac{\bar{X}_N}{\hat{F}'_N - \lambda_N}$ | |
| CIR-like | $\hat{F}'_N/\lambda_N - 1$ | $\hat{b}\lambda_N/\bar{X}_N$ | $\hat{b}\lambda_M/\bar{X}_M$ | | |
| Delayed splicing | $\hat{F}'_N/(2\lambda_N) - 1/2$ | | $\hat{b}\lambda_M/\bar{X}_M$ | | $\hat{b}\lambda_N/\bar{X}_N$ |
| Delayed degradation | $\hat{F}'_N/\lambda_N - 1$ | $\hat{b}\lambda_N/\bar{X}_N$ | | | $\hat{b}\lambda_M/\bar{X}_M$ |

### S2.2 Inference parameters

**Supplementary Table 7:** Search parameter bounds for grid scans and gradient descent.

| Parameter | Lower bound ($\log_{10}$) | Upper bound ($\log_{10}$) |
| --- | --- | --- |
| $C_N$ | -7.5 | -5.5 |
| $\lambda_M$ | -2 | 0 |
| $b$ | -1 | 4.2 |
| $\beta$ | -1.8 | 2.5 |
| $\gamma$ | -1.8 | 3.5 |

### S2.3 Datasets

**Supplementary Table 8:**
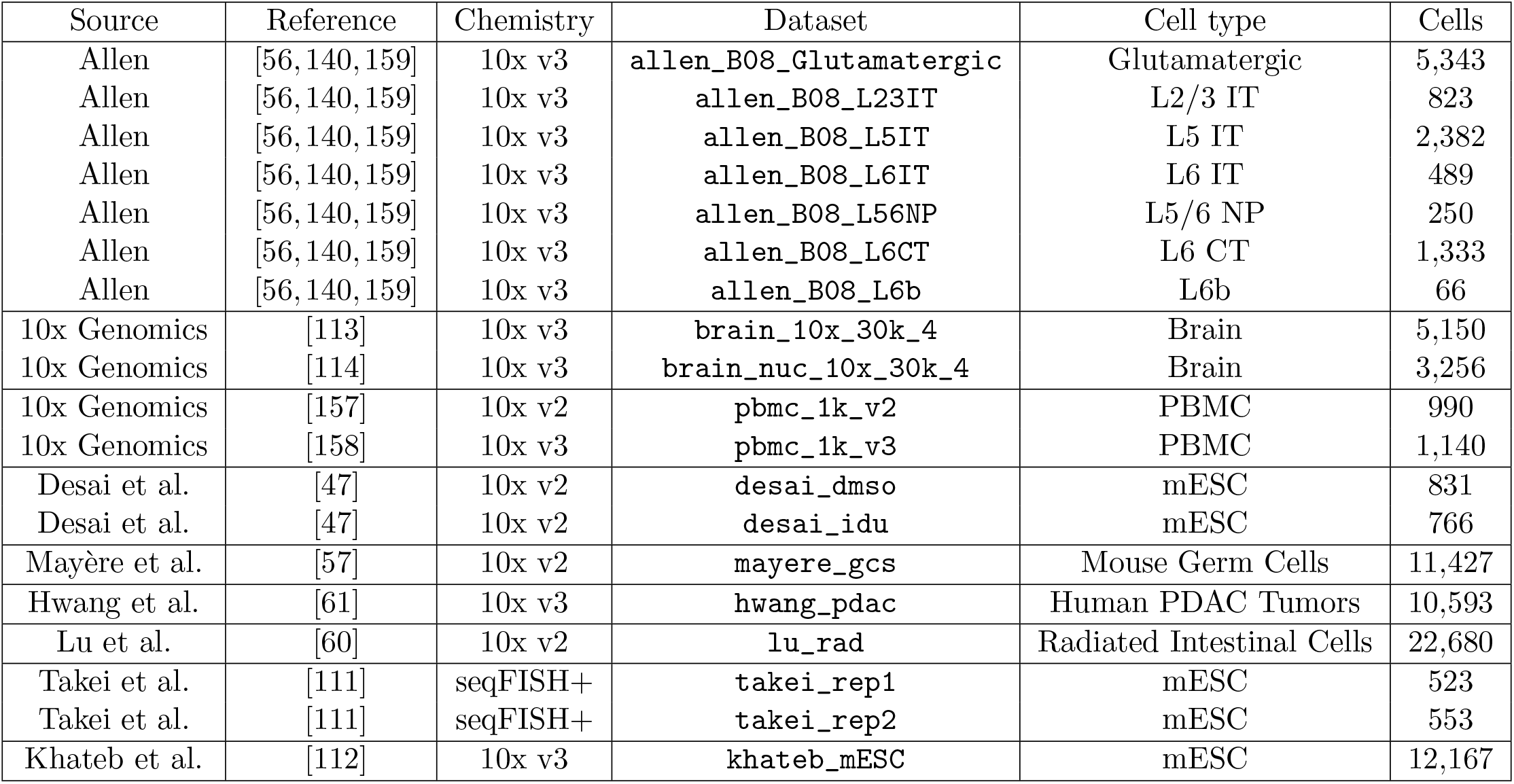
Summary of datasets used in the analysis. Unless otherwise noted, data originate from *M. musculus*. Data from PDAC tumors [61] are human samples. Cell counts are determined after bustools filtering for cells and after subsetting for the cell types used in analysis (based on prior annotation).

### S2.4 Inference timing

**Supplementary Table 9:** Summary of inference runtimes per biological model for mouse neurons from Allen sample B08 [56, 140, 159]. All models were run as described in Methods Section 6.2.3 over a 10 by 11 grid (110 gridpoints) of technical sampling parameters with a maximum of 15 iterations and 1 restart. Inference was performed on an Intel i9-10920X X-series Processor, 3.5GHz with a maximum of 20 processors (on 12 cores).

| Cell type | Cells | Genes | Model | Runtime | Runtime/gridpoint |
| --- | --- | --- | --- | --- | --- |
| L2/3 IT | 823 | 2,951 | Bursty | 181 min 17 seconds | 99 seconds |
| L2/3 IT | 823 | 2,951 | Constitutive | 31 min 14 seconds | 17 seconds |
| L2/3 IT | 823 | 2,951 | Extrinsic | 45 min 14 seconds | 25 seconds |
| L2/3 IT | 823 | 2,951 | Delayed degradation | 42 min 19 seconds | 23 seconds |
| L2/3 IT | 823 | 2,951 | Delayed splicing | 41 min 54 seconds | 22 seconds |
| L5 IT | 2,382 | 2,951 | Bursty | 243 min 43 seconds | 132 seconds |
| L5 IT | 2,382 | 2,951 | Constitutive | 34 min 30 seconds | 19 seconds |
| L5 IT | 2,382 | 2,951 | Extrinsic | 51 min 47 seconds | 28 seconds |
| L5 IT | 2,382 | 2,951 | Delayed degradation | 49 min 24 seconds | 27 seconds |
| L5 IT | 2,382 | 2,951 | Delayed splicing | 46 min 2 seconds | 25 seconds |
| L6 IT | 489 | 2,951 | Bursty | 137 min 10 seconds | 75 seconds |
| L6 IT | 489 | 2,951 | Constitutive | 28 min 42 seconds | 15 seconds |
| L6 IT | 489 | 2,951 | Extrinsic | 39 min 20 seconds | 21 seconds |
| L6 IT | 489 | 2,951 | Delayed degradation | 36 min 7 seconds | 20 seconds |
| L6 IT | 489 | 2,951 | Delayed splicing | 36 min 34 seconds | 19 seconds |
| L5/6 NP | 250 | 2,951 | Bursty | 77 min 37 seconds | 42 seconds |
| L5/6 NP | 250 | 2,951 | Constitutive | 22 min 10 seconds | 12 seconds |
| L5/6 NP | 250 | 2,951 | Extrinsic | 30 min 56 seconds | 16 seconds |
| L5/6 NP | 250 | 2,951 | Delayed degradation | 28 min 32 seconds | 15 seconds |
| L5/6 NP | 250 | 2,951 | Delayed splicing | 29 min 39 seconds | 16 seconds |
| L6 CT | 1,333 | 2,951 | Bursty | 159 min 15 seconds | 87 seconds |
| L6 CT | 1,333 | 2,951 | Constitutive | 30 min 50 seconds | 16 seconds |
| L6 CT | 1,333 | 2,951 | Extrinsic | 42 min 10 seconds | 23 seconds |
| L6 CT | 1,333 | 2,951 | Delayed degradation | 39 min 22 seconds | 21 seconds |
| L6 CT | 1,333 | 2,951 | Delayed splicing | 39 min 34 seconds | 21 seconds |
| L6b | 66 | 2,951 | Bursty | 100 min 24 seconds | 55 seconds |
| L6b | 66 | 2,951 | Constitutive | 23 min 59 seconds | 13 seconds |
| L6b | 66 | 2,951 | Extrinsic | 32 min 19 seconds | 18 seconds |
| L6b | 66 | 2,951 | Delayed degradation | 31 min 31 seconds | 17 seconds |
| L6b | 66 | 2,951 | Delayed splicing | 30 min 41 seconds | 16 seconds |

### S2.5 Inference memory usage

**Supplementary Table 10:** Maximum memory usage during inference for mouse T cells (samples from day 1) from Lu et al.’s radiation treatment study [60]. The bursty model was run as described in Methods Section 6.2.3 (though, for memory profiling purpose, at only a single gridpoint of Poisson sampling parameters). Inference was performed on an Intel i9-10920X X-series Processor, 3.5GHz with a maximum of 20 processors (on 12 cores).

| Sample | Cells | Genes | Model | Max. memory usage (MiB) |
| --- | --- | --- | --- | --- |
| S0 | 536 | 10 | Bursty | 303.3 |
| S0 | 536 | 100 | Bursty | 305.3 |
| S0 | 536 | 500 | Bursty | 321.8 |
| S0 | 536 | 1000 | Bursty | 343.7 |
| S22 | 404 | 10 | Bursty | 380.3 |
| S22 | 404 | 100 | Bursty | 302.9 |
| S22 | 404 | 500 | Bursty | 314.7 |
| S22 | 404 | 1000 | Bursty | 333.5 |
| S23 | 211 | 10 | Bursty | 377.1 |
| S23 | 211 | 100 | Bursty | 338.1 |
| S23 | 211 | 500 | Bursty | 338.0 |
| S23 | 211 | 1000 | Bursty | 337.5 |
| S26 | 798 | 10 | Bursty | 311.8 |
| S26 | 798 | 100 | Bursty | 313.0 |
| S26 | 798 | 500 | Bursty | 340.4 |
| S26 | 798 | 1000 | Bursty | 337.1 |

## S3 Supplementary figures

### S3.1 *Monod* workflow

**Supplementary Figure 1:**
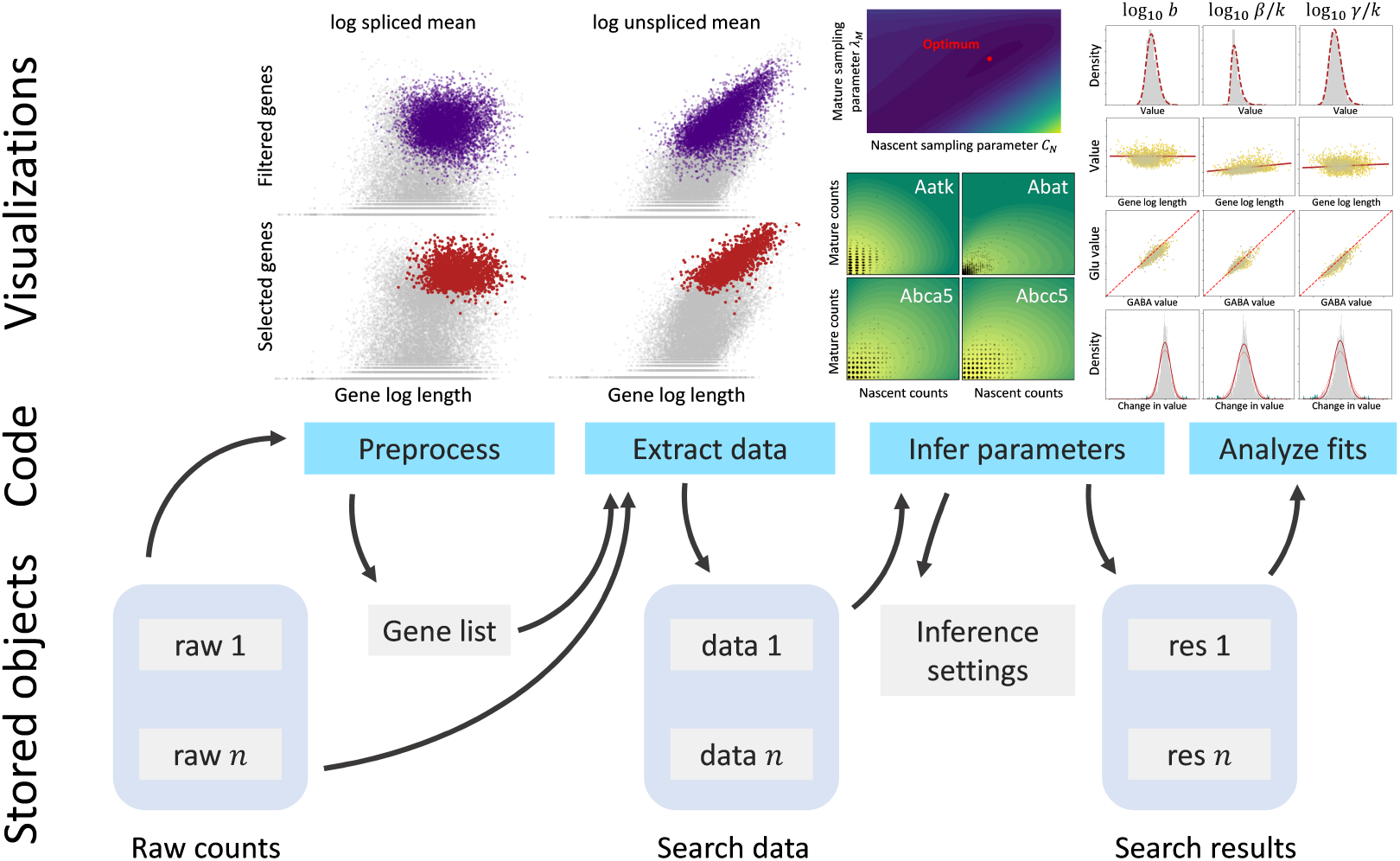
A schematic representation of the *Monod* workflow. Center: steps of the algorithm. Top: visualizations produced for the user. Bottom: data objects stored on disk.

### S3.2 Data and Parameter QC

**Supplementary Figure 2:**
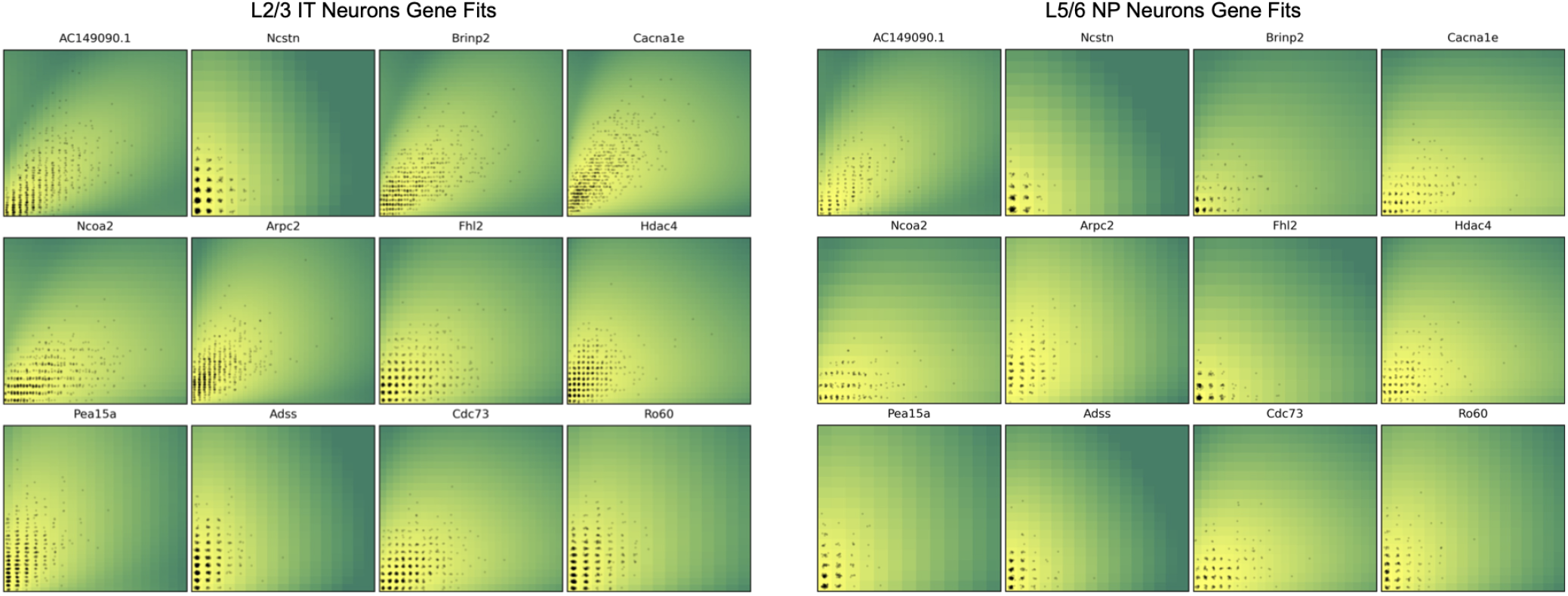
Empirical 2D histograms of genes in L2/3 IT or L5/6 NP neurons of the Allen B08 sample data in black. Gradient colors in the background denote 2D histograms of nascent and mature mRNA as calculated from the inferred parameters.

**Supplementary Figure 3:**
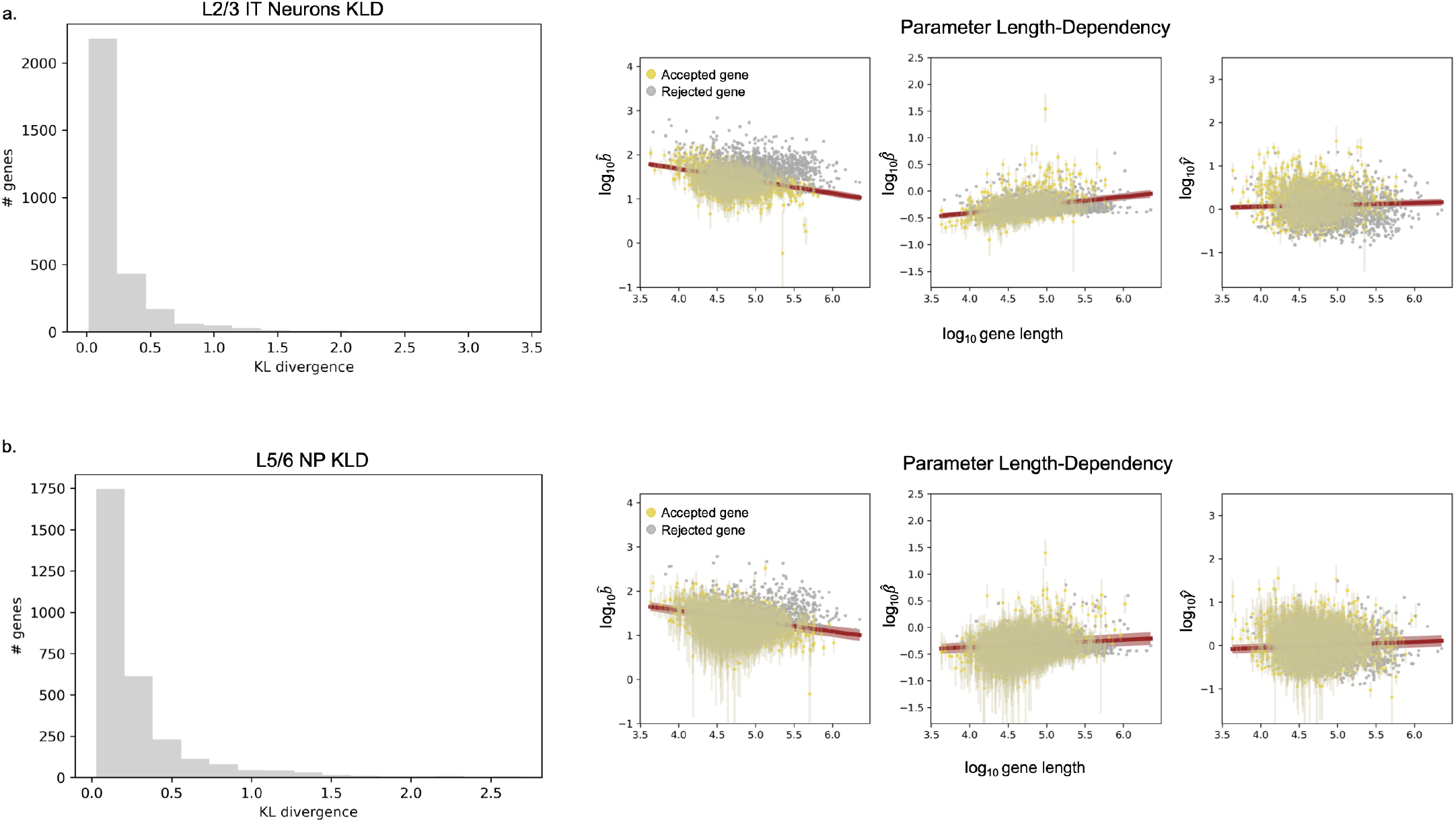
**a**. Histogram of KLDs for each gene fit for the L2/3 IT neurons under the bursty model of transcription with length-biased capture. Right plots show inferred parameters versus gene length, with golden dots denoting not rejected genes. Errors bars denote the 95% C.I. **b**. The same plots for the L5/6 NP neurons and inferred parameters. 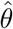 denotes inferred parameters.

### S3.3 Noise Modulation in Malignant CRTl-treated PDAC

**Supplementary Figure 4:**
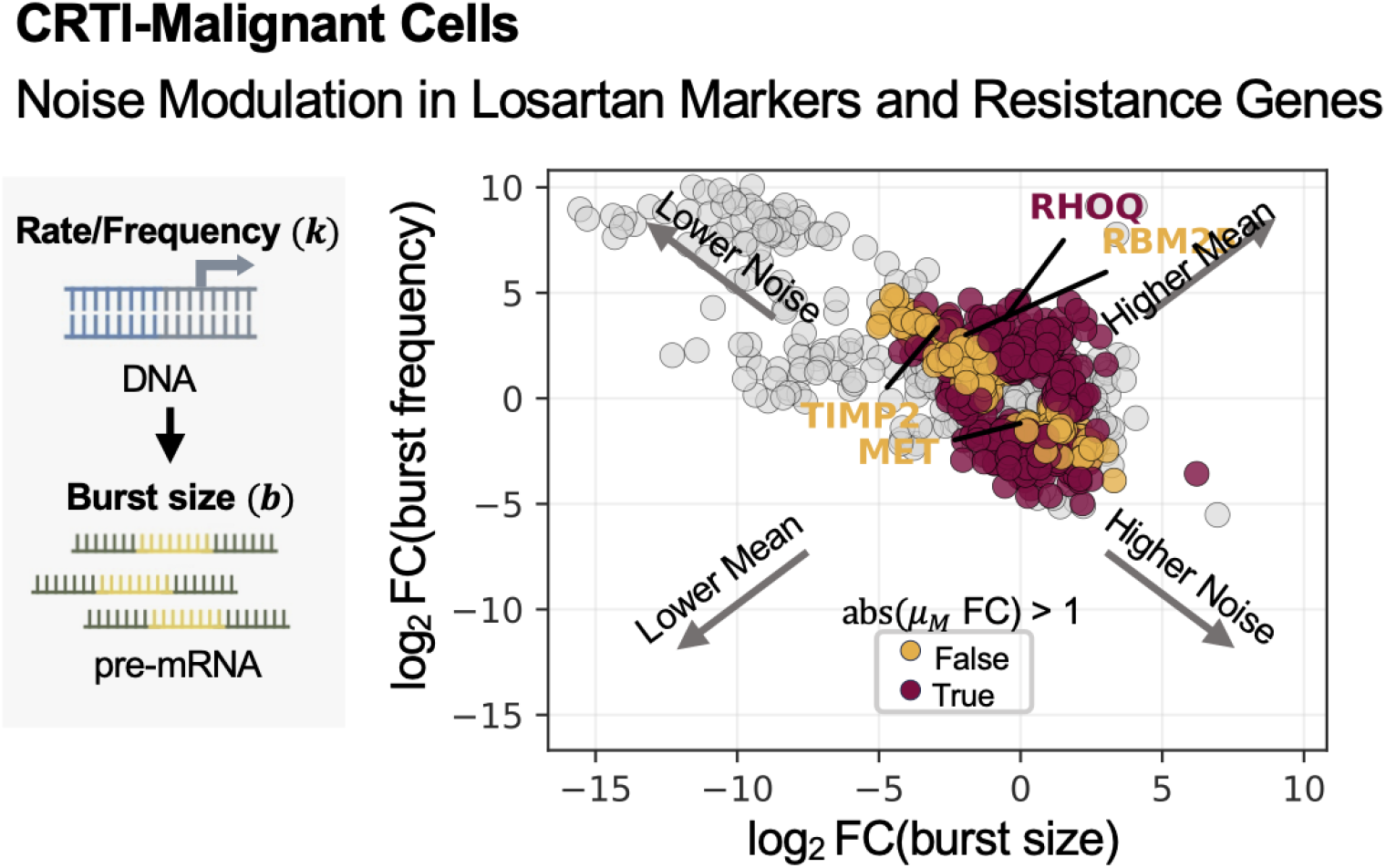
Plot of the log_2_ FC in burst frequency versus burst size for all non-rejected genes in n=182 CRTl-malignant cells. Genes with log_2_ FC *>* 1 in both/either parameter colored. Red denotes mean expression abs(log_2_ FC) *>* 1 and yellow denotes abs(log_2_ FC) *<* 1 (little/no change)

### S3.4 Sampling parameter landscapes

**Supplementary Figure 5:**
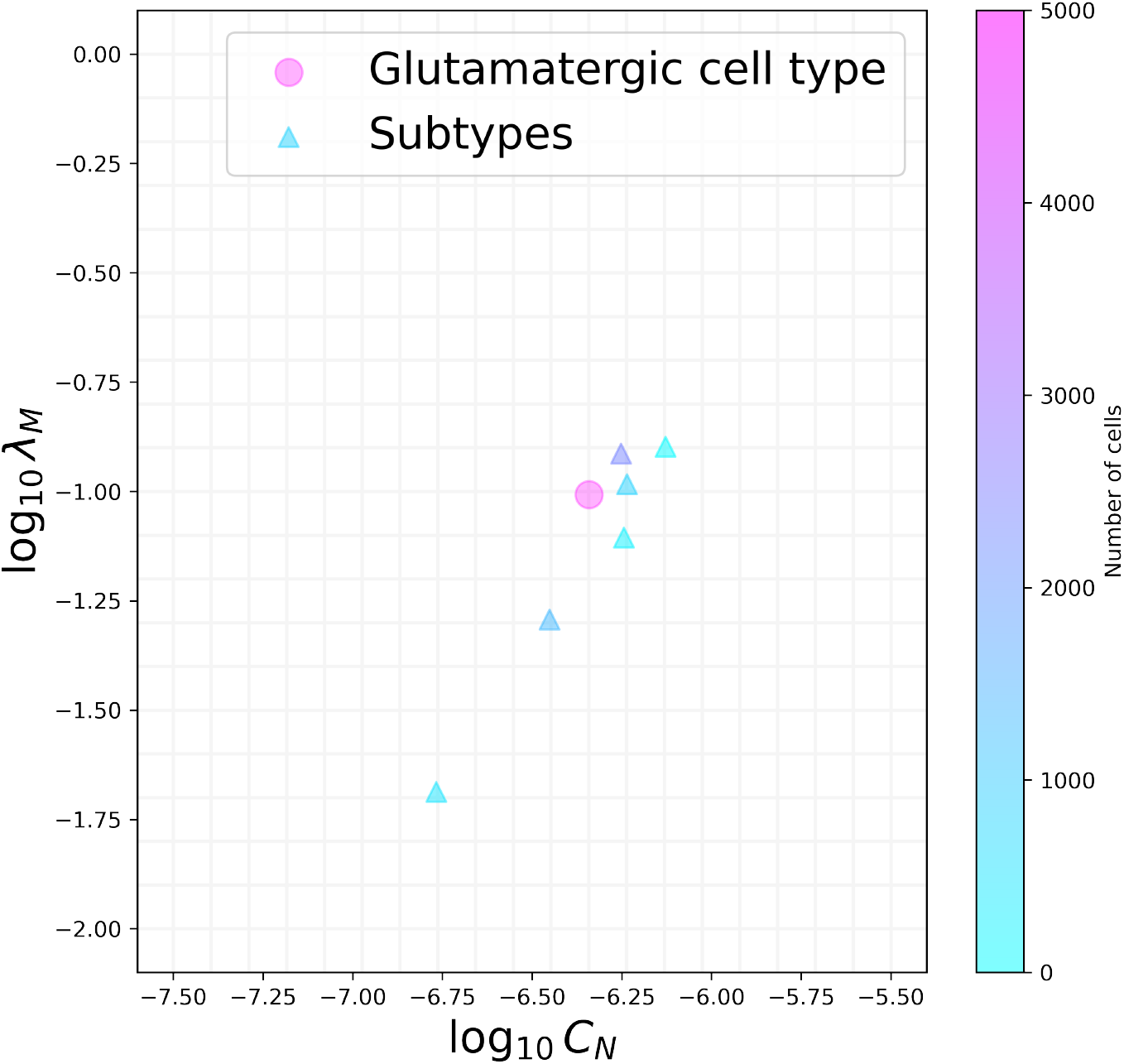
Inferred locations of technical noise parameters are largely concordant between cell subtypes, but show deviations at low cell numbers, motivating us to use the largest available dataset as the best estimate (circle: optimal technical noise parameters fit to the entire glutamatergic dataset from sample B08; triangles: optimal technical noise parameters fit to the glutamatergic subtypes; color: number of cells in the dataset; grid indicates the points evaluated during the inference process; Gaussian jitter added).

**Supplementary Figure 6:**
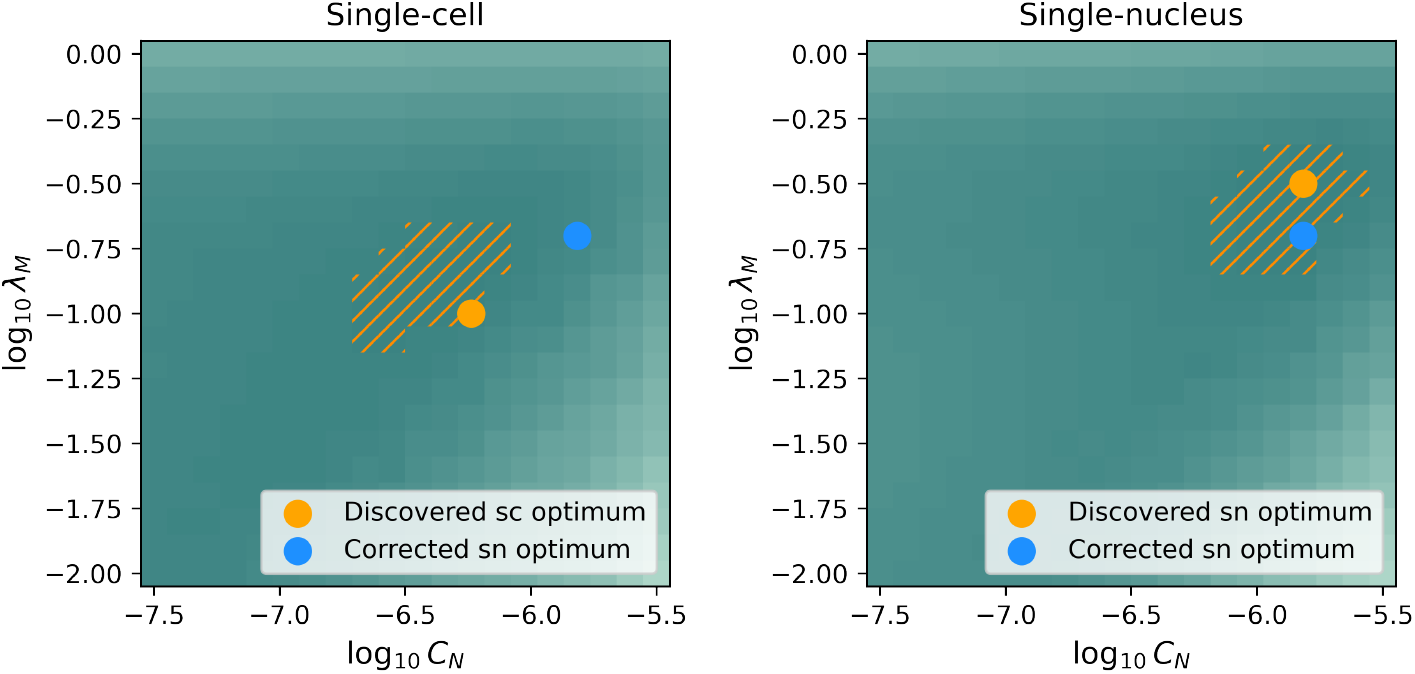
Inferred locations of technical noise parameters for a pair of single-cell and single-nucleus datasets suggest the single-nucleus dataset has higher parameter values, or more effective capture. We use a physical argument (Section 9.4.2) to propose a “corrected” single-nucleus optimum, which provides the best agreement with nascent RNA biological parameters. This optimum falls within the top 5th percentile of the technical noise parameter likelihood landscape (sc: single-cell; sn: single-nucleus; shade of teal: overall Kullback-Leibler divergence (KLD) between data and fit, darker is lower/better; orange points: optima discovered using the standard *Monod* procedure; hatched regions: top 5th percentile of the KLD landscape; blue points: the location of the single-nucleus optimum after correction).

**Supplementary Figure 7:**
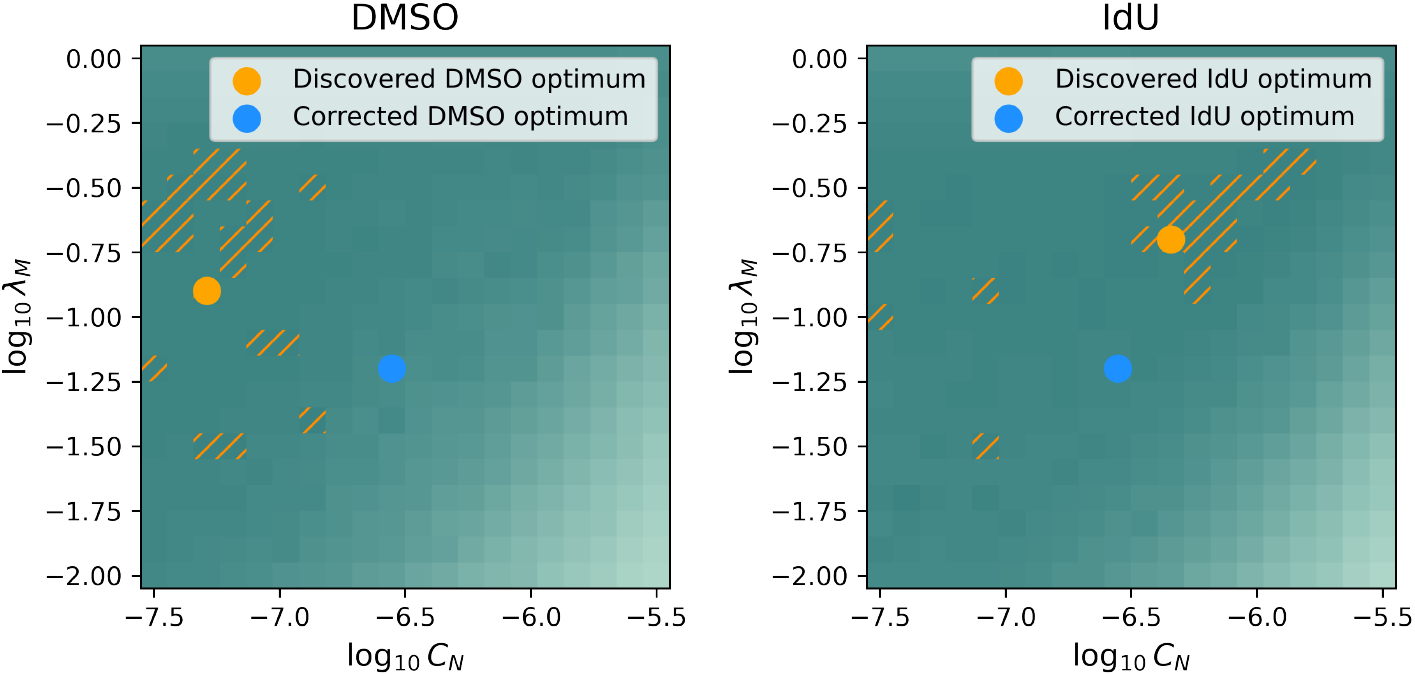
Technical noise parameter landscapes for a pair of 10x v2 single-cell datasets are rugged and poorly informative; although optima exist, they do not appear to be reliable. We tentatively propose a “corrected,” common optimum (Section 9.2.4), which is motivated by parameter values discovered by studying v2 and v3 technical replicates in a previous study [24] (shade of teal: overall Kullback-Leibler divergence (KLD) between data and fit, darker is lower/- better; orange points: optima discovered using the standard *Monod* procedure; hatched regions: top 5th percentile of the KLD landscape; blue points: the location of the optima after correction).

### S3.5 Data integration

#### S3.5.1 snRNA-seq datasets

**Supplementary Figure 8:**
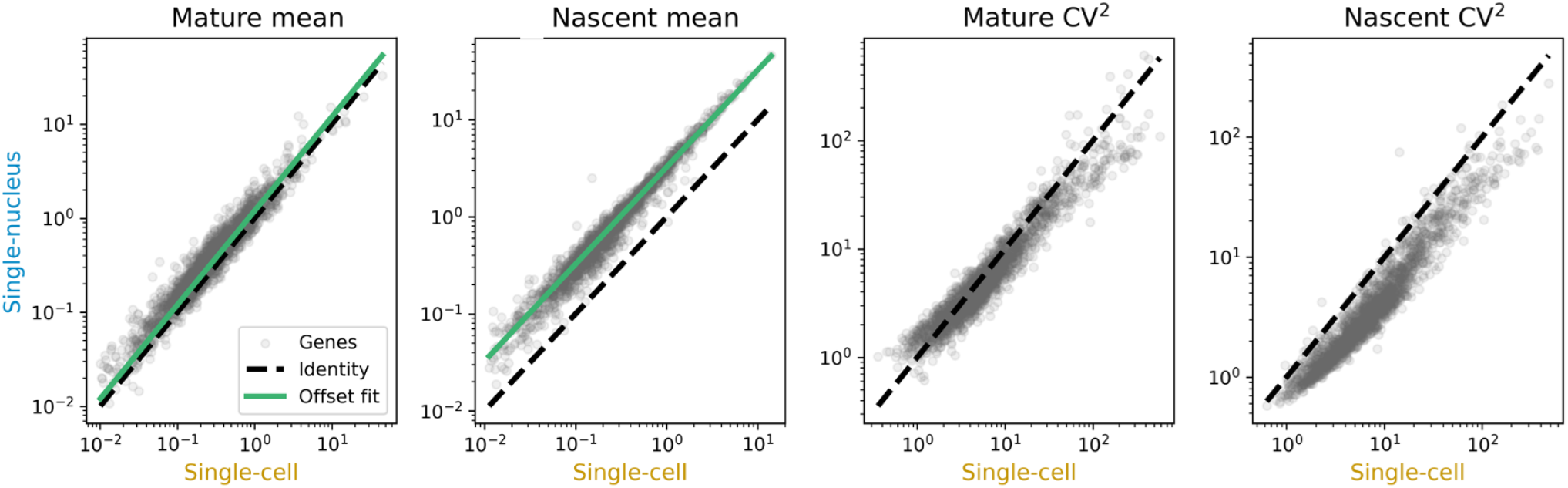
A single-nucleus RNA-seq dataset (n=3,256 cells) shows higher mature and nascent averages than a matched single-cell RNA-seq dataset (n=5,150 cells), with correspondingly lower noise (as quantified by the CV^2^).

#### S3.5.2 seqFISH datasets

**Supplementary Figure 9:**
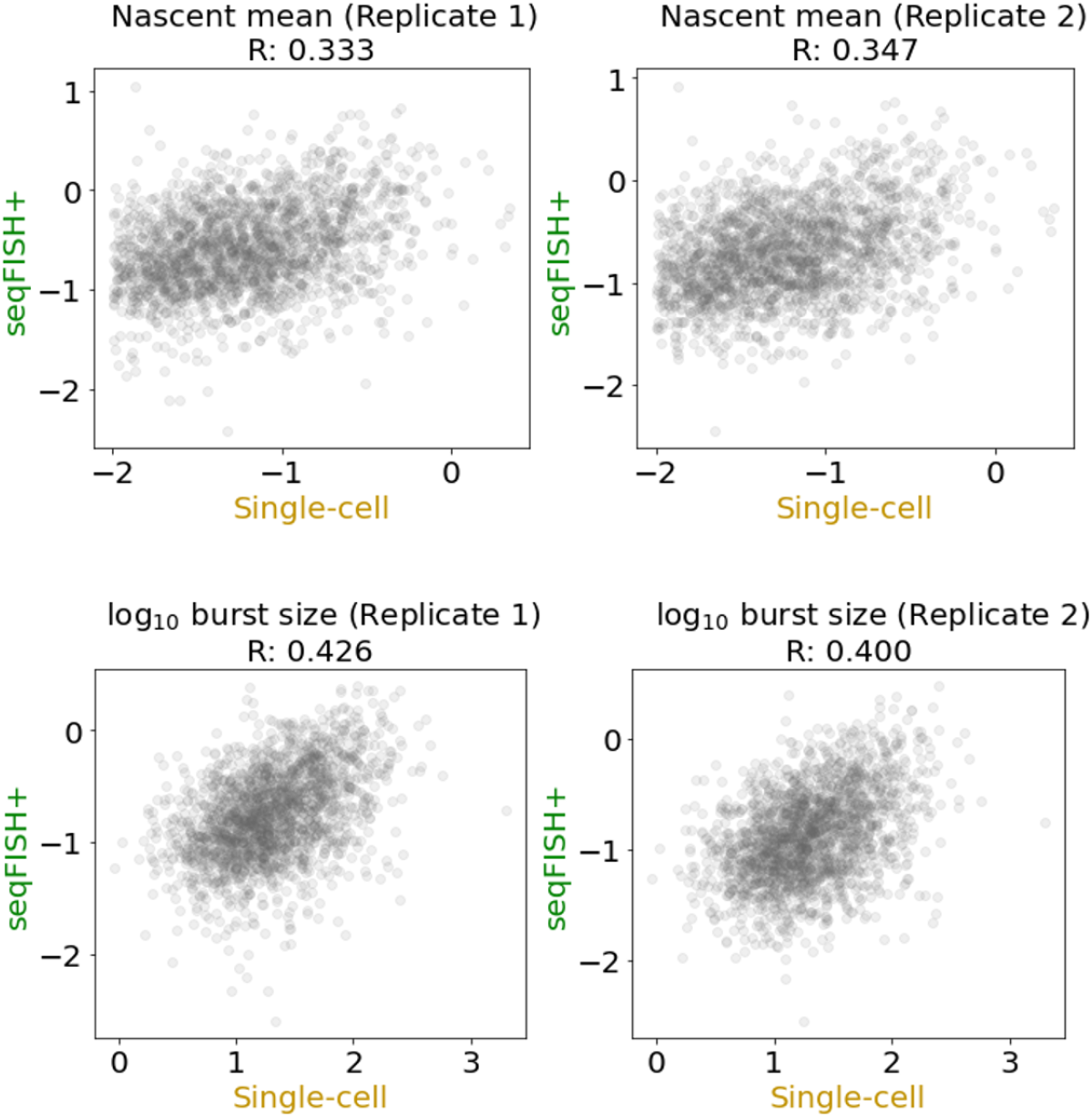
Top row: The seqFISH nascent RNA measurements do not match the scRNA-seq nascent RNA measurements (seqFISH+ means are taken over n=523 cells and n=553 cells for replicates 1 and 2, respectively; scRNA-seq means are taken over n=12,167 cells). Bottom row: The burst sizes tend to be somewhat more correlated, although their scales are divergent, possibly because the inference procedure for seqFISH does not incorporate modeling of technical noise.

### S3.6 Dimensionality reduction and normalization

#### S3.6.1 Model-free analysis of noise fractions and comparison to *Monod*

#### S3.6.2 Model-free analysis of correlations and comparison to *Monod*

**Supplementary Figure 10:**
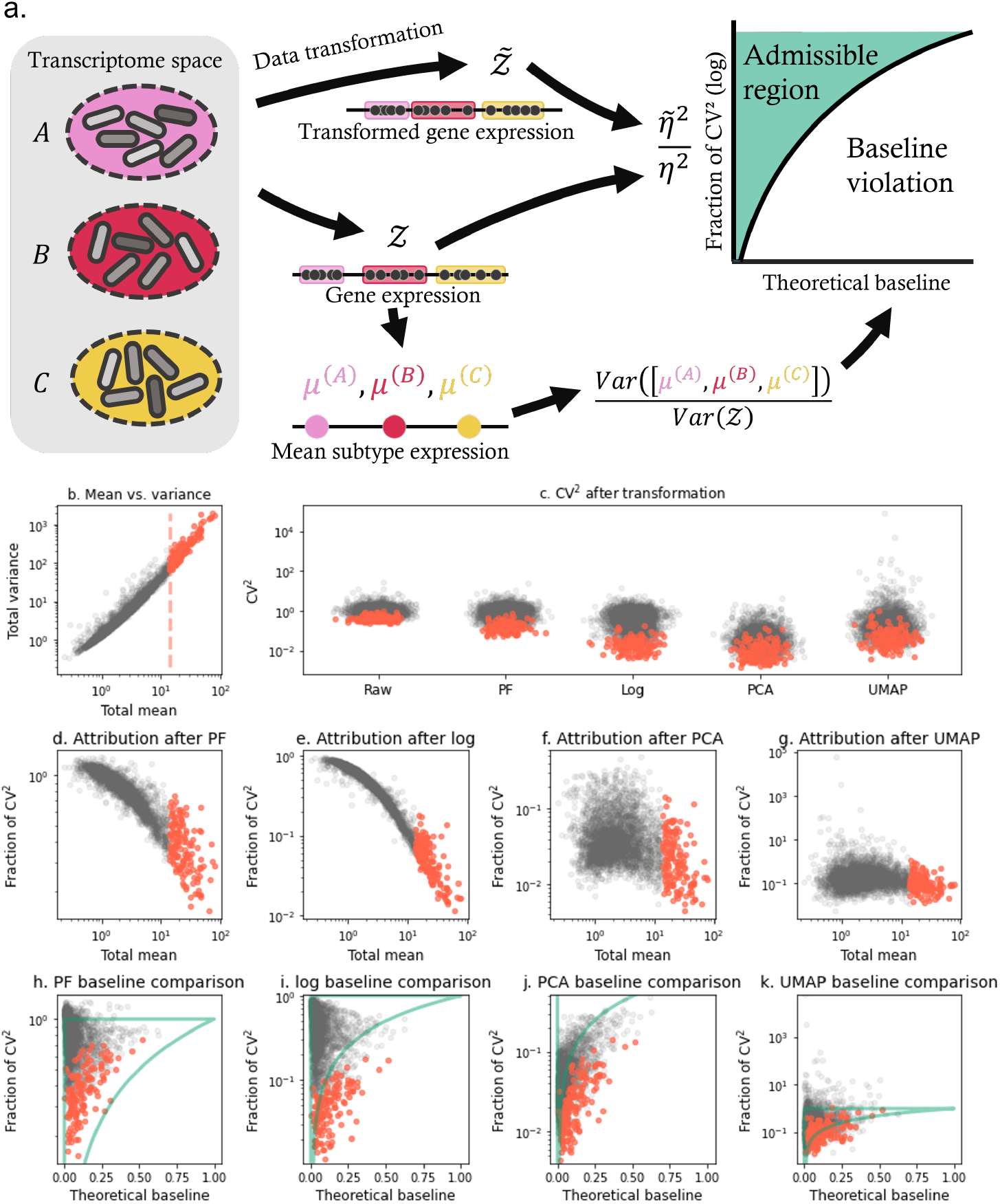
Normalization and dimensionality reduction distort and underestimate biological variation under a model-free baseline, especially in high-expression genes. **a**. A proposed baseline for the analysis of residual variation after data transformation: the fraction of biological variability can be bounded by a theoretical baseline, which is computed from the variation in average subpopulation expression. If this baseline is violated, the data transformation has discarded some biophysically meaningful variation. **b**. High-expression genes from glutamatergic neurons (n=5,343 cells) have high variance (gray points: genes below the 95th percentile by mature RNA expression; red points: genes above the 95th percentile by mean mature RNA expression, red line: percentile threshold). **c**. Proportional fitting size normalization (PF), log-transformation (log), and principal component analysis (PCA) globally deflate the squared coefficient of variation (CV^2^), whereas Uniform Manifold Approximation and Projection (UMAP) globally inflates it (gray and red points: as in **b**). **d**.-**g**. All four of the steps substantially deflate high-expression genes’ CV^2^ relative to raw data, implicitly attributing their variability to nuisance technical effects (gray and red points: as in **b**). **h**.-**k**. The deflation of variability results in the violation of the theoretical lower bound computed from cell subpopulation differences, particularly for high-expression genes (gray and red points: as in **b**; curved teal line: identity baseline, below which biological variability is removed; horizontal teal line: threshold, above which variability is inflated relative to raw data).

**Supplementary Figure 11:**
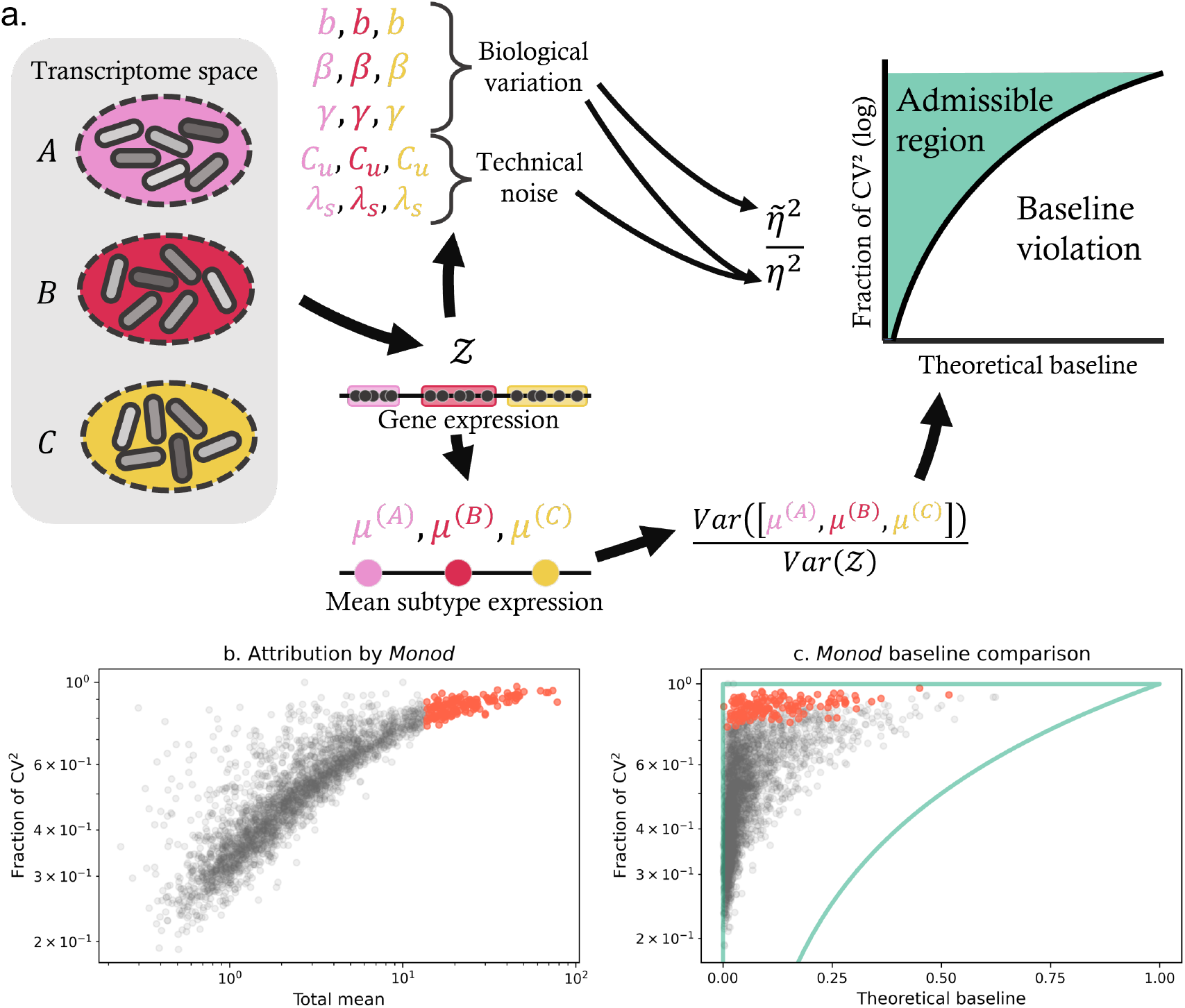
The *Monod* mechanistic analysis of biological and technical variability produces coherent results. **a**. The baseline introduced in Supplementary Figure 10a may be compared to point estimates of the biological variability fractions, which follow immediately from a fit to a parametric model of transcription and sequencing. **b**. The *Monod* fits explicitly attribute the variability in high-expression genes from glutamatergic neurons (n=5,343 cells) to biological phenomena (gray and red points: as in Supplementary Figure 10b). **c**. The *Monod* results lie entirely within the admissible region (gray and red points: as in **b**; curved teal line: identity baseline, below which inferred biological variability is lower than inter-cell population variability; horizontal teal line: threshold, above which inferred biological variability exceeds that of raw data).

**Supplementary Figure 12:**
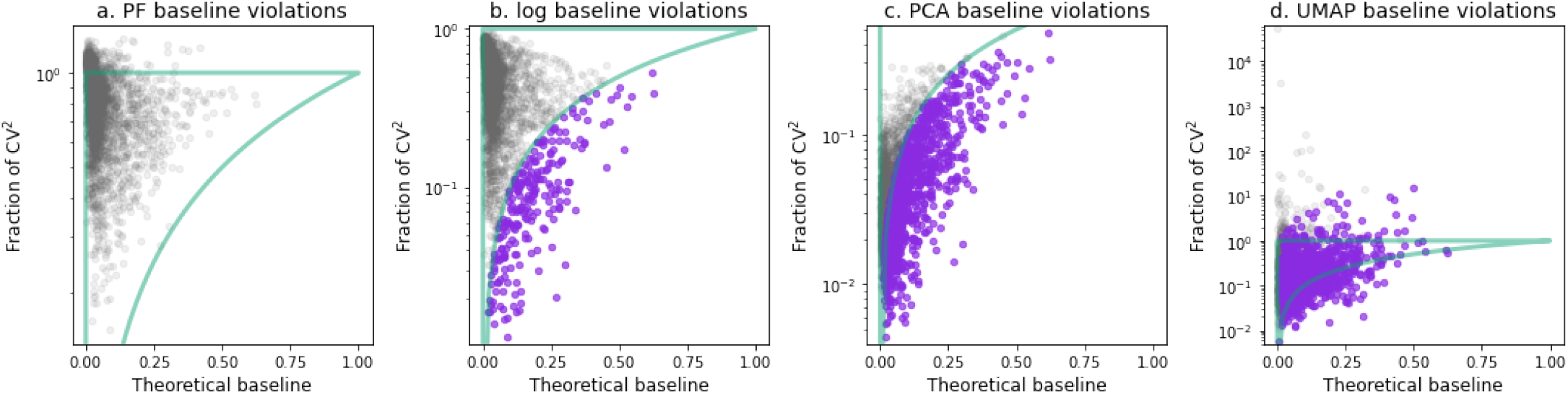
The deflation of variability results in the violation of the theoretical lower bound computed from cell subpopulation differences (across n=5,343 cells), which degrade the information available at later steps (violet points: genes that violated the theoretical baseline at the current or any previous step of data transformation; gray points: all other genes; teal lines: as in Supplementary Figure 10h-k).

**Supplementary Figure 13:**
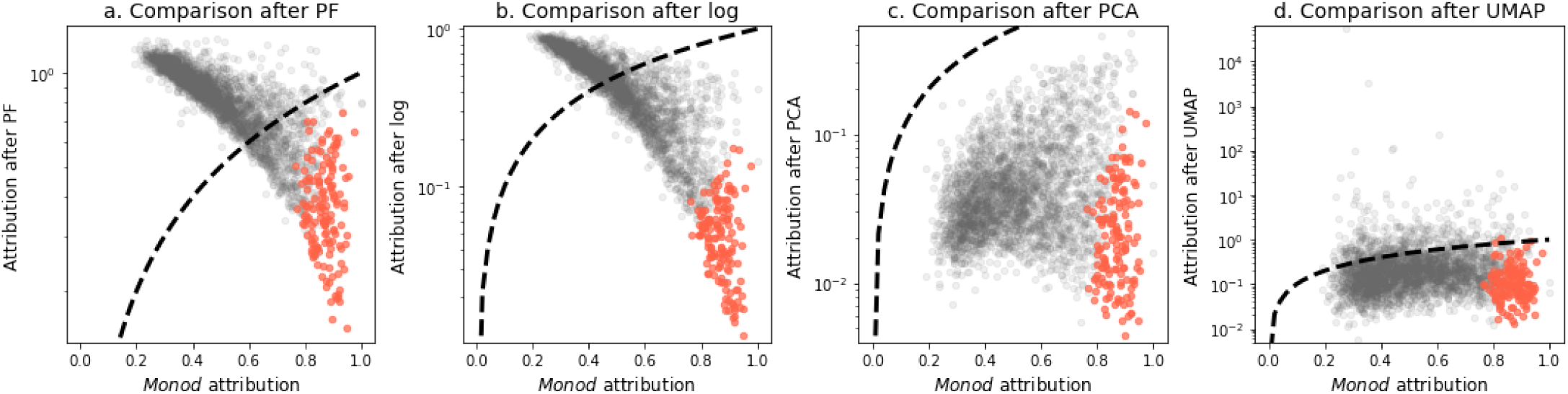
Standard single-cell RNA sequencing transformations fundamentally disagree with the *Monod* mechanistic analysis regarding the attribution of variability (calculated across n=5,343 cells), especially for high-expression genes: where the former proposes their variability is largely technical, the latter proposes it is largely biological (points: as in Supplementary Figure 10d-g; dashed black line: identity).

**Supplementary Figure 14:**
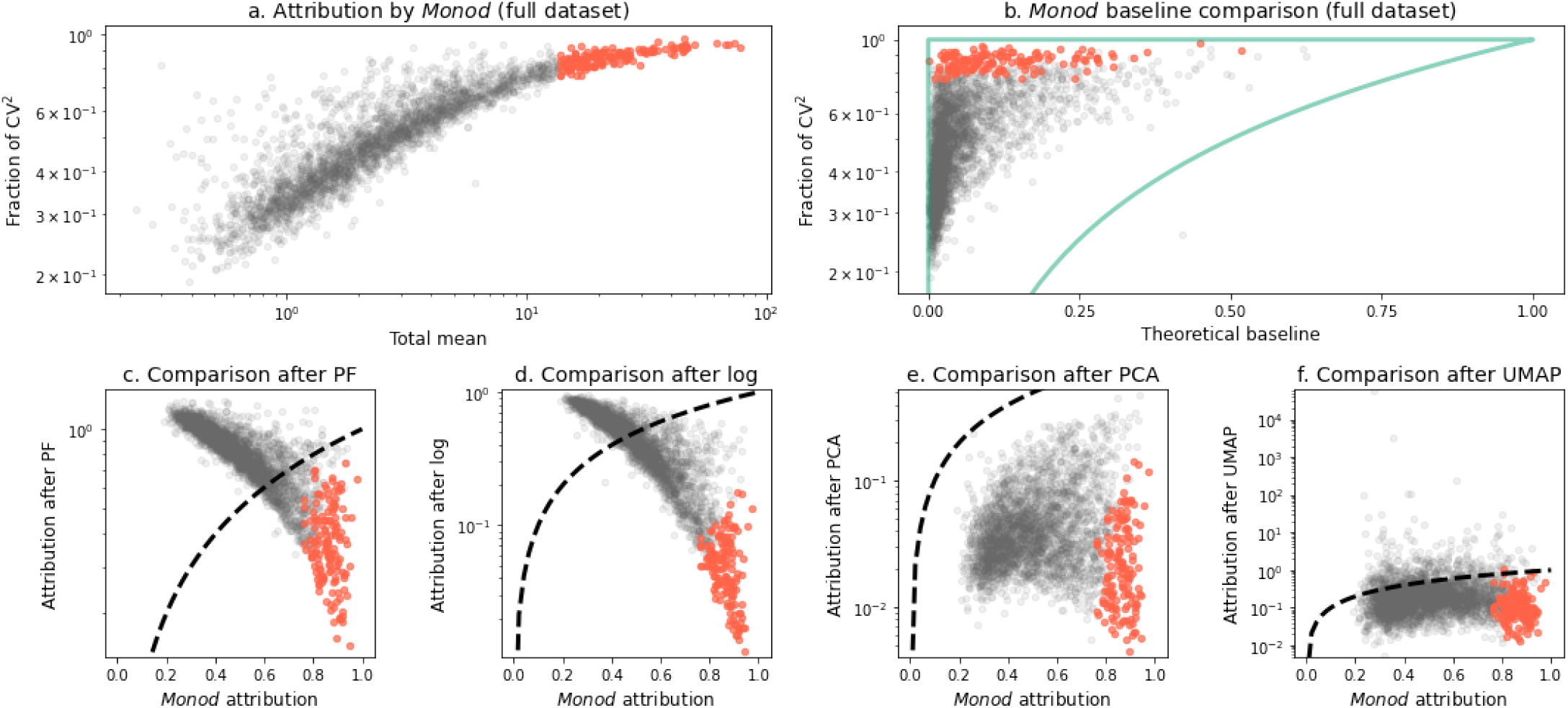
If the fits are sufficiently accurate, the procedure in Supplementary Figure 10a automatically enforces the theoretical baseline; however, the results largely hold even if the *Monod* fit is applied to the entire glutamatergic dataset, with no information about subpopulations. The striking similarities suggest that *Monod* largely attributes biological variability to transcriptional bursting, rather than cell type differences. **a**.-**b**. As in Supplementary Figure 10b-c, reproduced using a *Monod* fit to the union of glutamatergic cell subtypes (n=5,343 cells). This procedure yields consistent results and a single baseline violation. **c**.-**f**. As in Supplementary Figure 13, reproduced using a *Monod* fit to the union of glutamatergic cell subtypes. This procedure yields consistent results.

**Supplementary Figure 15:**
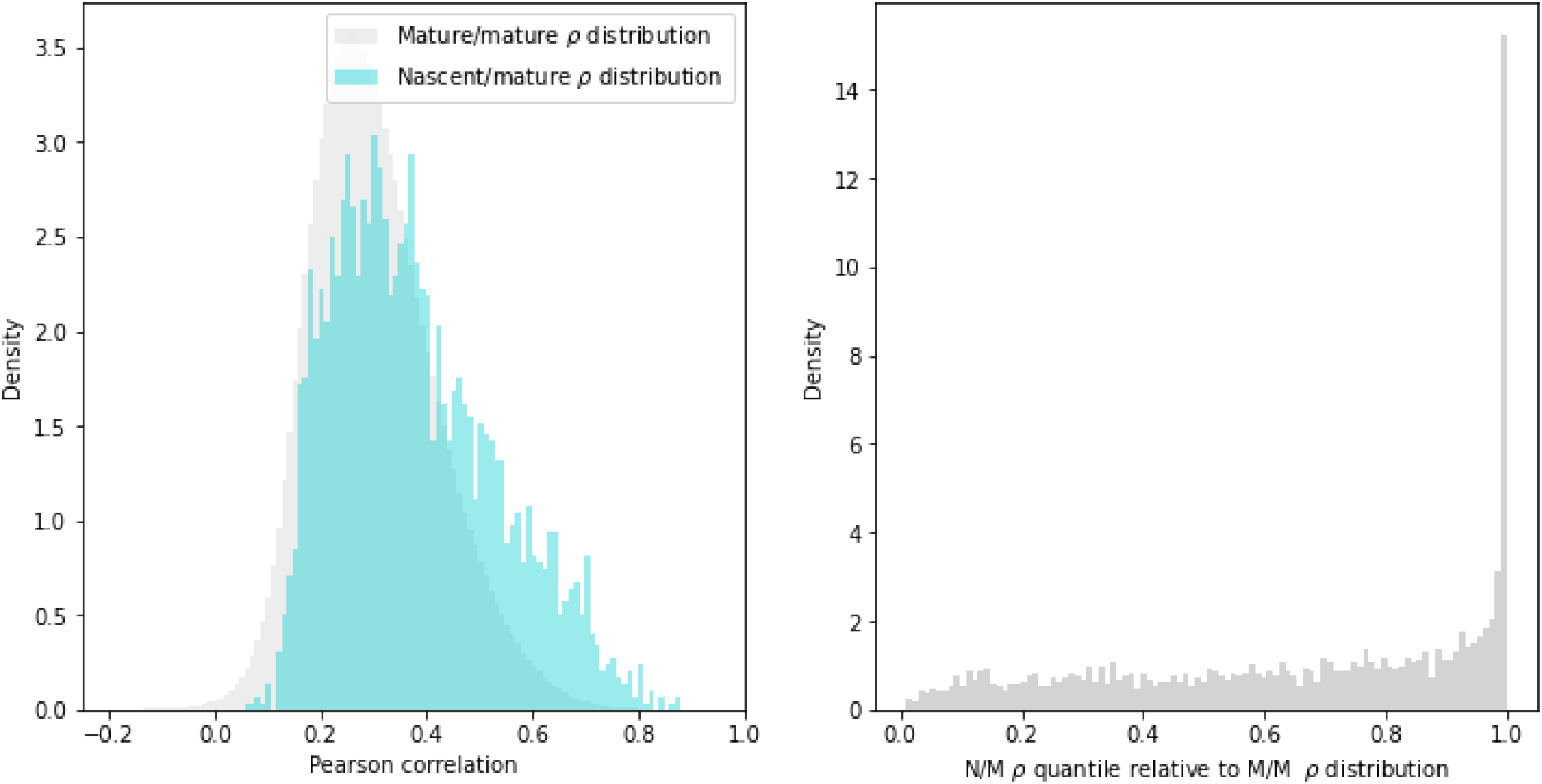
**a**. Nascent/mature correlations are high relative to the mature/mature correlation distribution (from glutamatergic neurons, n=5,343 cells). **b**. For a large fraction of genes, nascent/mature correlations are conspicuously higher relative to the mature/mature correlation distribution.

**Supplementary Figure 16:**
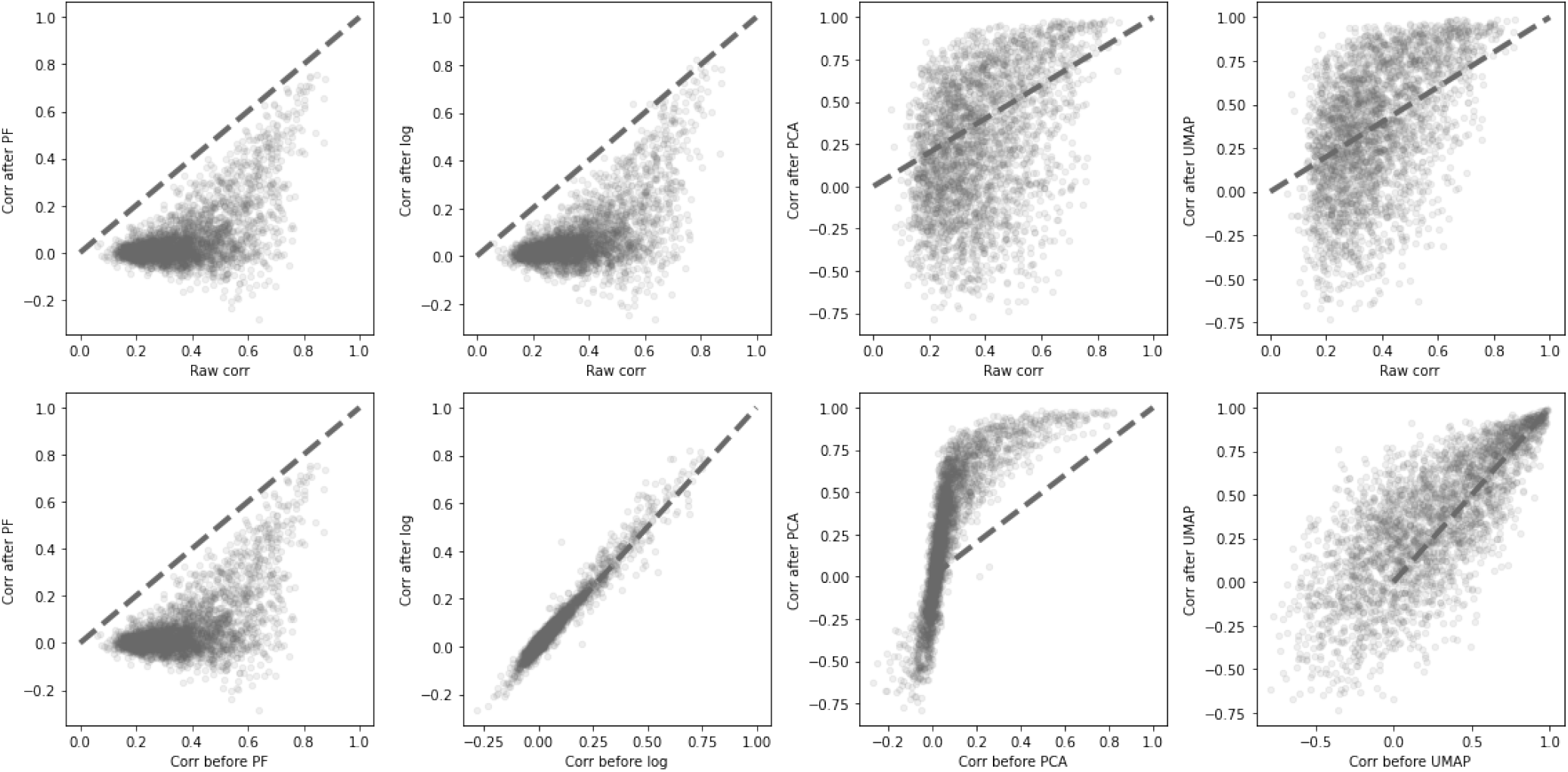
Nascent/mature correlations are substantially distorted by each successive step of data transformation (from glutamatergic neurons, n=5,343 cells), both relative to the original distribution and the immediate input of the step. This distortion often involves a change of sign.

**Supplementary Figure 17:**
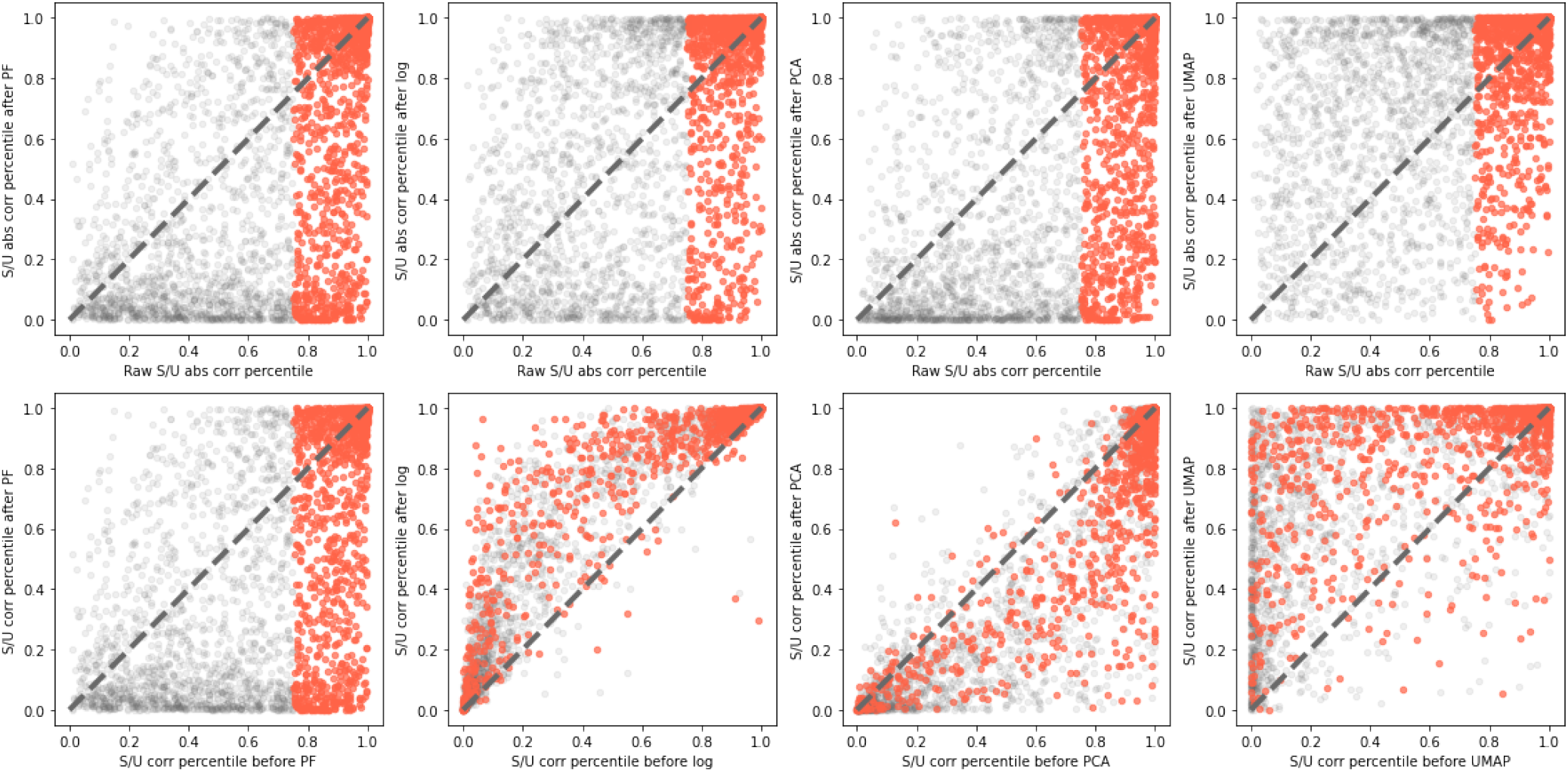
Nascent/mature correlations’ rank relative to mature/mature correlation distribution is substantially distorted by each successive step of data transformation (from glutamatergic neurons, n=5,343 cells), with a sizeable fraction of correlations in the top quartile of these ranks (orange points) being lost.

**Supplementary Figure 18:**
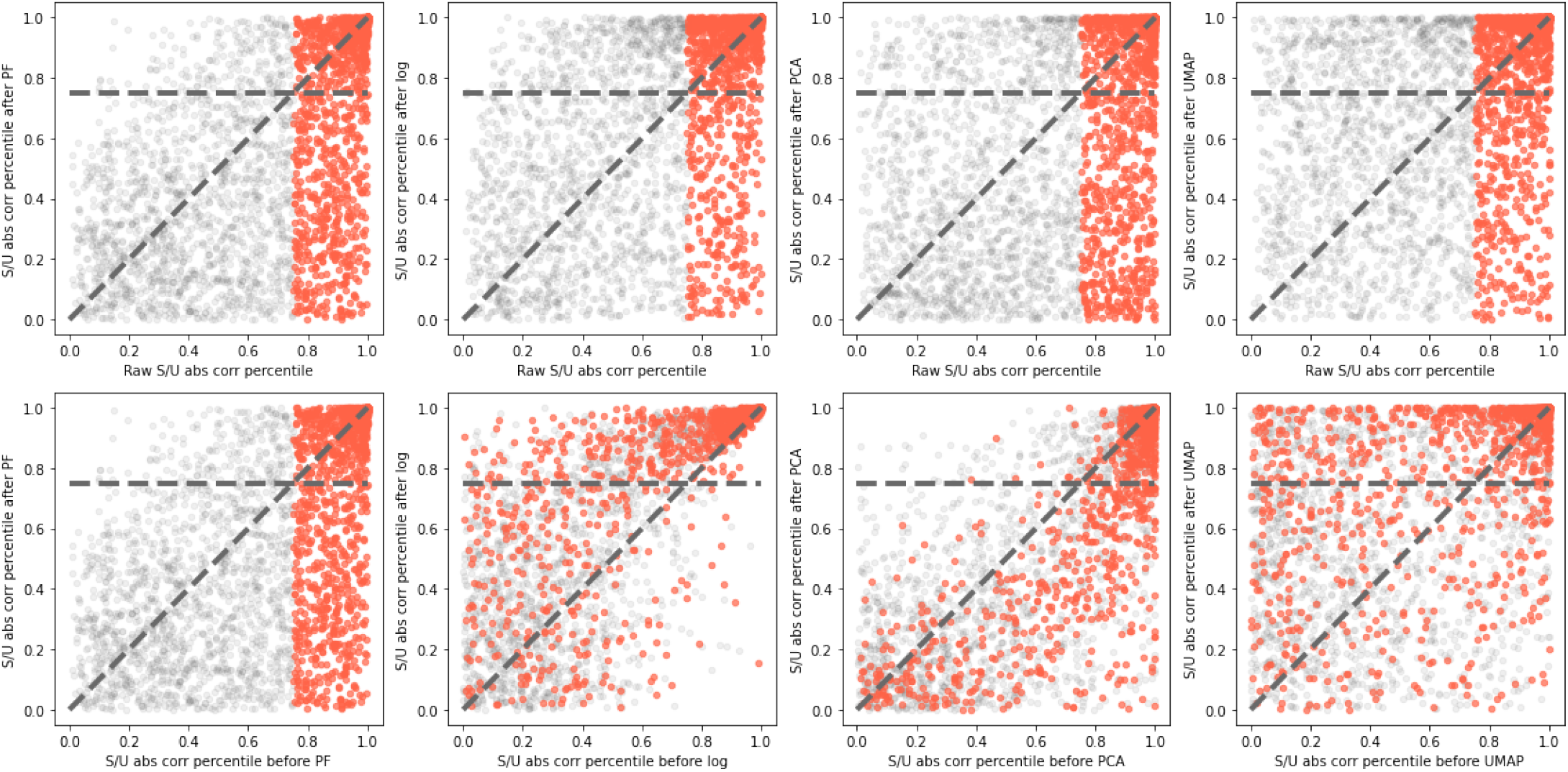
Nascent/mature absolute correlations’ rank relative to mature/mature absolute correlation distribution is substantially distorted by each successive step of data transformation (from glutamatergic neurons, n=5,343 cells), with a sizeable fraction of correlations in the top quartile of these ranks (orange points) being lost.

**Supplementary Figure 19:**
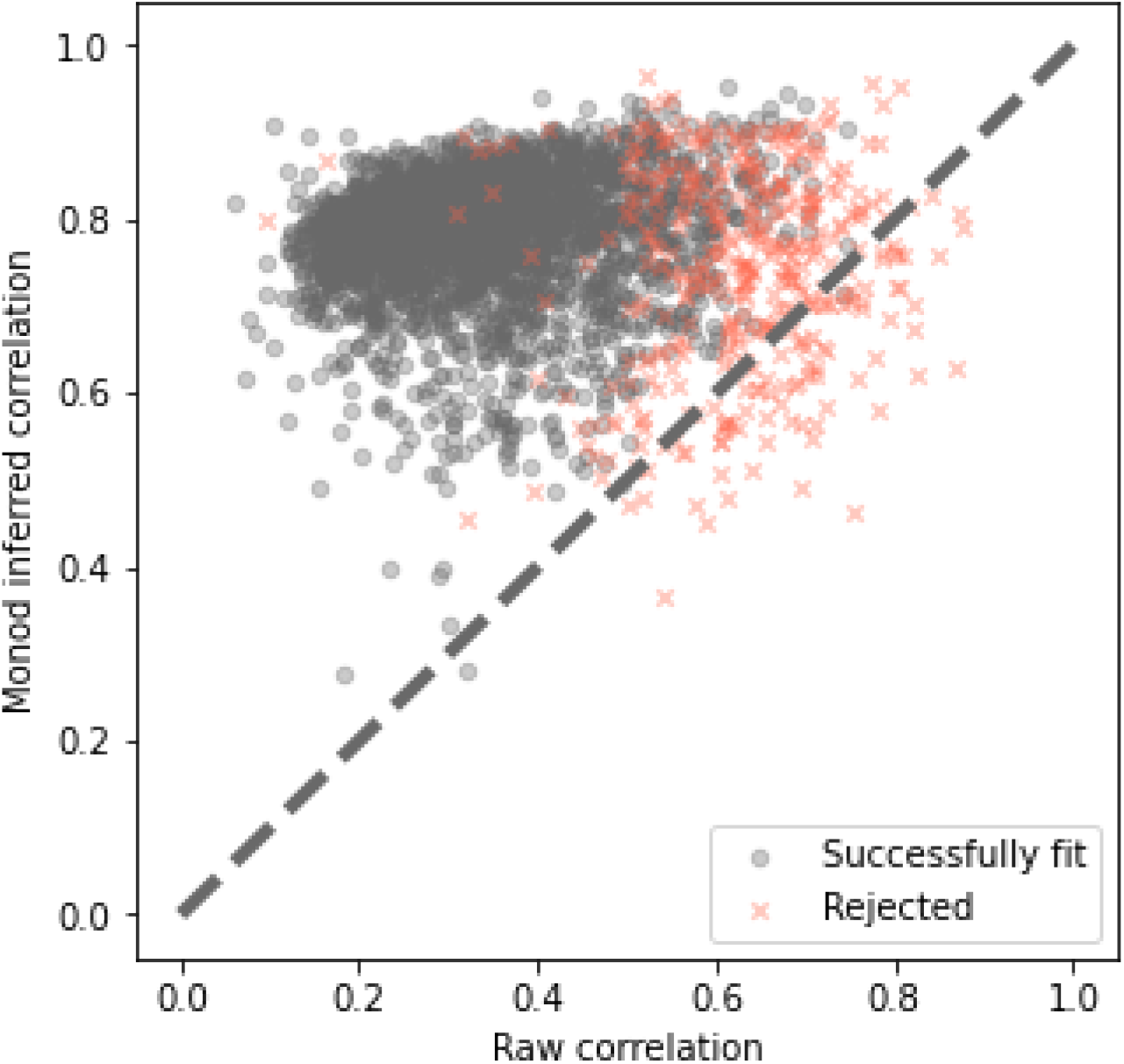
When fits are successful, the nascent/mature correlations predicted by the *Monod* model are strictly higher than raw correlations, according with intuition (from glutamatergic neurons, n=5,343 cells).

